# Predatory mites, a green pesticide, and an Entomopathogenic compound: A proposed IPM tactic based on pest species diversity indices and population dynamics

**DOI:** 10.1101/2022.02.12.480204

**Authors:** Islam Mohammad Zidan, Elsayed Mohamed Ahmed K. El-Saiedy, Gomaa Mohamed Abou-Elella, Mourad Fahmy Hassan

## Abstract

The study was aimed to conduct the population dynamics and diversity indices for three major pest species in order to design an IPM protocol in two experimental sites (Om Saber, Beheira Governorate 30°29’50.6”N, 30°46’18.8”E), and (Kom Oshim, Fayoum Governorate 29°34’40.9”N, 30°55’38.3”E). The resulted data showed statistically significant fluctuation, population dynamics, abundance, distribution, and diversity indices of the two-spotted spider mite (TSSM) *Tetranychus urticae* Koch (Acari: Tetranychidae), the silver leaf whitefly *Bemisia tabaci* Genn. (Hemiptera: Aleyrodidae), and the onion thrips *Thrips tabaci* Lindman (Thysanoptera: Thripidae) which recorded on four plant species belonging to Brassicaceae (Siberian (Russian) kale *Brassica napus var. pabularia* L. and Italian (Tuscan) kale *Brassica oleracea var. palmifolia*), and Lamiaceae (Spearmint *Mentha spicata* L. and Saudi Mint *Mentha longifolia* L.). The proposed IPM program consisted of predatory mites; *Phytoseiulus persimilis* Athias-Henriot, *Amblyseius swirskii* Athias-Henriot, and *Cydnoseius negevi* (Swirski & Amitai) (Acari: Phytoseiidae), a green pesticide, and an entomopathogenic compound. It was concluded that abiotic and biotic factors together help in explaining why various pest species build their communities rapidly and increase their parameters that become above the EIL. Such factors are hypothesized to affect the plant-arthropod, predator-herbivore, predator-predator, and tri-trophic interactions. And it recommends the application of such protocol should consider the timing of tacking an action and merging tactics together to get the maximum efficiency.

## INTRODUCTION

Different kinds of relations between plants and various taxa of arthropods in the Agro-ecosystem, whether plant taxa are, either economic crop, or spontaneous plant. Plant-herbivore interaction is one of the possible forms. It depends on diversified chemical and morphological relations (Frago *et al*., 2022). Another relation is the Predator-herbivore relation, which is considered a second level of trophic interactions occurring in an ecosystem (Alba *et al*., 2012; Gardarin *et al*., 2018). Connecting these interactions together forms a tri-trophic relation, that consisted of Plant-Herbivore-Predator interaction (Verkerk, 2004; Kavitha and Reddy, 2014), and this one we do hypothetically suggest, is the main method of the biological control applications and Integrated Pest Management (IPM) tactics which used to supress pest infestations.

IPM is a complex of agricultural, chemical, biological, and ecological procedures, as well as, good knowledge about biodiversity and distribution of living organisms within the habitat (El-Shafie, 2019). The most IPM strategy’s goal is to prevent pests from reaching economically damaging levels without causing a risk to the environment (Koul *et al*., 2004; Smagghe and Diaz, 2012). Successful IPM program may has some components such; monitoring crops for pests, pest identifying accuracy, detecting economic thresholds, implementing integrated pest control tactics, and evaluation. The factors that render crop habitats unsuitable for pests and diseases include limitation of resources, competition, parasitism, and predation (Drinkwater *et al*., 1995; Edwards-Johnes, 2001; Ehi-Eromosele *et al*., 2013).

The first use of biological control backs to the ancient Chinese at the 3^rd^ - 4^th^ centuries A.D. (Shijiang, 1983). Till now, very large number of predatory and/or parasite taxa been used to control pest infestations that caused massive economic losses (Smagghe and Diaz, 2012). Predatory phytoseiid mites have reported to be successful control means for spider mites (Negm *et al*., 2014; Alatawi *et al*., 2018), eriophyids (Momen *et al*., 2004, 2014; Momen and Abdel-Khalek, 2008; Abou-Elella *et al*., 2014; Melo *et al*., 2015; Abdel-Khalek and Momen, 2022; Ferreira *et al*., 2022), whiteflies (Teich, 1966; Nomikou *et al*., 2001, 2002, 2003), thrips (Messelink *et al*., 2006; Arthurs *et al*., 2009; Sanad and Hassan, 2019), aphids (Messelink *et al*., 2013), and they were proven to fed alternative sources as pollen (van Rijn and Tanigoshi, 1999; Abou-Elella *et al*., 2014; Delisle *et al*., 2015 b; Rahmani *et al*., 2021; Xin and Zhang, 2021), fungi (Zemek and Prenerová, 1997; Momen and Abdelkhader, 2010), or other factitious food/artificial diet to be mass produced (Janssen and Sabelis, 2015; Delisle *et al*., 2015 a, b; Momen *et al*., 2020; Xin and Zhang, 2021).

New trends of bio-agents which been used are the Entomopathogenic fungal species, e.g., *Beauveria bassiana* (Bals.) Vuill. and *Metarhizium anisopliae* (Metschn.) Sorokin to control insect and mite pests (Akmal *et al*., 2013; Wraight *et al*., 2016). These fungi can inhabit the plant/crop ecosystem; on leaves (Garrido-Jurado *et al*., 2015), soil (Evans, 1982), or endophytically (Greenfield *et al*., 2016). These fungi were highly recommended on IPM protocols due to their wide distribution and diversity of hosts in different localities and conditions (Lacey *et al*., 2015; McGuire and Northfield, 2020).

Another new trend of IPM is using the eco-friendly or green-pesticides, which consisting mainly of natural resources, such plant essential oils and extracts (Isman, 2006; Isman and Machial, 2006; Koul *et al*., 2008). Plant essential oils were reported to supress pest infestations on different climatic zones (Regnault-Roger *et al*., 1993; de Melo *et al*., 2019; Allam *et al*., 2020; Ebadollahi *et al*., 2020).

Some studies have proposed using the mixture of phytosiieds, entomopathogenic, and biopesticides to control different pest infestations (El-Saiedy *et al*., 2008, 2015; Abou-Awad *et al*., 2017; El-Saiedy and Fahim, 2021). On other side, the climate change factors are affecting different kinds of agricultural pests and beneficial arthropods (Shrestha *et al*., 2019; Skendžić *et al*., 2021), these changes affected the biological aspects, prey preferences, distribution, population dynamics, and threaten the IPM applications (Halsch *et al*., 2021: Karthik *et al*., 2021). Therefore, this study was aimed to conduct the population dynamics and diversity indices for three major pest species infested four Brassicaceae and Lamiaceae plants, in order to design a successful IPM protocol.

## MATERIAL AND METHODS

### 1. Experimental sites

Two locations were selected to perform the field experiments; Om Saber, Kom Hamada, El Beheira Governorate (30°29’50.6”N 30°46’18.8”E), and Kom Oshim, Fayoum Governorate (29°34’40.9”N 30°55’38.3”E) (Fig. 1). The data of population dynamics were recorded for two seasons; Mar to Aug 2016, and Feb to Jul 2017. The proposed IPM procedures were taken place form Apr to Aug 2017, and replicated from Feb to Jun 2018.

**Fig. 1.**
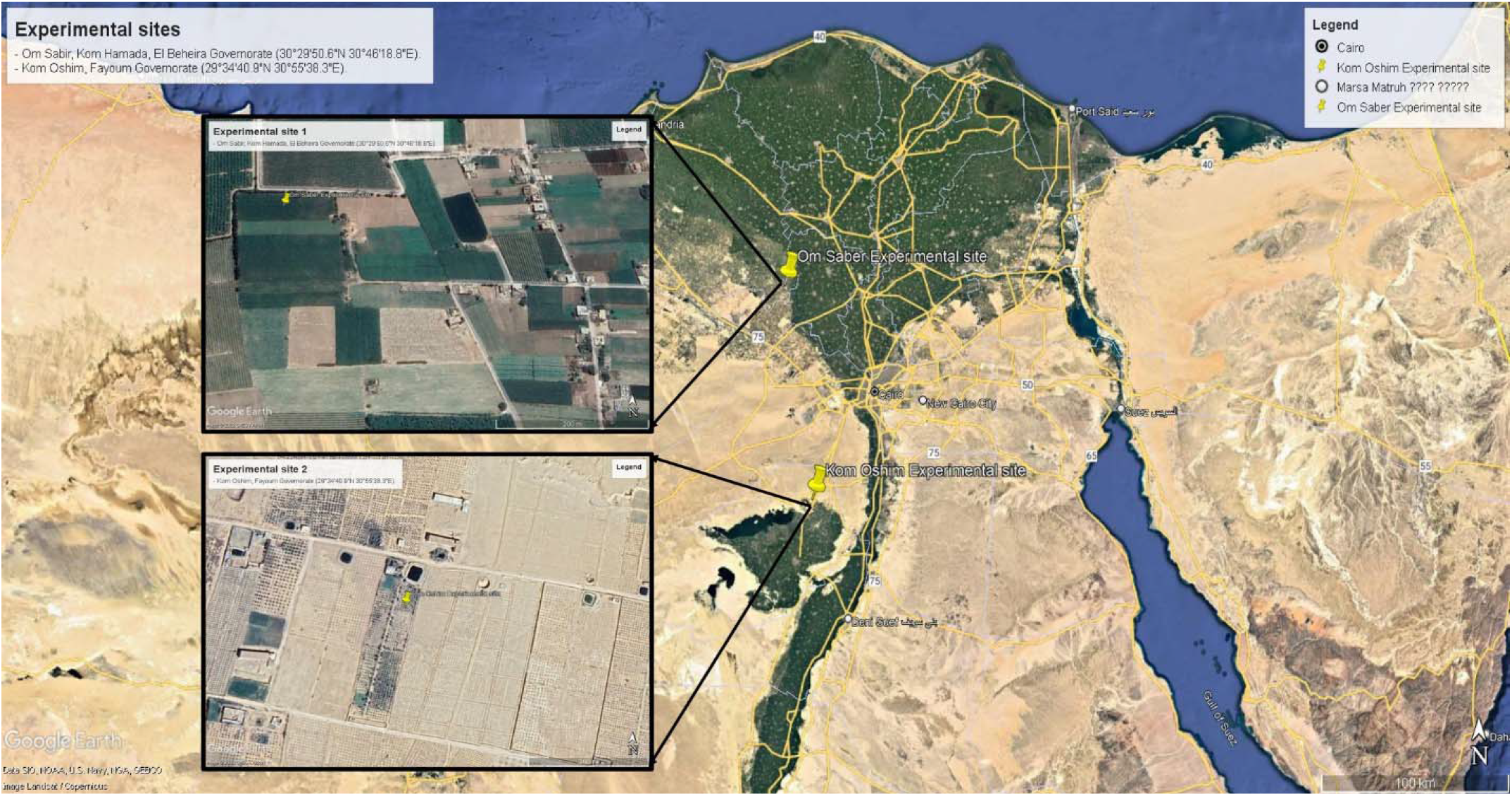
Google Earth map photography of the experimental locations (pointed with pin), i) Om Sabir, Kom Hamada, El Beheira Governorate (30°29’50.6”N 30°46’18.8”E), and ii) Kom Oshim, Fayoum Governorate (29°34’40.9”N 30°55’38.3”E).

### 2. Plant sources

Four plant species belonging to two families; Brassicaceae, Siberian (Russian) kale *Brassica napus var. pabularia* L., Italian (Tuscan) kale *Brassica oleracea var. palmifolia*, and Lamiaceae, Spearmint *Mentha spicata* L. and Saudi Mint *Mentha longifolia* L. Brassicaceae plants were introduced from Egyptian Hydrofarms farm (KM 53 Cairo-Alex Desert Road, inside Al-Azzazy Village, 30°09’07.0”N 30°51’00.2”E). While Lamiaceae plants were introduced from a private farm of herbal and medicinal plants in Fayoum Governorate (Ibshway, 29°21’07.4”N 30°44’17.8”E).

### 3. Planting

The design of the experiment is showed in Fig. 2. Mixture of compost and organic manure added in the soil before planting. Seedlings transferred into fertilized soil and covered with sand layer, then irrigated. Also, in each plot, there were five rows; for treatments. About 0.50 m between plant plots (in each treatment), and 1 m between treatment plots to easily handling, maintaining, recording, sampling, and to prevent predatory species movement to the other plots.

**Fig. 2.**
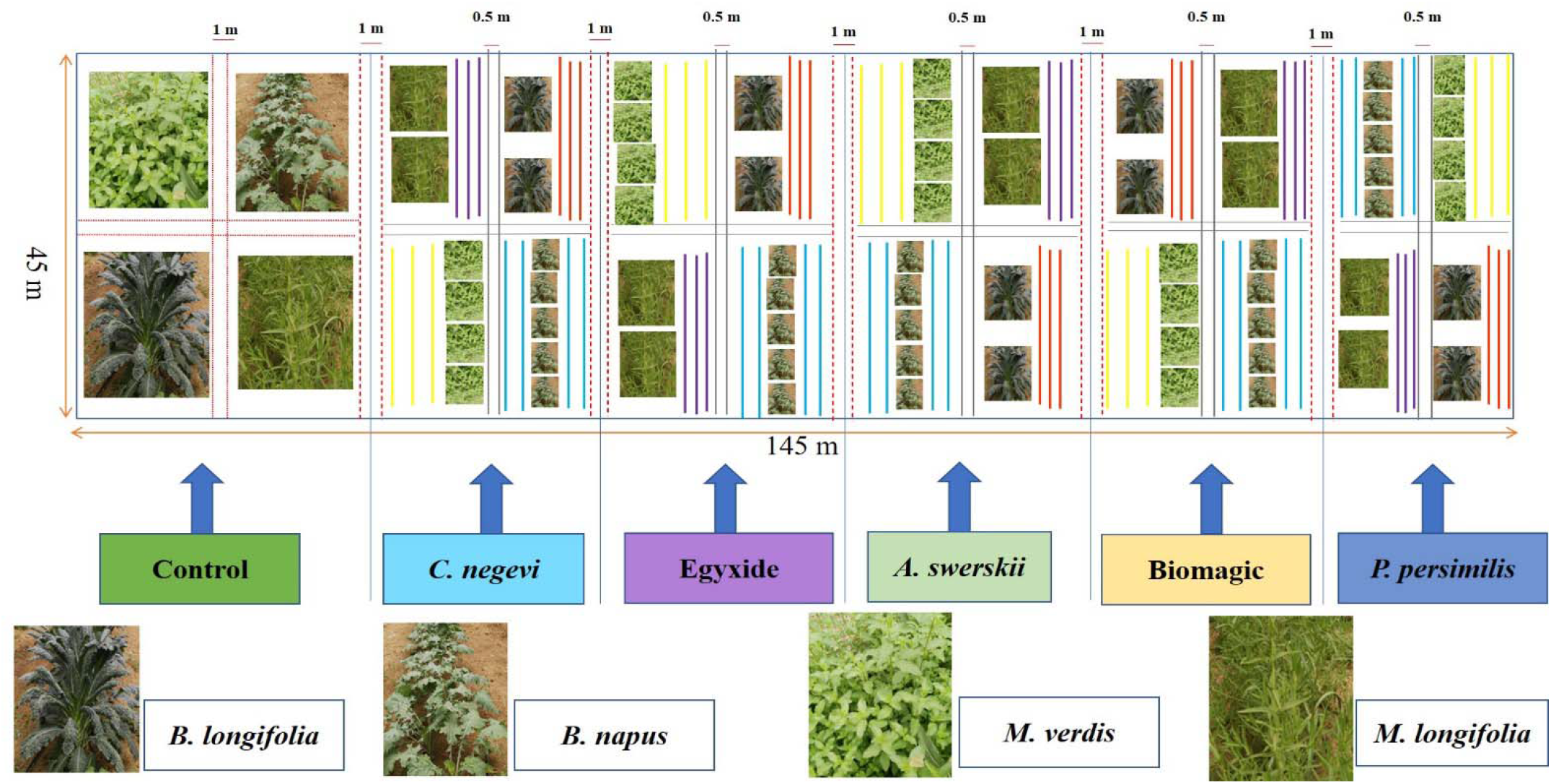
Schematic diagram of the experiment’s plantation

### 4. Sampling

Samples collected for the pest population dynamics weekly from each site, and were investigated in the Acarology lab, NRC. Active stages were counted and recorded. Besides, weeds that checked for arthropod occurrence to figure out the possible interactions. Ten leaves of each plant were checked for pest dynamics randomly. While, another ten leaves were checked randomly to record pest population before/after each treatment when control application been taken place.

### 5. Used control strategies

#### 5.1. Predatory mite species

Three phytoseiid species, *Phytoseiulus persimilis* Athias-Henriot, *Amblyseius swirskii* Athias-Henriot, and *Cydnoseius negevi* (Swirski & Amitai), were introduced as bio-agents to control pests existed in the experimental locations. Predatory mites were commercially available from private companies in Qaha, Al-Qalioubia Governorate, Badr City, Al Beheira Governorate, and Al-Ayat, Giza Governorate. A single mite package was containing about 1000 individuals. Releasing ratio was 1:5 predator to pest, based on sampled pest density in preliminary investigations.

#### 5.2. Plant extracts

Egyxide^®^ is a commercial compound recommended in organic farming systems protocols and clean farming in open and semi-field conditions. Egyxide is a water-soluble natural oils mixture, that used for mite and insect pests, also to prevent plant diseases. It is containing natural emulsified plant oils 10% and Glue 2%. Recommended as foliar application with dosage 5 ml/Liter, and was available at Royal for Agricultural Development, Cairo, Egypt. http://royalagri.com.

#### 5.3. Pest pathogens

Bio-Magic^®^ is a biological insecticide based on a selective strain of naturally-occurring entomopathogenic fungus *M. anisopliae*. It was available in liquid package contains spores and mycelial fragments (1×109 CFU’s/ml) that used as foliar spray 6 ml/Liter, and was available at Gaara Establishment for Import and Export, Cairo (www.gaara.com.eg).

### 6. Control experimental procedure

Pest infestations were recorded weekly. Ten leaves of each plant (of both locations) were sampled randomly, to check the pest population density/leaf/plant to detect the Economic injury level (EIL) and the Economic Threshold Level (ETL). When population reached the EIL we started applying the proposed strategy. Predators releasing ratio was 1:5 depending on pest density/plant. *Phytoseiulus persimilis* was introduced for TSSM infestations; while *A. swirskii* and *C. negevi* were planned for controlling the TSSM, whiteflies, and thrips. Pest population before and after treatments were counted and calculated due to Henderson and Tilton (1955) module as follow: Reduction 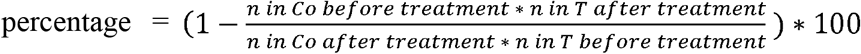 Where *n* is the pest population, *T* treated, *Co* control.

### 7. Statistical analysis

Hypothesis was tested by the Kruskal-Wallis test using SPSS computer program ver. 20.0. The null hypothesis *H_0_* suggested that distribution of pests would be the same across the host plants and locations during the experimental time. While the alternative hypothesis *H_1_* was designed as there were significant differences among pest populations in both tested plants and experimental locations. The test has conducted to determine either *H_0_* or *H_1_* is accepted. The test results rejected the *H_0_* due to significant differences within the three major pest populations recorded; *T. urticae, B. tabaci*, and *T. tabaci* (Suppl. 2; Figs. 9-12).

Results among two locations tested by Student’s test (T-test) using SPSS v. 20.0. Differences in the mean number of species before/after treatments were analysed by one-way analysis of variance ANOVA and were tested with Tukey’s test at 95% confidence level using SPSS v. 20.0. Biodiversity indices were; Shannon-Wiener index (H’), Simpson’s index (dominance D and species richness 1/D), and the similarity index between locations (Jaccard’s index), which calculated using the BioDiversity Pro. ver. 2.0 software (McAleece *et al*., 1997) and PAST ver. 4.08 software (Hammer *et al*., 2001).

## RESULTS

### 1. Population dynamics and diversity indices of three major pest species

#### a. Om Saber location

The population dynamics of *T. urticae, B. tabaci* and *T. tabaci* were detected for the two seasons, 2016 and 2017. The highest mean population of *T. urticae* was found on *B. napus var pabularia* (63.22 ± 6.10 immatures/10 leaves, and 57.11 ± 5.68 adults/ 10 leaves, *F*= 20.762, *P*= 0.000); while the lowest mean recorded in case of *M. spicata* (3.34 ± 0.21 immatures/10 leaves and 2.78 ± 0.18 immatures/10 leaves; *F*= 19.289, *P*= 0.000). *Thrips tabaci* was the lowest population that been recorded in both Lamiaceae plants. Similar results were detected on the second season, 2017, when all pest populations have recorded the highest levels of occurrence on *B. napus var pabularia* (*F*= 22.688, *P*= 0.001), and the lowest was recorded in case of *M. spicata* (*F*= 3.117, *P*= 0.001), and *B. tabaci* has the highest population recorded on the Brassicaceae in both seasons (Table 1).

**Table 1.**
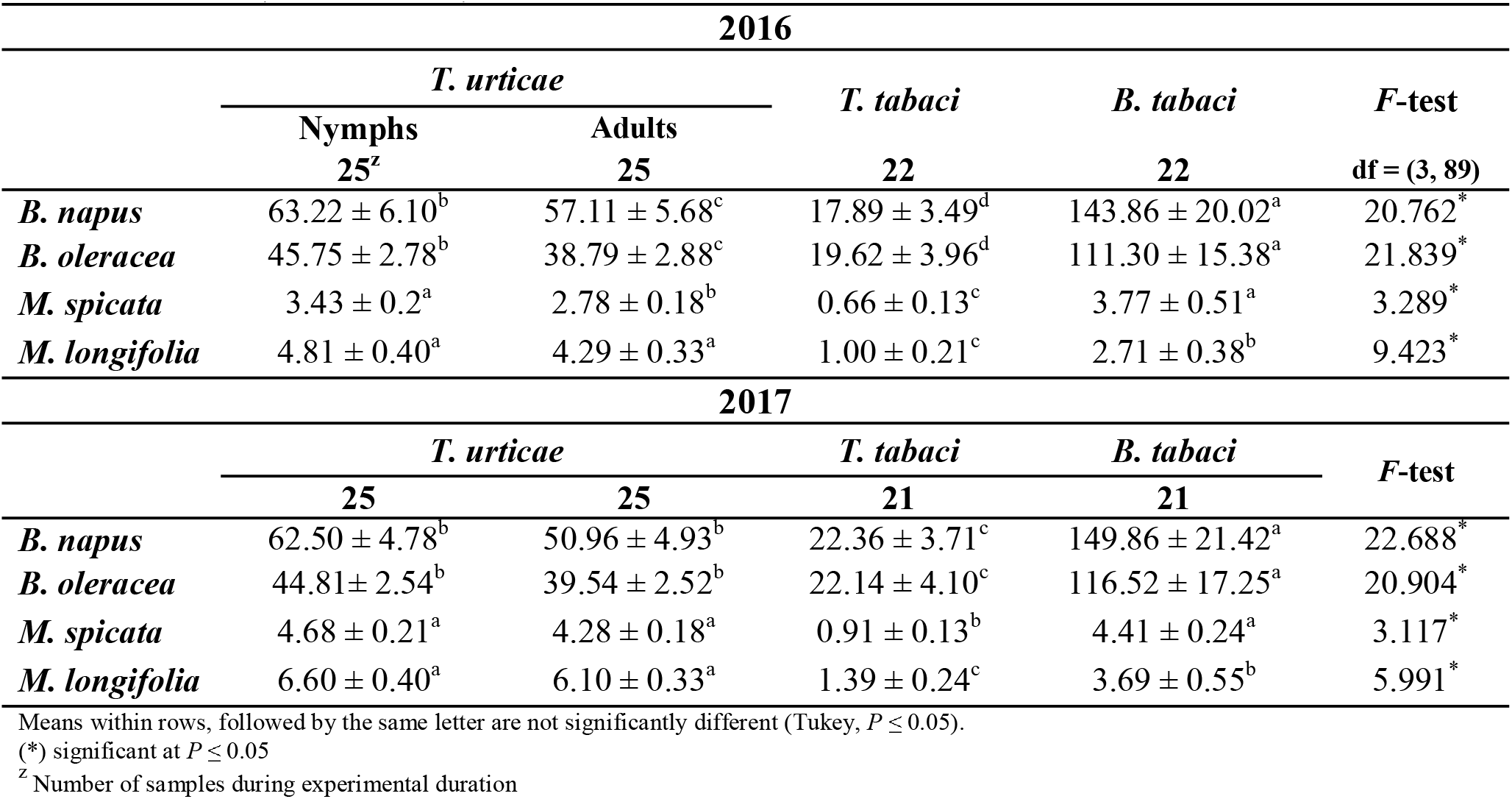
Pest species populations on four medicinal plant species (mean/10 leaves ± SE) in Om Saber location (2016 & 2017)

| 2016 |  |  |  |  |  |
| --- | --- | --- | --- | --- | --- |
|  | *T. urticae* |  | *T. tabaci* | *B. tabaci* | *F*-test |
|  | Nymphs | Adults |  |  |  |
|  | 25<sup>z</sup> | 25 | 22 | 22 | df = (3, 89) |
| *B. napus* | 63.22 ± 6.10<sup>b</sup> | 57.11 ± 5.68<sup>c</sup> | 17.89 ± 3.49<sup>d</sup> | 143.86 ± 20.02<sup>a</sup> | 20.762<sup>*</sup> |
| *B. oleracea* | 45.75 ± 2.78<sup>b</sup> | 38.79 ± 2.88<sup>c</sup> | 19.62 ± 3.96<sup>d</sup> | 111.30 ± 15.38<sup>a</sup> | 21.839<sup>*</sup> |
| *M. spicata* | 3.43 ± 0.2<sup>a</sup> | 2.78 ± 0.18<sup>b</sup> | 0.66 ± 0.13<sup>c</sup> | 3.77 ± 0.51<sup>a</sup> | 3.289<sup>*</sup> |
| *M. longifolia* | 4.81 ± 0.40<sup>a</sup> | 4.29 ± 0.33<sup>a</sup> | 1.00 ± 0.21<sup>c</sup> | 2.71 ± 0.38<sup>b</sup> | 9.423<sup>*</sup> |
| 2017 |  |  |  |  |  |
|  | *T. urticae* |  | *T. tabaci* | *B. tabaci* | *F*-test |
|  | 25 | 25 | 21 | 21 |  |
| *B. napus* | 62.50 ± 4.78<sup>b</sup> | 50.96 ± 4.93<sup>b</sup> | 22.36 ± 3.71<sup>c</sup> | 149.86 ± 21.42<sup>a</sup> | 22.688<sup>*</sup> |
| *B. oleracea* | 44.81 ± 2.54<sup>b</sup> | 39.54 ± 2.52<sup>b</sup> | 22.14 ± 4.10<sup>c</sup> | 116.52 ± 17.25<sup>a</sup> | 20.904<sup>*</sup> |
| *M. spicata* | 4.68 ± 0.21<sup>a</sup> | 4.28 ± 0.18<sup>a</sup> | 0.91 ± 0.13<sup>b</sup> | 4.41 ± 0.24<sup>a</sup> | 3.117<sup>*</sup> |
| *M. longifolia* | 6.60 ± 0.40<sup>a</sup> | 6.10 ± 0.33<sup>a</sup> | 1.39 ± 0.24<sup>c</sup> | 3.69 ± 0.55<sup>b</sup> | 5.991<sup>*</sup> |

Means within rows, followed by the same letter are not significantly different (Tukey, $P \leq 0.05$ ).
(\*) significant at $P \leq 0.05$
<sup>z</sup> Number of samples during experimental duration

Diversity indices have reflected the three pest species populations preferences. The dominance (D) was the highest in case of *B. oleracea var palmifolia* leaves (D= 0.423 in 2016 and 2017), the highest species richness (1/D= 2.362 in 2016 and 2017), and the highest Shannon-Winner diversity index (H’= 0.933 in 2016, and 0.935 in 2017) in both seasons (Table 2). Despite the occurrence of pest populations; species distribution was varied for each population. When *B. tabaci* has recorded the highest density distribution on the four tested plants χ2= 734.223, *P*= 0.000) in 2016, *T. tabaci* χ2= 3730.706, *P*= 0.000) in 2017, and *T. urticae* has a fluctuated significant distribution among plants in both seasons (2016: χ2= 5370.34, *P*= 0.000; 2017: χ2 805.3225, *P*= 0.000) (Table 3).

**Table 2.** Diversity indices of pest species infesting Brassicaceae and Lamiaceae in Om Saber experimental site 2016 - 2017.

|  | 2016 |  |  |  | 2017 |  |  |  |
| --- | --- | --- | --- | --- | --- | --- | --- | --- |
|  | *B. napus* | *B. oleracea* | *M. spicata* | *M. longifolia* | *B. napus* | *B. oleracea* | *M. spicata* | *M. longifolia* |
| Individuals | 6547 | 4974 | 251 | 306 | 6458 | 5019 | 335 | 423 |
| Simpson (D) | 0.448 | 0.423 | 0.488 | 0.586 | 0.436 | 0.423 | 0.525 | 0.600 |
| Simpson (1/D) | 2.233 | 2.362 | 2.050 | 1.706 | 2.296 | 2.367 | 1.911 | 1.671 |
| Shannon-Wiener (H') | 0.873 | 0.933 | 0.833 | 0.730 | 0.904 | 0.935 | 0.788 | 0.711 |
| Evenness (e<sup>H/S</sup>) | 0.798 | 0.847 | 0.767 | 0.692 | 0.823 | 0.849 | 0.733 | 0.679 |
| Brillouin | 0.869 | 0.930 | 0.790 | 0.677 | 0.901 | 0.930 | 0.760 | 0.683 |
| Menhinick | 0.037 | 0.043 | 0.189 | 0.171 | 0.037 | 0.042 | 0.164 | 0.146 |
| Margalef | 0.228 | 0.235 | 0.362 | 0.349 | 0.228 | 0.235 | 0.344 | 0.331 |
| Jaccard's index (J') | 0.794 | 0.849 | 0.758 | 0.665 | 0.823 | 0.851 | 0.717 | 0.648 |
| Fisher alpha | 0.300 | 0.34 | 0.479 | 0.461 | 0.301 | 0.309 | 0.454 | 0.436 |
| Berger-Parker | 0.483 | 0.484 | 0.615 | 0.737 | 0.487 | 0.487 | 0.667 | 0.748 |

Diversity indices were tested at $P \leq 0.05$

**Table 3.** Pest Species distribution in Om Saber location 2016 - 2017.

|  | *B. napus* | *B. oleracea* | *M. spicata* | *M. longifolia* |  |  |  |
| --- | --- | --- | --- | --- | --- | --- | --- |
| Pest Species | 2016 | | | | Variance | $\chi^2$ | *P* |
|  | Species distribution |  |  |  |  |  |  |
| *T. tabaci* | ++ | + | +++ | +++ | 2005760 | 4372.649 | 0.000 |
| *T. urticae* | +++ | +++ | + | ++ | 2557160 | 5370.342 | 0.000 |
| *B. tabaci* | +++ | +++ | +++ | +++ | 52963.15 | 734.223 | 0.000 |
| Pest Species | 2017 | | | | Variance | $\chi^2$ | *P* |
|  | Species distribution |  |  |  |  |  |  |
| *T. tabaci* | +++ | +++ | +++ | +++ | 1705812 | 3730.706 | 0.000 |
| *T. urticae* | +++ | +++ | ++ | ++ | 66332.41 | 805.3225 | 0.000 |
| *B. tabaci* | ++ | ++ | +++ | +++ | 2533514 | 5274.391 | 0.000 |

Distribution variance and $\chi^2$ were tested at $P \leq 0.05$

#### b. Kom Oshim location

The population dynamics of *T. urticae, B. tabaci* and *T. tabaci* have resulted significant differentiations among tested plants. The highest mean population recorded in Brassicaceae was in *B. napus var pabularia* (*F*= 80.963, *P*= 0.000), when *T. urticae* recorded 35.17 ± 2.91 adults/10 leaves, and *B. tabaci* 67.54 ± 7.60 individuals/10 leaves in 2016. While, *T. tabaci* highest mean population was recorded in *B. aleracea var. plamifolia* (11.04 ± 2.62 individuals/10 leaves) (Table 4). The second season showed a similarity in population density dynamics, and the recorded results were statistically significant on probability level of 95% (*B. napus var pabularia F*= 50.413; *B. aleracea var. plamifolia F*= 43.405; *M. spicata F*= 5.410; M. longifolia *F*= 7.462) (Table 4).

**Table 4.** Pest species populations on four medicinal plant species (mean/10 leaves ± SE) in Kom Oshim experimental site (2016 - 2017)

| 2016 |  |  |  |  |  |  |  |
| --- | --- | --- | --- | --- | --- | --- | --- |
|  | *T. urticae* (18)<sup>z</sup> |  | *T. tabaci* |  | *B. tabaci* |  | *F*-test |
| | Nymphs | Adults | N | Mean ± SE | N | Mean ± SE | |
| *B. napus* | 37.90 ± 2.52<sup>b</sup> | 35.17 ± 2.91<sup>c</sup> | 16 | 8.16 ± 1.90<sup>d</sup> | 18 | 67.54 ± 7.60<sup>a</sup> | 80.963<sup>*</sup> |
| *B. oleracea* | 35.12 ± 2.59<sup>b</sup> | 30.82 ± 2.51<sup>c</sup> | 16 | 11.04 ± 2.62<sup>d</sup> | 18 | 48.94 ± 6.42<sup>a</sup> | 82.651<sup>*</sup> |
| *M. spicata* | 5.75 ± 0.31<sup>a</sup> | 2.32 ± 0.26<sup>b</sup> | 10 | 1.05 ± 0.17<sup>c</sup> | 14 | 1.92 ± 0.25<sup>bc</sup> | 6.021<sup>*</sup> |
| *M. longifolia* | 9.92 ± 0.76<sup>a</sup> | 3.24 ± 0.23<sup>b</sup> | 10 | 1.23 ± 0.15<sup>c</sup> | 14 | 1.27 ± 0.25<sup>c</sup> | 9.736<sup>*</sup> |
| 2017 |  |  |  |  |  |  |  |
|  | *T. urticae* (18)<sup>z</sup> |  | *T. tabaci* |  | *B. tabaci* |  | *F*-test |
| | Nymphs | Adults | N | Mean ± SE | N | Mean ± SE | |
| *B. napus* | 40.26 ± 3.14 b | 36.38 ± 2.77 c | 16 | 15.16 ± 1.90 d | 18 | 62.54 ± 7.60 a | 50.413<sup>*</sup> |
| *B. oleracea* | 38.14 ± 3.23 b | 35.39 ± 2.50 c | 16 | 13.04 ± 2.62 d | 18 | 52.94 ± 6.42 a | 43.405<sup>*</sup> |
| *M. spicata* | 8.24 ± 0.36 a | 5.31 ± 0.26 c | 10 | 4.33 ± 0.17 d | 14 | 7.98 ± 0.25 b | 5.410<sup>*</sup> |
| *M. longifolia* | 11.14 ± 0.86 a | 6.53 ± 0.23 b | 10 | 3.44 ± 0.15 c | 14 | 6.44 ± 0.25 b | 7.462<sup>*</sup> |

Means within rows, followed by the same letter, are not significantly different (Tukey, $P \leq 0.05$ ).
(\*) significant at $P \leq 0.05$
<sup>z</sup> Number of samples during experimental duration

Diversity indices have not been changed too much between two seasons, when the sampled individuals were significantly differed. Although they have given a real indication of pest species abundance, richness and preference due to their significant values of dominance (D), species richness (1/D), diversity (H’), similarities (J’) and evenness (e^H/S) (Table 5). Besides, the species distribution of each species has statistically significant differences, when *B. tabaci* (2016 χ2= 257.95; 2017 χ2= 299.50, *P*= 0.000) and *T. tabaci* (2016 χ2= 1968.09; 2017 χ2= 1971.23, *P*= 0.000) were the most distributed species on the four tested plants in two experimental seasons, and *T. urticae* has recorded a moderate distribution on Brassicaceae and *M. spicata*, and heavy occurrence on *M. longifolia* in 2016 (χ2= 1575.60, *P*= 0.000), and heavily distribution on Lamiaceae plants only in the second season χ2= 219.82, *P*= 0.000) (Table 6).

**Table 5.** Diversity indices of pest species infesting Brassicaceae and Lamiaceae in Kom Oshim (2016 - 2017)

|  | 2016 |  |  |  | 2017 |  |  |  |
| --- | --- | --- | --- | --- | --- | --- | --- | --- |
|  | *B. napus* | *B. oleracea* | *M. spicata* | *M. longifolia* | *B. napus* | *B. oleracea* | *M. spicata* | *M. longifolia* |
| **Individuals** | 2593 | 2243 | 181 | 265 | 3503 | 3147 | 1560 | 1516 |
| **Simpson (D)** | 0.455 | 0.440 | 0.656 | 0.793 | 0.499 | 0.515 | 0.949 | 0.955 |
| **Simpson (1/D)** | 2.196 | 2.274 | 1.524 | 1.261 | 2.005 | 1.942 | 1.054 | 1.047 |
| **Shannon-Wiener (H')** | 0.856 | 0.904 | 0.627 | 0.429 | 0.805 | 0.807 | 0.138 | 0.124 |
| **Evenness (e<sup>H/S</sup>)** | 0.784 | 0.8232 | 0.624 | 0.512 | 0.745 | 0.747 | 0.383 | 0.378 |
| **Brillouin** | 0.780 | 0.823 | 0.571 | 0.39 | 0.798 | 0.802 | 0.13 | 0.115 |
| **Menhinick** | 0.850 | 0.895 | 0.558 | 0.372 | 0.051 | 0.053 | 0.076 | 0.077 |
| **Margalef** | 0.059 | 0.063 | 0.222 | 0.184 | 0.245 | 0.248 | 0.272 | 0.273 |
| **Jaccard's index (J')** | 0.254 | 0.259 | 0.385 | 0.358 | 0.733 | 0.735 | 0.126 | 0.113 |
| **Fisher alpha** | 0.335 | 0.341 | 0.51 | 0.473 | 0.323 | 0.327 | 0.358 | 0.359 |
| **Berger-Parker** | 0.507 | 0.529 | 0.794 | 0.884 | 0.622 | 0.659 | 0.974 | 0.977 |

Diversity indices were tested at $P \leq 0.05$

**Table 6.** Pest Species distribution in Kom Oshim location 2016 - 2017.

| Pest Species | *B. napus* | *B. oleracea* | *M. spicata* | *M. longifolia* | Variance | $\chi^2$ | *P* |
| --- | --- | --- | --- | --- | --- | --- | --- |
|  | 2016 |  |  |  |  |  |  |
|  | Species distribution |  |  |  |  |  |  |
| *T. urticae* | ++ | ++ | ++ | +++ | 378705 | 1575.60 | 0.000 |
| *B. tabaci* | +++ | +++ | +++ | +++ | 7091.29 | 257.95 | 0.000 |
| *T. tabaci* | +++ | +++ | +++ | +++ | 340108.5 | 1968.09 | 0.000 |
| Pest Species | 2017 | | | | Variance | $\chi^2$ | *P* |
|  | Species distribution |  |  |  |  |  |  |
|  | Species distribution |  |  |  |  |  |  |
| *T. urticae* | ++ | ++ | +++ | +++ | 132967.08 | 219.82 | 0.000 |
| *B. tabaci* | +++ | +++ | +++ | +++ | 9860.65 | 299.50 | 0.000 |
| *T. tabaci* | +++ | +++ | +++ | +++ | 341292.63 | 1971.23 | 0.000 |

Distribution variance and $\chi^2$ were tested at $P \leq 0.05$

### 2. The proposed IPM tactic for resulted pests

Using the predatory phytoseiid species was efficient to reduce the *T. urticae, B. tabci*, and *T. tabaci* populations on *B. napus var. pabularia, B. oleracea var palmifolia, M. spicata* and *M. longifolia*. Releasing *P. persimilis* and *A. swirskii* were effectively reduced the mean number of *T. urticae* in both Brassicaceae and Lamiaceae tested plants comparing with the control. *Amblyseius swirskii* was effective not only towards TSSM infestations, but also to reduce *T. tabaci* and *B. tabaci*. While *C. negevi* was less effective comparing with the two other phytoseiids, the average reduction percentage of releasing *C. negevi* against *T. urticae, T. tabaci*, and *B. tabaci* on the four tested plant species was 50% in Om Saber and 55% in Kom Oshim (Tables 7-10) (Suppl. 1).

**Table 7.**
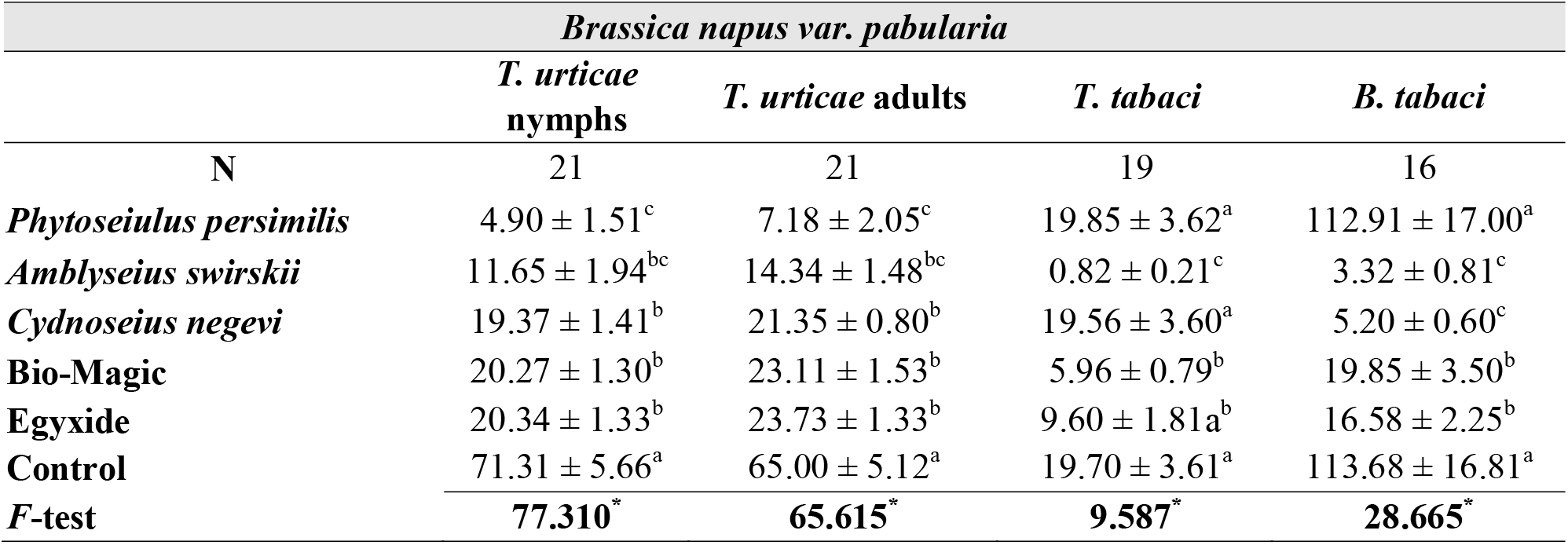
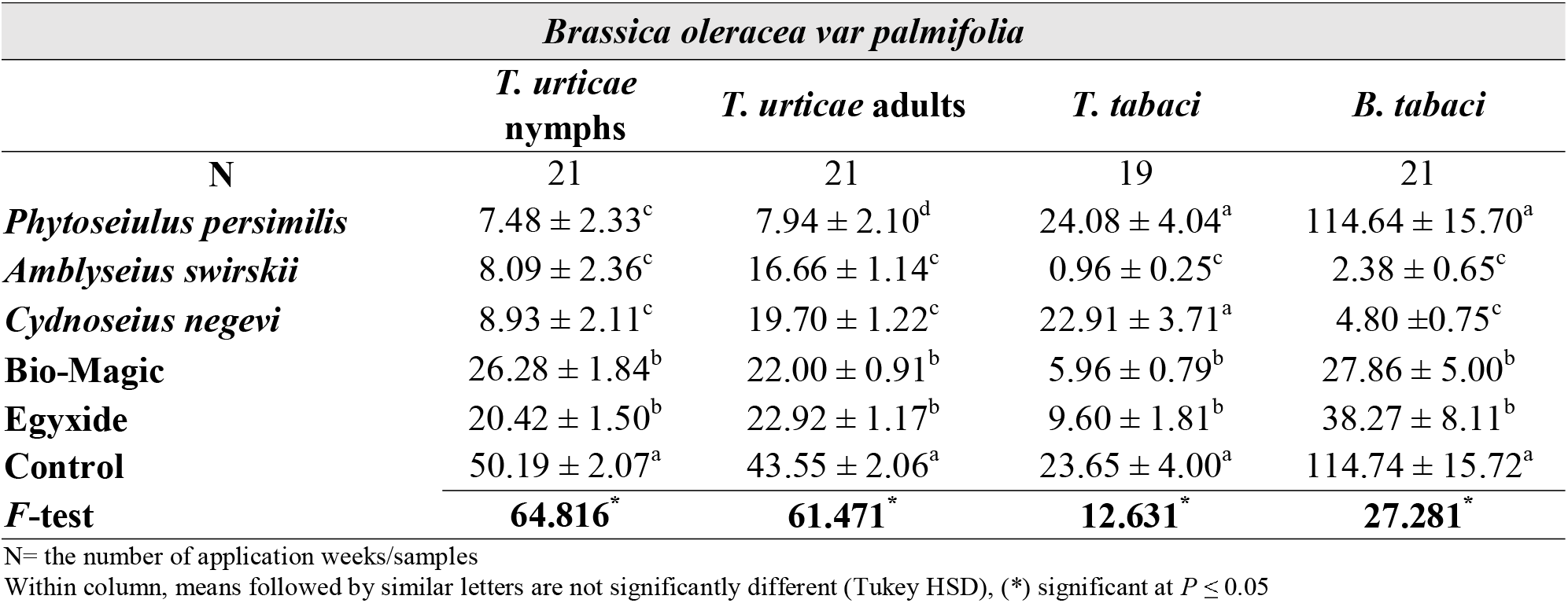
Mean number (± SE) of Brassicaceae pests after proposed application, Om Saber experimental site 2017.

**Table 8.** Mean number (± SE) of Lamiaceae pests after proposed application, Om Saber experimental site 2017.

| *Mentha spicata* |  |  |  |  |
| --- | --- | --- | --- | --- |
|  | *T. urticae* nymphs | *T. urticae* adults | *T. tabaci* | *B. tabaci* |
| N | 21 | 21 | 21 | 21 |
| *Phytoseiulus persimilis* | $0.29 \pm 0.13^c$ | $0.40 \pm 0.14^d$ | $1.10 \pm 0.11^a$ | $5.48 \pm 0.40^a$ |
| *Amblyseius swirskii* | $0.46 \pm 0.16^c$ | $0.68 \pm 0.14^{cd}$ | $0.04 \pm 0.02^c$ | $0.95 \pm 0.31^c$ |
| *Cydnoseius negevi* | $0.52 \pm 0.15^c$ | $1.52 \pm 0.05^b$ | $1.05 \pm 0.10^a$ | $0.87 \pm 0.22^c$ |
| Bio-Magic | $1.43 \pm 0.20^b$ | $0.97 \pm 0.06^c$ | $0.52 \pm 0.10^b$ | $2.53 \pm 0.27^b$ |
| Egyxide | $1.29 \pm 0.19^b$ | $0.95 \pm 0.07^c$ | $0.70 \pm 0.12^b$ | $3.23 \pm 0.37^b$ |
| Control | $3.74 \pm 0.20^a$ | $3.06 \pm 0.14^a$ | $1.08 \pm 0.11^a$ | $5.27 \pm 0.34^a$ |
| **F-test** | **56.907\*** | **77.717\*** | **17.946\*** | **38.640\*** |
| *Mentha longifolia* |  |  |  |  |
|  | *T. urticae* nymphs | *T. urticae* adults | *T. tabaci* | *B. tabaci* |
| N | 14 | 14 | 14 | 14 |
| *Phytoseiulus persimilis* | $0.32 \pm 0.15^c$ | $0.41 \pm 0.16^d$ | $1.63 \pm 0.19^a$ | $4.75 \pm 0.43^a$ |
| *Amblyseius swirskii* | $0.58 \pm 0.17^c$ | $1.20 \pm 0.15^c$ | $0.01 \pm 0.01^c$ | $0.62 \pm 0.21^c$ |
| *Cydnoseius negevi* | $0.91 \pm 0.18^c$ | $2.06 \pm 0.07^b$ | $1.60 \pm 0.18^a$ | $1.14 \pm 0.21^c$ |
| Bio-Magic | $2.10 \pm 0.15^b$ | $2.44 \pm 0.20^b$ | $0.01 \pm 0.00^b$ | $1.66 \pm 0.15^b$ |
| Egyxide | $2.27 \pm 0.16^b$ | $1.97 \pm 0.12^b$ | $0.60 \pm 0.15^c$ | $1.96 \pm 0.24^b$ |
| Control | $5.33 \pm 0.38^a$ | $4.70 \pm 0.33^a$ | $1.62 \pm 0.18^a$ | $3.88 \pm 0.25^a$ |
| **F-test** | **72.997\*** | **59.199\*** | **30.995\*** | **37.860\*** |

N= the number of application weeks
The means followed by similar letters in the column are not significantly different (Tukey HSD), (\*) significant at $P \leq 0.05$

**Table 9.**
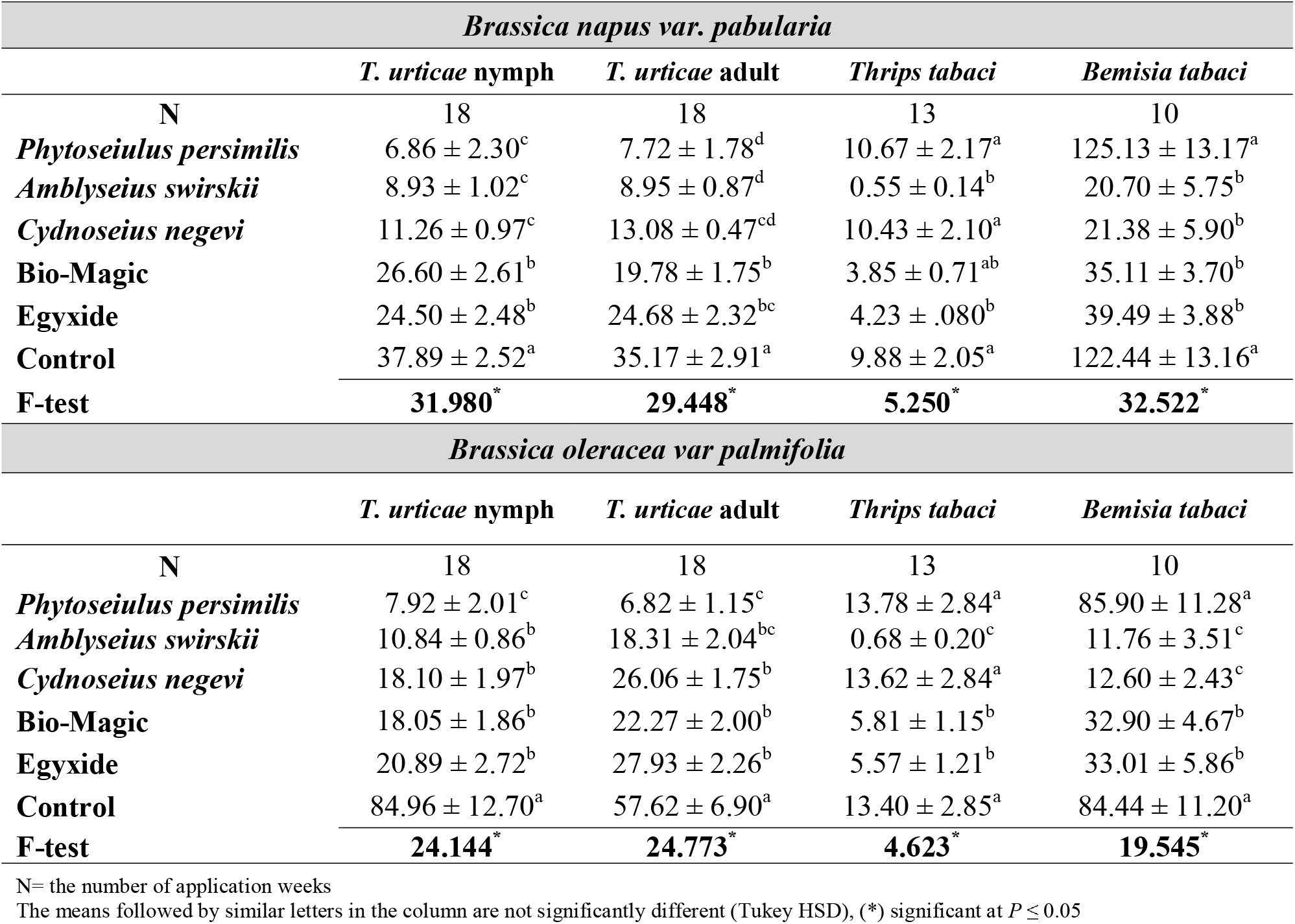
Mean number (± SE) of Brassicaceae pests after proposed application in Kom Oshim experimental site 2017.

**Table 10.** Mean number (± SE) of Lamiaceae pests after proposed application in Kom Oshim experimental site 2017.

| *Mentha spicata* |  |  |  |  |
| --- | --- | --- | --- | --- |
|  | *T. urticae*<br>nymphs | *T. urticae* adult | *T. tabaci* | *B. tabaci* |
| N | 18 | 18 | 13 | 10 |
| *Phytoseiulus persimilis* | 1.01 ± 0.32<sup>c</sup> | 1.43 ± 0.30<sup>d</sup> | 1.71 ± 0.20<sup>a</sup> | 4.17 ± 0.31<sup>a</sup> |
| *Amblyseius swirskii* | 1.13 ± 0.24<sup>b</sup> | 2.15 ± 0.24<sup>d</sup> | 0.25 ± 0.12<sup>b</sup> | 0.54 ± 0.20<sup>d</sup> |
| *Cydnoseius negevi* | 0.96 ± 0.20<sup>b</sup> | 2.32 ± 0.30<sup>c</sup> | 1.30 ± 0.14<sup>a</sup> | 0.57 ± 0.15<sup>d</sup> |
| Bio-Magic | 1.23 ± 0.13<sup>b</sup> | 3.55 ± 0.10<sup>b</sup> | 0.60 ± 0.10<sup>b</sup> | 1.78 ± 0.10<sup>c</sup> |
| Egyxide | 1.06 ± 0.16<sup>b</sup> | 3.13 ± 0.12<sup>bc</sup> | 0.63 ± 0.07<sup>b</sup> | 1.80 ± 0.07<sup>c</sup> |
| Control | 3.77 ± 0.22<sup>a</sup> | 5.75 ± 0.34<sup>a</sup> | 1.27 ± 0.14<sup>a</sup> | 2.42 ± 0.11<sup>b</sup> |
| **F-test** | **36.327\*** | **40.602\*** | **13.811\*** | **63.012\*** |
| *Mentha longifolia* |  |  |  |  |
|  | *T. urticae*<br>nymphs | *T. urticae* adult | *T. tabaci* | *B. tabaci* |
| N | 18 | 18 | 13 | 10 |
| *Phytoseiulus persimilis* | 0.64 ± 0.30<sup>d</sup> | 0.98 ± 0.27<sup>d</sup> | 1.75 ± 0.11<sup>a</sup> | 3.69 ± 0.25<sup>a</sup> |
| *Amblyseius swirskii* | 0.77 ± 0.25<sup>c</sup> | 1.42 ± 0.24<sup>c</sup> | 0.32 ± 0.12<sup>c</sup> | 0.11 ± 0.03<sup>b</sup> |
| *Cydnoseius negevi* | 1.63 ± 0.10<sup>bc</sup> | 2.31 ± 0.10<sup>bc</sup> | 1.41 ± 0.11<sup>a</sup> | 0.11 ± 0.02<sup>b</sup> |
| Bio-Magic | 3.47 ± 0.31<sup>b</sup> | 2.46 ± 0.30<sup>b</sup> | 0.75 ± 0.10<sup>b</sup> | 0.22 ± 0.07<sup>b</sup> |
| Egyxide | 3.15 ± 0.30<sup>bc</sup> | 2.15 ± 0.29<sup>bc</sup> | 0.81 ± 0.08<sup>b</sup> | 0.22 ± 0.06<sup>b</sup> |
| Control | 5.03 ± 0.20<sup>a</sup> | 3.43 ± 0.20<sup>a</sup> | 1.43 ± 0.11<sup>a</sup> | 3.92 ± 0.21<sup>a</sup> |
| **F-test** | **25.858\*** | **19.857\*** | **21.880\*** | **37.201\*** |

Within column, means followed by similar letters are not significantly different (Tukey HSD), (\*) significant at $P \leq 0.05$

Weather data were recorded during the experimental duration period; the average temperature was 28° C in Om Saber and Kom Oshim, and the relative humidity was 54% in Om Saber and 52% in Kom Oshim. The least reduction percentage was recorded in Bio-Magic and Egyxide applications. These two means were replicated (spray application) three times every three weeks due to their low performance. The least reduction percentage recorded in case of using Bio-Magic was 34% in Om Saber, and 40% in Kom Oshim, and in case of Egyxide it was recorded 40% in Om Saber and 45% in Kom Oshim (Tables 11) (Suppl. 1).

**Table 11.** Reduction percentages of control applications. An evaluation between 2017 and 2018 results using Student’s test and ANOVA.

|  | Om Saber |  |  | Kom Oshim |  |  |
| --- | --- | --- | --- | --- | --- | --- |
|  | 2017 | 2018 | T-test | 2017 | 2018 | T-test |
| *Phytoseiulus persimilis* | 88% | 85% | **1.50<sup>ns</sup>** | 90% | 88% | **1.042<sup>ns</sup>** |
| *Amblyseius swirskii* | 75% | 73% | **1.000<sup>ns</sup>** | 80% | 80% | - |
| *Cydnoseius negevi* | 50% | 51% | **0.561<sup>ns</sup>** | 55% | 57% | **1.035<sup>ns</sup>** |
| Bio-Magic | 34% | 35% | **0.675<sup>ns</sup>** | 40% | 40% | - |
| Egyxide | 40% | 42% | **1.860<sup>ns</sup>** | 45% | 44% | **0.809<sup>ns</sup>** |
| **ANOVA** | **40.667\*** | **41.025\*** |  | **46.145\*** | **45.004\*** |  |

T-test df= 18; $P \leq 0.05$

A Kruskal-Wallis test was carried out to determin which hypothesis to accept; either the null hypothesis (*H_0_*) (which suggested that there were no significant differences between seasons 2017 and 2018), or the alternative hypothesis (*H_1_*) (which suggested that there were significant differences), at confidence level = 95%. Kruskal-Wallis resulted was to reject the alternative hypothesis *H_1_* and accept the null hypothesis *H_0_*. Results obtained in 2018 of the two experimental locations were not significantly different, when compared with those gathered in 2017 using Student’s test and ANOVA. Bio-Magic and Egyxide have the least reduction percentage resulted, when *A. swirskii* was the most effective for control targeted pests, then *C. negevi* which success in suppression *B. tabaci* and *T. urticae* populations. The specialist *P. persimilis* showed a successful application toward TSSM populations in both experimental locations in the two seasons (Table 11).

## DISCUSSION

Four principles of integrated pest management in an Agro-ecosystem; prevention, monitoring, avoidance, and suppression (EPA, 2021). According to these tactics, the current study was designed. The prevention phase was occurred by managing weeds which were found in each location, by mechanical removal. However, diverse weed species have been recorded in both locations; e.g., the slender amaranth *Amaranthus viridis* L. (Amaranthaceae), the Cheese weed *Malva parviflora* L. (Malvaceae), the burning nettle *Urtica urens* L. (Urticaceae), and the common cocklebur *Xanthium strumarium* L. (Asteraceae). These shared weeds have been employed as shelters for the recorded pests, besides, other secondary lepidopterans, aphids, and other phytophagous tydied, tarsonemid, and tenuipalpid mite species. Despite, some studies considered weeds not much harm as prospected and they might do another ecological role in an ecosystem; as being shelter/banker plants for predatory species (Parolin *et al*., 2012; 2013), and/or trap plants for herbivore species as applied in commercial applications in Europe (Shelton and Badenes-Pérez, 2006) which had been vital for the entire process (Soloneski and Larramendy, 2013).

Monitoring, as a second tactic, was taking place alongside during the experiment. Detecting climatic factors was very essential for most of the phytophagous species such as spider mites (Praslička and Huszár, 2004; White and Liburd, 2005; Zou *et al*., 2018), whiteflies (Jha and Kumar, 2017; Khan, 2019; Gamarra *et al*., 2020; Chandi *et al*., 2021), thrips (McDonald *et al*., 1998; Bergant *et al*., 2005; Cao *et al*., 2018; Garrick and Liburd, 2018), also for predacious mite (Tixier, 2018; Urbaneja-Bernat and Jaques, 2022), as in the current study. As well, for predatory insect (Schuldiner-Harpaz and Coll, 2013), or spider (Blamires and Sellers, 2019; Napiórkowska *et al*., 2021) species, and even for the stored product and public health pests (Beckett, 2011; Mahakittikun *et al*., 2011) life table parameters, population dynamics, and behaviours which affected due to climate changes.

The present study recorded the mean numbers of *T. urticae, T. tabaci*, and *B. tabaci* populations, which were significantly varied, increased, and changed (Tables 1 and 4). However, *T. urticae* population growth was not significantly changed in the two experimental seasons, where the fluctuation of pest distribution was affected (*T. urticae* immatures *T*=0.719, *P*=0.474; *T. urticae* adults *T*=1.011, *P*=0.314). While *T. tabaci*, and *B. tabaci* populations growth were highly significant increased (*T. tabaci T*=3.999, *P*=0.000; *B. tabaci T*=6.086, *P*=0.000) (Supplementary).

Abou-Elella *et al*., (2021) have concluded that organic fertilization could affect the life table parameters of the TSSM positively. That helps in explaining why these pest species built their populations rapidly and increase their fluctuations that became higher than the EIL, which needed to be controlled. Climatic factors, also, affect the tri-trophic interactions, by changing the chemical responses of herbivore-plant and predator-prey interactions (Laws, 2017), which reflects on the ecosystem and any possible IPM program may applied.

Despite, climatic factors were not only the reason that causes pest populations to increase; variations could be occurred due to different abiotic (e.g., plant nutritional contents, soil fertilization plant, and climatic factors), and biotic factors (e.g., plant morphological characters, herbivore physiological characters, plant-herbivore relation, and herbivore-herbivore relation) (Laws, 2017; Skendžić *et al*., 2021; Zidan, 2021).

Together, these factors are sustainable for the agroecosystem balance and suppression of harmful pests. As faunal and floral diversities, which have substantial roles in pest and disease management in the Agro-ecosystems (Westerman *et al*., 2003; Hajjar *et al*., 2008).

Another example to understand the increasing of pest fluctuations and abundance, the presence of primary and secondary pests on the weed species in both sites; as the monitoring has helped in indicating the most abundance and dominant pest species for control applications (secondary pest species have been statistically neglected due to not causing significant damages). As the mathematical modulation of diversity indices have measured the community species diversity, and diversity differences in populations within the ecosystem (Magurran, 2004).

Avoidance, as the third protocol, was carried for maintaining the predatory species, keeping the pest populations under the EIL/ETL, and to reduce/prevent the need for chemical application. Some arthropods (e.g., *T. urticae*) could build resistant generations toward chemical compound (Ramasubramanian *et al*., 2005), as a specialist predator; *P. persimilis* was very efficient towards *T. urticae* (reduction percentage average= 90% in the whole study), and useless for other pests, due to its diet specialty (McMurtry and Croft, 1997; McMurtry *et al*., 2013). To keep other pest species under the EIL, therefore, we used the predatory species; *A. swirskii* and *C. negevi* which were the best candidates, as an indigenous species in the Mediterranean basin, and have a wide range of feeding preferences (e.g., whiteflies, thrips, eriophyid and tetranychid pests) in both open fields and greenhouses (Nomikou *et al*., 2001; Wimmer *et al*., 2008; Arthurs *et al*., 2009; Stansly and Castillo, 2010; Calvo *et al*., 2011; Doğramaci *et al*., 2011; Onzo *et al*., 2012; Xiao *et al*., 2012; Negm *et al*., 2014; Alatawi *et al*., 2018; Sanad and Hassan, 2019).

Interactions such as competition, intraguild predation (IGP), extraguild predation, and cannibalism would affect the plant-prey-predator relations in the IPM procedure, thus using multiple predatory species different in their dietary preferences and predatory behaviour is better than using single ones (Schausberger and Walzer, 2001; Momen, 2010; Momen and El-Borolossy, 2010; Momen *et al*., 2013; Guo *et al*., 2016; Knapp *et al*., 2018; Döker *et al*., 2021; Momen and Abdel-Khalek, 2009 a, b, 2021).

Otherwise, the reduction percentages of Bio-Magic and Egyxide compounds in both locations were negative (especially in July and August) (Supplementary file). This indicated that using these methods in that time was not useful, despite, their active ingredients are natural origin (*M. anisopliue* fungi and plant essential oils). It was stated that 28 °C and a photoperiod of 16 hours were the most suitable conditions for *M. anisopliue* (Alves *et al*., 1984). Although, the UV-A and UV-B of solar waves radiation extremely reduce *M. anisopliue* capability to form conidia which cause targeted pest mortality (Francisco *et al*., 2008). Besides, storage, transportation, and field application are also limitation factors (Parra, 2014; Sinha *et al*., 2016).

The suppression, the final tactic in the present study; represented by *A. swirskii* and *C. negevi* results in reducing whitefly and thrips populations in both Brassicaceae and Lamiaceae plantations. Also, the role of *P. persimilis* as a specialist for *T. urticae*, and hypothetically, therefore, getting a large loss if this specialist is used as a “solo” tool.

The combination of using these three phytoseiid species is successful in the Agro-ecosystem, but we should take in consideration the release timing and capacity (Knapp *et al*., 2018; Fonseca *et al*., 2020). The combination may include Egyxide and/or Bio-Magic means to suppress targeted pest populations below the EIL and ETL, however, they might be failure methods if used individually; due to degradation by weather conditions such as temperature, humidity, daylight UV waves, and wind speed (Allam *et al*., 2020).

## CONCLUSION

Within an Agro-ecosystem, abiotic and biotic factors together help in explaining why various pest species build their communities rapidly and increase their parameters that become above the EIL. Such factors are hypothesized to affect the plant-arthropod, predator-herbivore, predator-predator, and tri-trophic interactions. We proposed such IPM tactic aimed to help in the suppression of pest infestations. The study is recommending that application of such protocol should consider the timing of tacking an action and merging tactics together to get the maximum efficiency.

## ACKNOWLEDGMENTS

The authors express their gratitude due to Dr. Mohamed A. Gesraha, Prof. of Entomology, Pests and Plant Protection Department (NRC), for his valuable comments of the statistical analysis and data preparation. Authors would dedicate this work for the late Prof. Awad A. F. Al-Bahrawy, Agricultural Zoology Department, Suez Canal University, may his soul rest in peace. Deep thanks are due to Mr. Khaled A. Dawoud, technician, Pests and Plant Protection Department (NRC), for his valuable assisting during this work.

## DATA AVAILABILITY

The data that support the findings of this study are available and attached in the supplementary file.

## REFERENCES

Abdel-Khalek, A.A. & Momen, F.M. (2022). Biology and life table parameters of *Proprioseiopsis lindquisti* on three eriophyid mites (Acari: Phytoseiidae: Eriophyidae). Persian Journal of Acarology, 11(1): 59–69. Available at https://www.biotaxa.org/pja/article/view/68574/70141.

Abou-Awad, B.A., Afia, S.I. & El-Saiedy, E.M.A. (2017). Efficiency of two preadatory phytoseiid mites, biopesticide and fungal pathogen for controlling *Tetranychus urticae* Koch (Acari: Tetranychidae) on watermelon and muskmelon at Behera Governorate Egypt. Bioscience Research, 14(4): 1042–1049. Available at https://www.isisn.org/BR-14-2017/1042-1049-14(4)2017BR-1566.pdf

Abou-Elella, G.M., Hassan, M.F., Nawar, M.S. & Zidan, I.M. (2014). Survival, development and reproduction of *Euseius finlandicus* (Oudemans) (Acari: Phytoseiidae) fed on various kinds of food substances. Archives of Phytopathology and Plant Protection, 47(7): 857-868. https://doi.org/10.1080/03235408.2013.823715.

Akmal, M., Freed, S., Malik, M.N. & Gul, H.T. (2013). Efficacy of *Beauveria bassiana* (Deuteromycotina: Hypomycetes) against different aphid species under laboratory conditions. Pakistan Journal of Zoology, 45, 71–78.

Alba, J.M., Bleeker, P.M., Glas, J.J., Schimmel, B.C.J., van Wijk, M., Sabelis, M.W., Schuurink, R.C. & Kant, M.R. (2012). The Impact of Induced Plant Volatiles on Plant-Arthropod Interactions. Pp: 15–73. In: G. Smagghe & I. Diaz (eds). Arthropod-Plant Interactions. Progress in Biological Control, vol 14. Springer, Dordrecht. https://doi.org/10.1007/978-94-007-3873-7_2.

Allam S.F., Mahmoud M.A-E., Hassan M.F. & Mabrouk A.H. (2020). Field application of six commercial essential oils against Date Palm mite, *Phyllotetranychus aegypticus* (Acari: Tenuipalpidae) in Egypt. Persian Journal of Acarology, 9(4): 377–389. Available at https://www.biotaxa.org/pja/article/view/202048.

Alves, S.B., Risco, S.H. & Almeida, L.C. (1984). Influence of photoperiod and temperature on the development and sporulation of *Metarhizium anisopliae* (Metsch.). Journal of Applied Entomology, 97(1-5): 127–129. https://doi.org/10.1111/j.1439-0418.1984.tb03726.x

Arthurs, S., Mckenzie, C.L., Chen, J., Dogramaci, M., Brennan, M., Houben, K. & Osborne, L. (2009) Evaluation of *Neoseiulus cucumeris* and *Amblyseius swirskii* (Acari: Phytoseiidae) as biological control agents of chilli thrips, *Scirtothrips dorsalis* (Thysanoptera: Thripidae) on pepper. Biological Control, 49(1):91–96. https://doi.org/10.1016/j.biocontrol.2009.01.002.

Beckett, S.J. (2011). Insect and mite control by manipulating temperature and moisture before and during chemical-free storage. Journal of Stored Products Research, 47(4): 284–292. https://doi.org/10.1016/j.jspr.2011.08.002.

Bergant, K., Stanislav, T., Znidarcic, D., Črepinšek, Z. & Kajfež-Bogataj, L. (2005). Impact of climate change on developmental dynamics of *Thrips tabaci* (Thysanoptera: Thripidae): Can it be quantified? Environmental Entomology, 34(4): 755–766. https://doi.org/10.1603/0046-225X-34.4.755

Blamires, S.J. & Sellers, W.I. (2019). Modelling temperature and humidity effects on web performance: implications for predicting orb-web spider *(Argiope* spp.) foraging under Australian climate change scenarios. Conservation Physiology, 7(1): coz083. https://doi.org/10.1093/conphys/coz083.

Cao, Y., Li, C., Yang, W.-J., Meng, Y.-L., Wang, L.-J., Shang, B.-Z. & Gao, Y.-L. (2018). Effects of temperature on the development and reproduction of *Thrips hawaiiensis* (Thysanoptera: Thripidae). Journal of Economic Entomology, 111(2): 755–760. https://doi.org/10.1093/jee/tox359

Chandi, R.S., Kataria, S.K. & Fand, B.B. (2021). Effect of temperature on biological parameters of cotton whitefly, *Bemisia tabaci* (Gennadius) (Hemiptera: Aleyrodidae). International Journal of Tropical Insect Science, 41: 1823–1833. https://doi.org/10.1007/s42690-020-00397-0

Delisle, J.F., Brodeur, J. & Shipp, L. (2015 a). Evaluation of various types of supplemental food for two species of predatory mites, *Amblyseius swirskii* and *Neoseiulus cucumeris* (Acari: Phytoseiidae). Experimental & Applied Acarology, 65(4): 483–494. https://doi.org/10.1007/s10493-014-9862-3.

Delisle, J.F., Shipp, L. & Brodeur, J. (2015 b). Apple pollen as a supplemental food source for the control of western flower thrips by two predatory mites, *Amblyseius swirskii* and *Neoseiulus cucumeris* (Acari: Phytoseiidae), on potted chrysanthemum. Experimental & Applied Acarology, 65(4): 495–509. https://doi.org/10.1007/s10493-014-9863-2.

de Melo, T.M.P., da Silva, E.M., da Silva, A.G., Vieira, G.H. & Lopes, PG. (2019). *Syzygium aromaticum* essential oil to the control of *Tenuipalpus heveae*. Journal of Agricultural Science, 11(8): 1916–9752. https://doi.org/10.5539/jas.v11n8p295.

Döker, I., Revynthi, A.M., Kazak, C. & Carrillo, D. (2021). Interactions among exotic and native phytoseiids (Acari: Phytoseiidae) affect biocontrol of two-spotted spider mite on papaya. Biological Control, 163: 104758. https://doi.org/10.1016/j.biocontrol.2021.104758.

Ebadollahi, A., Ziaee, M. & Palla, F. (2020). Essential oils extracted from different species of the Lamiaceae plants family as prospective bioagents against several detrimental pests. Molecules, 25: 1556. https://doi.or/10.3390/molecules25071556.

Ehi-Eromosele, C.O., Nwinyi, O.C. & Ajani, O.O. (2013). Integrated pest management. pp: 105–115 http://dx.doi.org/10.5772/54476. In: S. Soloneski and M. Larramendy. Weed and pest control - conventional and new challenges. IntechOpen, Available from https://www.intechopen.com/chapters/42758

El-Saiedy, E.M.A., Abou-Elella, G.M. & Alotaibi, S.A. (2008). Efficiency of three predatory phytoseiid mites and biocide chemical for controlling *Tetranychus urticae* Koch on egg plant at Beheira governorate. Research Journal of Agriculture and Biological Sciences, 4: 238–244.

El-Saiedy, ES.M. & Fahim, S.F. (2021). Evaluation of two predatory mites and acaricide to suppress *Tetranychus urticae* (Acari: Tetranychidae) on strawberry. Bulletin of the National Research Centre, 45: 97. https://doi.org/10.1186/s42269-021-00558-2.

El-Saiedy, E.M.A., Hassan, M.F., El-Bahrawy, A.F., El-Kady, G.A. & Kemel, M.S. (2015). Efficacy of two phytoseiid predators and a biopesticide against *Tetranychus curcubitacearum* (Sayed) (Acari: Tetranychidae) on eggplant at Ismailia Governorate, Egypt. Egyptian Journal of Biological Pest Control, 25: 71–74.

EPA (United States Environmental Protection Agency) (2021). Integrated Pest Management (IPM) Principles. A web page, Available at https://www.epa.gov/safepestcontrol/integrated-pest-management-ipm-principles.

Evans, H.C. (1982). Entomogenous fungi in tropical forest ecosystems: an appraisal. Ecological Entomology, 7: 47–60. https://doi.org/10.1111/j.1365-2311.1982.tb00643.x.

Ferreira, C.T., Noronha, A.C., Souza Neto, E.P., De Oliveira, R.P., Lins, P.M.P. & Batista, T.F.V. (2022). Functional and numerical responses of the predatory mite *Amblyseius aerialis* (Acari: Phytoseiidae) to *Aceria guerreronis* (Acari: Eriophyidae). Acarologia, 62(1): 27–35. https://doi.org/10.24349/w600-25ar.

Frago, E., Gols, R., Schweiger, R., Müller, C., Dicke, M. & Godfray, H.C.J. (2022). Herbivore-induced plant volatiles, not natural enemies, mediate a positive indirect interaction between insect herbivores. Oecologia, https://doi.org/10.1007/s00442-021-05097-1

Francisco, E.A., Rangel, D. & La Scala Jr, N. (2008). Exposure of *Metarhizium anisopliae* conidia to UV-B radiation reduces its virulence. Journal of Anhui Agricultural University, 35: 246–249.

Gamarra, H., Sporleder, M., Carhuapoma, P., Kroschel, J. & Kreuze, J. (2020). A temperature-dependent phenology model for the greenhouse whitefly *Trialeurodes vaporariorum* (Hemiptera: Aleyrodidae). Virus Research, 289: 198107. https://doi.org/10.1016/j.virusres.2020.198107.

Gardarin, A., Plantegenest, M., Bischoff, A. & Valantin-Morison, M. (2018). Understanding plant-arthropod interactions in multitrophic communities to improve conservation biological control: useful traits and metrics. Journal of Pest Science, 91(3): 943–955. https://doi.org/10.1007/s10340-018-0958-0.

Garrick, T.A., & Liburd, O.E. (2018). Impact of climate change on a key agricultural pest: Thrips. In: I. Management Association (Ed.). Climate change and environmental concerns: Breakthroughs in Research and Practice (pp. 65–87). IGI Global. http://doi:10.4018/978-1-5225-5487-5.ch004

Garrido-Jurado, I., Fernandez-Bravo, M., Campos, C., & Quesada-Moraga, E. (2015). Diversity of entomopathogenic Hypocreales in soil and phylloplanes of five Mediterranean cropping systems. Journal of Invertebrate Pathology, 130: 97–106. https://doi.org/10.1016/j.jip.2015.06.001.

Greenfield, M., Gomez-Jimenez, M. I., Ortiz, V., Vega, F. E., Kramer, M., & Parsa, S. (2016). *Beauveria bassiana* and *Metarhizium anisopliae* endophytically colonize cassava roots following soil drench inoculation. Biological Control, 95: 40–48. https://doi.org/10.1016/j.biocontrol.2016.01.002.

Guo, Y., Lv, J., Jiang, X., Wang, B., Gao, Y., Wang, E. & Xu, X. (2016). Intraguild predation between *Amblyseius swirskii* and two native Chinese predatory mite species and their development on intraguild prey. Scientific Reports, 6: 22992. https://doi.org/10.1038/srep22992

Halsch, C.A., Shapiro, A.M., Fordyce, J.A., Nice, C.C., Thorne, J.H., Waetjen, D.P. & Forister, M.L. (2021). Insects and recent climate change. Proceedings of the National Academy of Sciences of the United States of America, 118(2): e2002543117. https://doi.org/10.1073/pnas.2002543117

Henderson, C.E. & Tilton, E.W. (1995). Tests with acaricides against the brown wheat mites. Journal of Economic Entomology, 84(2): 157–161. https://doi.org/10.1093/jee/48.2.157

Isman, M.B. (2006). Botanical insecticides, deterrents, and repellents in modern agriculture and increasingly regulated world. Annual Review of Entomology, 51: 45–66. https://doi.org/10.1146/annurev.ento.51.110104.151146.

Isman, M.B. & Machial, C.M. (2006). Pesticides based on plant essential oils: from traditional practice to commercialization. In: M. Rai and M.C. Carpinella (eds.). Naturally occurring bioactive compounds, Elsevier, pp 29–44.

Janssen, A. & Sabelis, M.W. (2015) Alternative food and biological control by generalist predatory mites: the case of *Amblyseius swirskii*. Experimental & Applied Acarology, 65(4): 413–418. https://doi.org/10.1007/s10493-015-9901-8.

Jha, S.K. & Kumar, M. (2017). Effect of weather parameters on incidence of whitefly, *Bemisia tabaci* (Gennadius) on tomato. Journal of Entomology and Zoology Studies, 5(6): 304–306. Available at https://www.entomoljournal.com/archives/2017/vol5issue6/PartE/5-5-89-519.pdf

Karthik, S, Reddy, M.S. & Yashaswini, G. (2021). Climate change and its potential impacts on insect-plant interactions [Online First]. IntechOpen, https://doi.org/10.5772/intechopen.98203. Available from: https://www.intechopen.com/online-first/77171.

Kavitha, K., & Reddy, K. D. (2014). Tritrophy-A new dimension in IPM-A review. Journal of Progressive Agriculture, 5(2): 99–111. https://doi.org/10.5958/0976-0741.2014.00907.6.

Khan, M.M.H. (2019). Effect of temperature and relative humidity on the population dynamics of brinjal and tomato infesting whitefly, *Bemisia tabaci*. Jahangirnagar University Journal of Biological Sciences, 8(1), 83–86. https://doi.org/10.3329/jujbs.v8i1.42471

Koul, O., Dhaliwal, G.S. & Cuperus, G.W. (eds.) (2004). Integrated Pest Management: potential, constraints, and challenges. CABI Publishing, UK, 342 pp. Available at https://tripleis.org/wp-content/uploads/2019/12/integrated-pest-management-cabi-publishing.pdf#page=67.

Koul, O., Walia, S. & Dhaliwal, G.S. (2008). Essential Oils as Green Pesticides: Potential and Constraints. Biopesticides International, 4(1): 63–84. Available at http://www.connectjournals.com/achivestoc2.php?fulltext=105501H_63-84.pdf&&bookmark=CJ-023217&&issue_id=01&&yaer=2008.

Laws, A.N. (2017). Climate change effects on predator-prey interactions. Current Opinion in Insect Science, 23: 28–34. https://doi.org/10.1016/j.cois.2017.06.010.

Li, D. & Jackson, R.R. (1996). How temperature affects development and reproduction in spiders: A review. Journal of Thermal Biology, 21(4): 245–274. https://doi.org/10.1016/0306-4565(96)00009-5.

Mahakittikun, V., Boitano, J., Ninsanit, P., Wangapai, T. & Ralukruedej K. (2011). Effects of high and low temperatures on development time and mortality of house dust mite eggs. Experimental & applied acarology, 55(4): 339–347. https://doi.org/10.1007/s10493-011-9480-2.

McDonald, J.R., Bale, J.S. & Walters, K.F.A. (1998). Effect of temperature on development of the Western Flower Thrips, *Frankliniella occidentalis* (Thysanoptera: Thripidae). European Journal of Entomology, 95: 301–306. Available at https://www.eje.cz/pdfs/eje/1998/02/14.pdf

Melo, J.W.S., Lima, D.B., Staudacher, H., Silva, F.R., Gondim Jr, M.G.C. & Sabelis, M.W. (2015). Evidence of *Amblyseius largoensis* and *Euseius alatus* as biological control agent of *Aceria guerreronis*. Experimental & Applied Acarology, 67: 411–421. https://doi.org/10.1007/s10493-015-9963-7.

Messelink, G.J., Bloemhard, C.M.J., Sabelis, M.W. & Janssen, A. (2013). Biological control of aphids in the presence of thrips and their enemies. BioControl 58, 45–55. https://doi.org/10.1007/s10526-012-9462-2.

Messelink, G.J., Van Steenpaal, S.E.F. & Ramakers, P.M.J. (2006). Evaluation of phytoseiid predators for control of western flower thrips on greenhouse cucumber. BioControl, 51(6): 753–768. https://doi.org/10.1007/s10526-006-9013-9.

Momen, F.M. & Abdelkhader, M.M. (2010). Fungi as food source for the generalist predator *Neoseiulus barkeri* (Hughes) (Acari: Phytoseiidae). Acta Phytopathologica et Entomologica Hungarica, 45(2): 401–409. https://doi.org/10.1556/aphyt.45.2010.2.18.

Momen, F.M. & Abdel-Khalek, A.A. (2008). Effect of the tomato rust mite *Aculops lycopersici* (Acari: Eriophyidae) on the development and reproduction of three predatory phytoseiid mites. International Journal of Tropical Insect Science, 28(1): 53–57. https://doi.org/10.1017/S1742758408942594.

Momen, F.M. & Abdel-Khalek, A.A. (2009 a). Cannibalism and intraguild predation in the phytoseiid mites *Typhlodromips swirskii, Euseius scutalis* and *Typhlodromus athiasae* (Acari: Phytoseiidae). Acarina, 17(2): 223–229. Available at https://acarina.utmn.ru/journal/release-archive/2009/17-2/172472/

Momen, F.M. & Abdel-Khalek, A.A. (2009 b). Juvenile survival and development of *Typhlodromips swirskii, Euseius scutalis* and *Typhlodromus athiasae* (Acari: Phytoseiidae) feeding on con-and heterospecific immatures. Acta Phytopathologica et Entomologica Hungarica, 44(1): 167–176. https://doi.org/10.1556/aphyt.44.2009.1.18

Momen, F.M. & Abdel-Khalek, A.A. (2021). Intraguild predation in three generalist predatory mites of the family Phytoseiidae (Acari: Phytoseiidae). Egyptian Journal of Biological Pest Control, 31: 8. https://doi.org/10.1186/s41938-020-00355-5

Momen, F.M. & El-Borolossy, M. (2010). Juvenile survival and development in three Phytoseiid species (Acari: Phytoseiidae) feeding on con-and heterospecific immatures. Acta Phytopathologica et Entomologica Hungarica, 45(2): 349–357. https://doi.org/10.1556/aphyt.45.2010.2.12

Momen, F.M. (2010). Intra-and interspecific predation by *Neoseiulus barkeri* and *Typhlodromus negevi* (Acari: Phytoseiidae) on different life stages: Predation rates and effects on reproduction and juvenile development. Acarina, 18(1): 81–88. Available at https://acarina.utmn.ru/journal/release-archive/2010/18-1/172486/

Momen, F.M., Hassan, M.F. & Lamlom, M. (2020). Evaluation of two factitious preys for rearing *Neoseiulus barkeri* (Acari: Phytoseiidae). International Journal of Acarology, 46(6): 387–393. https://doi.org/10.1080/01647954.2020.1804998.

Momen, F.M., Hussein, H. & Reda, A. (2013). Intra-guild vs extra-guild prey: Effect on development, predation and preference of *Typhlodromus negevi* Swirski and Amitai and *Typhlodromips swirskii* (Athias-Henriot) (Acari: Phytoseiidae). Acta Phytopathologica et Entomologica Hungarica, 48(1): 95–106. https://doi.org/10.1556/aphyt.48.2013.1.9

Momen, F.M., Metwally, A.M., Nasr, A.K., Abdallah, A.A. & Saleh, K.M. (2014). Life history of *Proprioseiopsis badri* feeding on four eriophyid mite species (Acari: Phytoseiidae and Eriophyidae). Phytoparasitica, 42(1): 23–30. https://doi.org/10.1007/s12600-013-0333-x.

Momen, F.M., Rasmy, A.H., Zaher, M.A., Nawar, M.S. & Abou-Elella, G.M. (2004). Dietary effect on the development, reproduction and sex-ratio of the predatory mite *Amblyseius denmarkeri* Zaher & El-Borolossy (Acari: Phytoseiidae). International Journal of Tropical Insect Science, 24(2): 192–195. https://doi.org/10.1079/IJT200414.

Napiórkowska, T., Templin, J., Grodzicki, P. & Kobak, J. (2021). Thermal preferences of two spider species: an orb-web weaver and a synanthropic funnel-web weaver. The European Zoological Journal, 88(1): 824–836. https://doi.org/10.1080/24750263.2021.1950223.

Nomikou, M., Janssen, A., Schraag, R. & Sabelis, M.W. (2001). Phytoseiid predators as potential biological control agents for *Bemisia tabaci*. Experimental & Applied Acarology, 25: 270–290.

Nomikou, M., Janssen, A., Schraag, R. & Sabelis, M.W. (2002). Phytoseiid predators suppress populations of *Bemisia tabaci* on cucumber plants with alternative food. Experimental & Applied Acarology, 27: 57–68.

Nomikou, M., Janssen, A. & Sabelis, M.W. (2003). Phytoseiid predator of whitefly feeds on plant tissue. Experimental & Applied Acarology, 31: 27–36. https://doi.org/10.1023/B:APPA.0000005150.33813.04

Parra, J.R.P. (2014). Biological control in Brazil: an overview. Scientia Agricola, 71(5): 420–429. https://doi.org/10.1590/0103-9016-2014-0167

Praslicka, J. & Huszár, J. (2004): Influence of temperature and host plants on the development and fecundity of the spider mite *Tetranychus urticae* (Acarina: Tetranychidae). Plant Protection Science, 40(4): 141–144. Available at https://www.agriculturejournals.cz/publicFiles/20268.pdf

Regnault-Roger, C., Hamraoui, A., Holeman, M., Theron, E. & Pinel, R. (1993) Insecticidal effect of essential oils from Mediterranean plants upon *Acanthoscelides obtectus* Say (Coleoptera, Bruchidae), a pest of kidney bean *(Phaseolus vulgaris* L.). Journal of Chemical Ecology, 19: 1233–1244. https://doi.org/10.1007/BF00987383.

Sanad, A.S. & Hassan, G.M. (2019). Controlling the western flower thrips, *Frankliniella occidentalis* (Pergande) (Thysanoptera: Thripidae) by releasing the predatory phytoseiid mites and pesticides on pepper in a greenhouse. Egyptian Journal of Biological Pest Control, 29(1): 95. https://doi.org/10.1186/s41938-019-0186-9.

Schuldiner-Harpaz, T. & Coll, M. (2013). Effects of Global Warming on Predatory Bugs Supported by Data Across Geographic and Seasonal Climatic Gradients. PLoS ONE, 8(6): e66622. https://doi.org/10.1371/journal.pone.0066622

Shelton, A. & Badenes-Pérez, F. (2006). Concepts and applications of trap cropping in pest management. Annual Review of Entomology, 51(1): 285–308. https://doi.org/10.1146/annurev.ento.51.110104.150959.

Shijiang, P. (1983). Biological Control - One of the fine traditions of ancient Chinese agricultural techniques. Scientia Agricultura Sinica, 1: 92–98.

Shrestha, S. (2019). Effects of climate change in agricultural insect pest. Acta Scientific Agriculture, 3(12): 74–80. https://doi.org/10.31080/ASAG.2019.03.0727.

Sinha, K.K., Choudhary, A.K. & Kumari, P. (2016). Entomopathogenic fungi. pp: 475–505. https://doi.org/10.1016/B978-0-12-803265-7.00015-4. In: Omkar (ed.). Ecofriendly pest management for food security. Academic Press, Elsevier. https://doi.org/10.1016/C2014-0-04228-1

Skendžić, S.; Zovko, M.; Živković, I.P.; Lešić, V.; Lemić, D. (2021). The impact of climate change on agricultural insect pests. Insects, 12: 440. https://doi.org/10.3390/insects12050440

Smagghe, G. & Diaz, I. (eds) (2012). Arthropod-Plant Interactions. Progress in Biological Control, vol 14. Springer, Dordrecht. https://doi.org/10.1007/978-94-007-3873-7.

Soloneski, S. & Larramendy, M. (Eds.) (2013). Weed and pest control - conventional and new challenges. IntechOpen, Available from https://www.intechopen.com/chapters/42758

Teich, Y. (1966). Mites of the family of Phytoseiidae as predators of the tobacco whitefly, *Bemisia tabaci* (Gennadius). Israel Journal of Agricultural Research, 16: 141–142.

Tixier, M.-S. (2018). Predatory mites (Acari: Phytoseiidae) in Agro-ecosystems and conservation biological control: A Review and explorative approach for forecasting plant-predatory mite interactions and mite dispersal. Frontiers in Ecology and Evolution, 6: 192. https://doi.org/10.3389/fevo.2018.00192

Urbaneja-Bernat, P. & Jaques, J.A. (2022). Can pollen provision mitigate competition interactions between three phytoseiid predators of *Tetranychus urticae* under future climate change conditions? Biological Control, 165: 104789. https://doi.org/10.1016/j.biocontrol.2021.104789.

Van Rijn, P.C. & Tanigoshi, L.K. (1999). Pollen as Food for the Predatory Mites *Iphiseius Degenerans and Neoseiulus Cucumeris* (Acari: Phytoseiidae): Dietary Range and Life History. Experimental & Applied Acarology, 23, 785–802. https://doi.org/10.1023/A:1006227704122.

Verkerk, R. H. (2004). Manipulation of tritrophic interactions for IPM. pp: 55-72. In: O. Koul, G.S. Dhaliwal & G.W. Cuperus (eds.). Integrated Pest Management: potential, constraints, and challenges. CABI Publishing, UK, 342 pp. Available at https://tripleis.org/wp-content/uploads/2019/12/integrated-pest-management-cabi-publishing.pdf#page=67.

White, J. & Liburd, O. (2005). Effects of soil moisture and temperature on reproduction and development of Two spotted Spider Mite (Acari: Tetranychidae) in Strawberries. Journal of Economic Entomology, 98(1): 154–8. https://doi.org/10.1603/0022-0493-98.1.154.

Wraight, S. P., Ugine, T. A., Ramos, M. E., and Sanderson, J. P. (2016). Efficacy of spray applications of entomopathogenic fungi against western flower thrips infesting greenhouse impatiens under variable moisture conditions. Biological Control, 97: 31–47. https://doi.org/10.1016/j.biocontrol.2016.02.016.

Xin, T.-R., & Zhang, Z.-Q. (2021). Suitability of pollen as an alternative food source for different developmental stages of *Amblyseius herbicolus* (Chant) (Acari: Phytoseiidae) to facilitate predation on whitefly eggs. Acarologia, 61(4): 790–801. https://doi.org/10.24349/bIV1-2heN

Zemek, R. & Prenerová, E. (1997). Powdery mildew (Ascomycotina: Erysiphales) - an alternative food for the predatory mite *Typhlodromus pyri* Scheuten (Acari: Phytoseiidae). Experimental & Applied Acarology. 21: 405–414. https://doi.org/10.1023/A:1018427812075

Zidan, I.M. (2021). Biodiversity and natural control of mites associated with medicinal plants in organic farming systems. PhD Thesis, Faculty of Agriculture, Cairo University, 304 pp.

Zou, Z., Xi, J., Liu, G., Song, S., Xin, T., & Xia, B. (2018). Effect of temperature on development and reproduction of the carmine spider mite, *Tetranychus cinnabarinus* (Acari: Tetranychiae), fed on cassava leaves. Experimental & applied acarology, 74(4): 383–394. https://doi.org/10.1007/s10493-018-0241-3

