## Supplementary file 1 for "Predatory mites, a green pesticide, and an Entomopathogenic compound: A proposed IPM tactic based on pest species diversity indices and population dynamics"

| **Table (1). Weather data in Om Saber location during season 2017.** | | | | |
| --- | --- | --- | --- | --- |
| **Sample date** | **Optimum Temperature**  **°C (Max. – Min.)** | | **Relative Humidity**  **R.H.% (Max. – Min.)** | |
| **02 Apr, 2017** | 19.00 | (23 - 15) | 60.50 | (83 - 38) |
| **09 Apr, 2017** | 21.00 | (25 - 17) | 63.50 | (83 - 44) |
| **16 Apr, 2017** | 21.50 | (27 - 16) | 54.00 | (88 - 20) |
| **23 Apr, 2017** | 21.00 | (25 - 17) | 52.00 | (73 - 31) |
| **30 Apr, 2017** | 25.50 | (32 - 19) | 53.50 | (88 - 19) |
| **07 May, 2017** | 26.00 | (32 - 20) | 51.50 | (83 - 20) |
| **14 May, 2017** | 30.50 | (38 - 23) | 34.00 | (51 - 17) |
| **21 May, 2017** | 25.00 | (30 - 20) | 53.50 | (83 - 24) |
| **28 May, 2017** | 28.50 | (35 - 22) | 47.50 | (78 - 17) |
| **04 Jun, 2017** | 29.00 | (35 - 23) | 53.00 | (83 - 23) |
| **11 Jun, 2017** | 27.50 | (33 - 22) | 61.00 | (88 - 34) |
| **18 Jun, 2017** | 31.50 | (38 - 25) | 40.50 | (65 - 16) |
| **25 Jun, 2017** | 30.00 | (36 - 24) | 55.50 | (89 - 22) |
| **02 Jul, 2017** | 28.50 | (33 - 24) | 54.00 | (74 - 34) |
| **09 Jul, 2017** | 32.00 | (37 - 27) | 54.00 | (74 - 34) |
| **16 July, 2017** | 31.00 | (37 - 25) | 61.00 | (89 - 33) |
| **23 Jul, 2017** | 32.00 | (39 - 25) | 59.50 | (89 - 30) |
| **30 July, 2017** | 30.50 | (36 - 25) | 54.50 | (79 - 30) |
| **06 Aug, 2017** | 31.50 | (36 - 27) | 54.00 | (74 - 34) |
| **13 Aug, 2017** | 31.00 | (36 - 26) | 56.00 | (84 - 28) |
| **20 Aug, 2017** | 31.00 | (36 - 26) | 55.50 | (79 - 32) |

| **Table (2). Mean number *T. urticae* on *B. napus*, *B. oleracea*, *M. spicata* and *M. longifolia* leaves at Om Saber location after releasing *P. persimilis.*** | | | | | | | | |
| --- | --- | --- | --- | --- | --- | --- | --- | --- |
|  | ***B. napus*** | **check** | ***B. oleracea*** | **check** | ***M. spicata*** | **check** | ***M. longifolia*** | **check** |
| **Precount** | **47.41** | **63.22** | **59.48** | **59.90** | **3.93** | **4.02** | **4.80** | **5.53** |
| **09 Apr, 2017** | 43.28 | 67.55 | 50.28 | 64.37 | 3.15 | 4.37 | 3.68 | 6.22 |
| **16 Apr, 2017** | 36.66 | 73.38 | 45.05 | 70.04 | 2.54 | 4.44 | 3.05 | 6.66 |
| **23 Apr, 2017** | 31.75 | 79.50 | 39.83 | 74.73 | 2.13 | 4.99 | 1.86 | 6.86 |
| **30 Apr, 2017** | 27.56 | 86.28 | 36.89 | 80.60 | 1.14 | 5.65 | 1.21 | 7.15 |
| **07 May, 2017** | 24.60 | 94.34 | 33.10 | 86.83 | 0.93 | 6.00 | 0.53 | 7.76 |
| **14 May, 2017** | 17.47 | 104.34 | 23.94 | 90.70 | 0.44 | 6.37 | 0.21 | 8.57 |
| **21 May, 2017** | 12.77 | 109.31 | 16.65 | 93.25 | 0.08 | 6.94 | 0.08 | 9.40 |
| **28 May, 2017** | 6.98 | 119.24 | 9.97 | 96.33 | 0.00 | 7.21 | 0.00 | 9.99 |
| **04 Jun, 2017** | 4.01 | 125.41 | 5.25 | 102.72 | 0.00 | 7.53 | 0.00 | 10.35 |
| **11 Jun, 2017** | 1.01 | 132.64 | 2.00 | 108.71 | 0.00 | 7.70 | 0.00 | 11.50 |
| **18 Jun, 2017** | 0.10 | 144.64 | 0.87 | 111.60 | 0.00 | 7.73 | 0.00 | 12.03 |
| **25 Jun, 2017** | 0.00 | 149.28 | 0.05 | 118.01 | 0.00 | 7.84 | 0.00 | 13.41 |
| **02 Jul, 2017** | 0.00 | 163.67 | 0.00 | 120.93 | 0.00 | 8.00 | 0.00 | 15.09 |
| **Mean over all** | **18.11^b^** | **108.06** | **23.10^a^** | **91.34** | **1.02^c^** | **6.34** | **1.10^c^** | **9.32** |
| ***P*_0.05_** | **0.000** |  | **0.000** |  | **0.000** |  | **0.000** |  |

**Supplementary file 1 (IPM data)**

| **Table (3). Mean number of *T. urticae* on Brassicaceae and Lamiaceae plants at Om Saber location after releasing *A. swirskii.*** | | | | | | | | |
| --- | --- | --- | --- | --- | --- | --- | --- | --- |
|  | ***B. napus*** | **check** | ***B. oleracea*** | **check** | ***M. spicata*** | **check** | ***M. longifolia*** | **check** |
| **Precount** | **54.28** | **63.22** | **57.11** | **59.90** | **3.85** | **4.02** | **4.76** | **5.53** |
| **09 Apr, 2017** | 50.20 | 67.55 | 53.49 | 64.37 | 3.68 | 4.37 | 4.50 | 6.22 |
| **16 Apr, 2017** | 46.92 | 73.38 | 49.94 | 70.04 | 3.38 | 4.44 | 4.23 | 6.66 |
| **23 Apr, 2017** | 42.67 | 79.50 | 44.08 | 74.73 | 2.95 | 4.99 | 3.76 | 6.86 |
| **30 Apr, 2017** | 41.09 | 86.28 | 41.03 | 80.60 | 2.59 | 5.65 | 3.21 | 7.15 |
| **07 May, 2017** | 37.13 | 94.34 | 38.25 | 86.83 | 2.08 | 6.00 | 2.67 | 7.76 |
| **14 May, 2017** | 31.83 | 104.34 | 29.69 | 90.70 | 1.50 | 6.37 | 2.05 | 8.57 |
| **21 May, 2017** | 27.18 | 109.31 | 24.91 | 93.25 | 1.05 | 6.94 | 1.78 | 9.40 |
| **28 May, 2017** | 25.34 | 119.24 | 20.18 | 96.33 | 0.88 | 7.21 | 1.48 | 9.99 |
| **04 Jun, 2017** | 24.41 | 125.41 | 16.51 | 102.72 | 0.62 | 7.53 | 1.21 | 10.35 |
| **11 Jun, 2017** | 23.54 | 132.64 | 16.01 | 108.71 | 0.45 | 7.70 | 1.10 | 11.50 |
| **18 Jun, 2017** | 24.55 | 144.64 | 16.30 | 111.60 | 0.30 | 7.73 | 1.00 | 12.03 |
| **25 Jun, 2017** | 24.99 | 149.28 | 17.85 | 118.01 | 0.22 | 7.84 | 0.98 | 13.41 |
| **02 Jul, 2017** | 26.88 | 163.67 | 14.90 | 120.93 | 0.16 | 8.00 | 1.00 | 15.09 |
| **09 Jul, 2017** | 16.40 | 171.49 | 13.16 | 124.78 | 0.10 | 8.91 | 0.88 | 15.52 |
| **16 Jul, 2017** | 13.83 | 188.28 | 12.48 | 117.93 | 0.07 | 8.89 | 0.74 | 15.21 |
| **23 Jul, 2017** | 10.25 | 201.31 | 11.55 | 100.16 | 0.02 | 8.67 | 0.61 | 14.04 |
| **30 July, 2017** | 9.42 | 205.63 | 11.00 | 95.57 | 0.01 | 8.45 | 0.49 | 10.77 |
| **06 Aug, 2017** | 6.12 | 196.46 | 10.89 | 88.27 | 0.00 | 6.48 | 0.40 | 9.27 |
| **13 Aug, 2017** | 5.26 | 194.11 | 10.43 | 83.67 | 0.00 | 6.37 | 0.35 | 7.77 |
| **20 Aug, 2017** | 3.50 | 192.43 | 10.01 | 79.53 | 0.00 | 6.23 | 0.10 | 7.27 |
| **Mean over all** | **25.99^a^** | **136.31** | **24.75^a^** | **93.74** | **1.14^b^** | **6.80** | **1.78^b^** | **10.02** |
| ***P*_0.05_** | **0.000** |  | **0.000** |  | **0.000** |  | **0.000** |  |

| **Table (4). Mean number *T. urticae* after releasing *C. negevi.*** | | | | | | | | |
| --- | --- | --- | --- | --- | --- | --- | --- | --- |
|  | ***B. napus*** | **check** | ***B. oleracea*** | **check** | ***M. spicata*** | **check** | ***M. longifolia*** | **check** |
| **Precount** | **52.70** | **63.22** | **56.90** | **59.90** | **3.85** | **4.02** | **4.98** | **5.53** |
| **09 Apr, 2017** | 50.43 | 67.55 | 49.60 | 64.37 | 3.68 | 4.37 | 4.61 | 6.22 |
| **16 Apr, 2017** | 48.84 | 73.38 | 46.15 | 70.04 | 3.46 | 4.44 | 4.48 | 6.66 |
| **23 Apr, 2017** | 49.26 | 79.50 | 47.58 | 74.73 | 3.12 | 4.99 | 4.30 | 6.86 |
| **30 Apr, 2017** | 49.04 | 86.28 | 47.60 | 80.60 | 2.93 | 5.65 | 4.16 | 7.15 |
| **07 May, 2017** | 48.07 | 94.34 | 48.50 | 86.83 | 2.70 | 6.00 | 3.92 | 7.76 |
| **14 May, 2017** | 43.56 | 104.34 | 40.69 | 90.70 | 2.53 | 6.37 | 3.64 | 8.57 |
| **21 May, 2017** | 40.69 | 109.31 | 38.00 | 93.25 | 2.18 | 6.94 | 3.46 | 9.40 |
| **28 May, 2017** | 37.70 | 119.24 | 34.88 | 96.33 | 1.90 | 7.21 | 3.30 | 9.99 |
| **11 Jun, 2017** | 37.98 | 132.64 | 31.70 | 108.71 | 1.55 | 7.70 | 2.99 | 11.50 |
| **18 Jun, 2017** | 38.11 | 144.64 | 30.67 | 111.60 | 1.49 | 7.73 | 2.75 | 12.03 |
| **25 Jun, 2017** | 40.77 | 149.28 | 30.89 | 118.01 | 1.47 | 7.84 | 2.59 | 13.41 |
| **02 Jul, 2017** | 42.71 | 163.67 | 26.52 | 120.93 | 1.46 | 8.00 | 2.20 | 15.09 |
| **09 Jul, 2017** | 37.83 | 171.49 | 25.25 | 124.78 | 1.31 | 8.91 | 1.99 | 15.52 |
| **16 Jul, 2017** | 34.74 | 188.28 | 23.10 | 117.93 | 1.29 | 8.89 | 1.85 | 15.21 |
| **23 Jul, 2017** | 30.45 | 201.31 | 22.20 | 100.16 | 1.27 | 8.67 | 1.81 | 14.04 |
| **30 Jul, 2017** | 29.37 | 205.63 | 21.67 | 95.57 | 1.25 | 8.45 | 1.74 | 10.77 |
| **06 Aug, 2017** | 31.23 | 196.46 | 22.57 | 88.27 | 1.20 | 6.48 | 1.65 | 9.27 |
| **13 Aug, 2017** | 35.99 | 194.11 | 23.00 | 83.67 | 1.18 | 6.37 | 1.49 | 7.77 |
| **20 Aug, 2017** | 39.15 | 192.43 | 25.33 | 79.53 | 1.11 | 6.23 | 1.38 | 7.27 |
| **Mean over all** | **40.72^a^** | **136.31** | **34.64^b^** | **93.74** | **2.03^c^** | **6.80** | **2.97^c^** | **10.02** |
| ***P*_0.05_** | **0.000** |  | **0.000** |  | **0.000** |  | **0.000** |  |

| **Table (5). Mean number *T. urticae* after releasing Bio-Magic.** | | | | | | | | |
| --- | --- | --- | --- | --- | --- | --- | --- | --- |
|  | ***B. napus*** | **check** | ***B. oleracea*** | **check** | ***M. spicata*** | **check** | ***M. longifolia*** | **check** |
| **Precount** | **36.68** | **63.22** | **28.42** | **59.90** | **5.34** | **4.02** | **4.34** | **5.53** |
| **09 Apr, 2017** | 26.45 | 67.55 | 18.52 | 64.37 | 4.41 | 4.37 | 2.40 | 6.22 |
| **16 Apr, 2017** | 23.50 | 73.38 | 20.10 | 70.04 | 4.35 | 4.44 | 2.36 | 6.66 |
| **23 Apr, 2017** | 26.50 | 79.50 | 25.98 | 74.73 | 4.39 | 4.99 | 4.00 | 6.86 |
| **30 Apr, 2017** | 35.20 | 86.28 | 34.22 | 80.60 | 4.23 | 5.65 | 5.18 | 7.15 |
| **07 May, 2017** | 45.05 | 94.34 | 44.84 | 86.83 | 4.52 | 6.00 | 6.20 | 7.76 |
| **14 May, 2017** | 24.00 | 104.34 | 37.12 | 90.70 | 3.90 | 6.37 | 2.00 | 8.57 |
| **21 May, 2017** | 33.30 | 109.31 | 36.38 | 93.25 | 3.80 | 6.94 | 1.80 | 9.40 |
| **28 May, 2017** | 50.55 | 119.24 | 40.36 | 96.33 | 3.60 | 7.21 | 1.94 | 9.99 |
| **04 Jun, 2017** | 64.55 | 125.41 | 54.41 | 102.72 | 3.67 | 7.53 | 2.20 | 10.35 |
| **11 Jun, 2017** | 69.25 | 132.64 | 61.28 | 108.71 | 4.15 | 7.70 | 4.86 | 11.50 |
| **18 Jun, 2017** | 84.65 | 144.64 | 67.72 | 111.60 | 4.74 | 7.73 | 7.74 | 12.03 |
| **25 Jun, 2017** | 36.19 | 149.28 | 27.91 | 118.01 | 5.30 | 7.84 | 9.20 | 13.41 |
| **02 Jul, 2017** | 40.00 | 163.67 | 27.84 | 120.93 | 5.77 | 8.00 | 10.64 | 15.09 |
| **09 Jul, 2017** | 50.68 | 171.49 | 38.57 | 124.78 | 6.09 | 8.91 | 5.94 | 15.52 |
| **16 July, 2017** | 56.33 | 188.28 | 46.61 | 117.93 | 5.76 | 8.89 | 5.50 | 15.21 |
| **23 Jul, 2017** | 62.35 | 201.31 | 53.10 | 100.16 | 5.92 | 8.67 | 6.00 | 14.04 |
| **30 Jul, 2017** | 69.77 | 205.63 | 62.37 | 95.57 | 5.91 | 8.45 | 6.22 | 10.77 |
| **Mean over all** | **46.39^a^** | **126.64** | **40.32^a^** | **95.40** | **4.77^b^** | **6.87** | **4.92^b^** | **10.34** |
| ***P*_0.05_** | **0.000** |  | **0.000** |  | **0.000** |  | **0.000** |  |

| **Table (6). Mean number *T. urticae* after application of Egyxide.** | | | | | | | | |
| --- | --- | --- | --- | --- | --- | --- | --- | --- |
|  | ***B. napus*** | **check** | ***B. oleracea*** | **check** | ***M. spicata*** | **check** | ***M. longifolia*** | **check** |
| **Precount** | **36.30** | **63.22** | **28.21** | **59.90** | **5.57** | **4.02** | **5.42** | **5.53** |
| **09 Apr, 2017** | 26.16 | 67.55 | 20.46 | 64.37 | 2.95 | 4.37 | 2.00 | 6.22 |
| **16 Apr, 2017** | 20.09 | 73.38 | 22.20 | 70.04 | 2.88 | 4.44 | 2.02 | 6.66 |
| **23 Apr, 2017** | 34.25 | 79.50 | 27.49 | 74.73 | 3.33 | 4.99 | 2.42 | 6.86 |
| **30 Apr, 2017** | 44.30 | 86.28 | 39.08 | 80.60 | 3.83 | 5.65 | 4.60 | 7.15 |
| **07 May, 2017** | 50.47 | 94.34 | 47.58 | 86.83 | 4.08 | 6.00 | 5.78 | 7.76 |
| **14 May, 2017** | 25.45 | 104.34 | 38.51 | 90.70 | 2.89 | 6.37 | 1.00 | 8.57 |
| **21 May, 2017** | 31.70 | 109.31 | 38.10 | 93.25 | 2.75 | 6.94 | 0.94 | 9.40 |
| **28 May, 2017** | 48.02 | 119.24 | 41.06 | 96.33 | 3.14 | 7.21 | 1.46 | 9.99 |
| **04 Jun, 2017** | 57.01 | 125.41 | 65.46 | 102.72 | 3.86 | 7.53 | 2.76 | 10.35 |
| **11 Jun, 2017** | 70.55 | 132.64 | 70.95 | 108.71 | 4.32 | 7.70 | 4.54 | 11.50 |
| **18 Jun, 2017** | 85.27 | 144.64 | 80.44 | 111.60 | 5.28 | 7.73 | 6.24 | 12.03 |
| **25 Jun, 2017** | 34.02 | 149.28 | 44.15 | 118.01 | 5.93 | 7.84 | 7.80 | 13.41 |
| **02 Jul, 2017** | 40.71 | 163.67 | 44.82 | 120.93 | 6.29 | 8.00 | 9.74 | 15.09 |
| **09 Jul, 2017** | 60.87 | 171.49 | 54.06 | 124.78 | 3.75 | 8.91 | 4.00 | 15.52 |
| **16 July, 2017** | 66.55 | 188.28 | 61.25 | 117.93 | 4.25 | 8.89 | 4.20 | 15.21 |
| **23 Jul, 2017** | 72.89 | 201.31 | 71.78 | 100.16 | 4.77 | 8.67 | 6.10 | 14.04 |
| **30 Jul, 2017** | 80.30 | 205.63 | 83.05 | 95.57 | 5.48 | 8.45 | 6.40 | 10.77 |
| **Mean over all** | **49.16** | **126.64** | **48.81** | **95.40** | **4.19** | **6.87** | **4.30** | **10.34** |
| ***P*_0.05_** | **0.000** |  | **0.000** |  | **0.000** |  | **0.000** |  |

| **Table (7). Reduction percentage of *T. urticae* on Russian kale *Brassica napus var. pabularia* leaves.** | | | | | |
| --- | --- | --- | --- | --- | --- |
| **Sampling date** | ***P. persimilis*** | ***A. swirskii*** | ***C. negevi*** | **Bio-Magic** | **Egyxide** |
| **09 Apr, 2017** | 35.05 | 13.66 | 10.34 | 31.59 | 54.65 |
| **16 Apr, 2017** | 47.70 | 26.85 | 17.25 | 44.33 | 53.03 |
| **23 Apr, 2017** | 57.83 | 31.96 | 16.61 | 44.75 | 28.15 |
| **30 Apr, 2017** | 60.00 | 39.96 | 20.50 | 26.22 | 6.78 |
| **14 May, 2017** | 68.34 | 14.67 | 13.21 | 64.41 | 63.12 |
| **21 May, 2017** | 82.62 | 25.81 | 21.72 | 61.17 | 40.73 |
| **28 May, 2017** | 92.51 | 38.58 | 32.50 | 46.69 | 26.13 |
| **04 Jun, 2017** | 98.03 | 51.00 | 39.50 | 43.27 | 20.08 |
| **11 Jun, 2017** | 100.00 | 61.48 | 43.22 | 34.62 | 7.16 |
| **18 Jun, 2017** | - | 68.58 | 53.53 | 26.27 | 12.27 |
| **25 Jun, 2017** | - | 70.70 | 54.62 | 13.65 | 1.92 |
| **09 Jul, 2017** | - | 45.17 | 18.51 | 49.35 | 64.90 |
| **16 Jul, 2017** | - | 65.61 | 38.79 | 21.40 | 58.99 |
| **23 Jul, 2017** | - | 89.72 | 50.50 | -12.84 | 30.00 |
| **30 Jul, 2017** | - | 93.66 | 55.95 | -24.03 | 28.55 |
| **06 Aug, 2017** | - | 100.00 | 56.67 | -21.69 | 20.36 |
| **13 Aug, 2017** | - | - | 56.30 | -14.61 | 11.05 |
| **Mean** | **71.34^a^** | **52.34^b^** | **36.51^c^** | **22.89^d^** | **29.24^d^** |
| **L.S.D. at 0.05 = 26.31** | | | | | |

| **Table (8). Reduction percentage of the *T. urticae* on Italian kale *Brassica oleracea var palmifolia* leaves.** | | | | | |
| --- | --- | --- | --- | --- | --- |
| **Sampling date** | ***P. persimilis*** | ***A. swirskii*** | ***C. negevi*** | **Bio-Magic** | **Egyxide** |
| **09 Apr, 2017** | 27.30 | 14.05 | 22.91 | 37.75 | 43.66 |
| **16 Apr, 2017** | 39.70 | 28.03 | 40.13 | 43.32 | 48.37 |
| **23 Apr, 2017** | 49.37 | 41.15 | 45.24 | 40.06 | 47.40 |
| **30 Apr, 2017** | 58.23 | 51.85 | 52.09 | 39.38 | 44.93 |
| **14 May, 2017** | 33.68 | 37.59 | 9.75 | 39.68 | 41.27 |
| **21 May, 2017** | 62.58 | 55.85 | 20.02 | 42.71 | 41.83 |
| **28 May, 2017** | 85.24 | 73.17 | 32.72 | 39.06 | 34.90 |
| **04 Jun, 2017** | 94.57 | 88.48 | 35.58 | -19.99 | -79.36 |
| **11 Jun, 2017** | 100.00 | 95.76 | 42.66 | -40.06 | -86.03 |
| **18 Jun, 2017** | - | 100.00 | 46.86 | -45.28 | -95.21 |
| **25 Jun, 2017** | - | - | 50.07 | -45.34 | -97.83 |
| **09 Jul, 2017** | - | - | 6.86 | 61.84 | 54.13 |
| **16 Jul, 2017** | - | - | -0.30 | 56.26 | 49.54 |
| **23 Jul, 2017** | - | - | -7.61 | 38.66 | 29.96 |
| **30 Jul, 2017** | - | - | 8.01 | 25.59 | 15.79 |
| **06 Aug, 2017** | - | - | 12.43 | -1.82 | -15.93 |
| **13 Aug, 2017** | - | - | 15.57 | -44.94 | -61.05 |
| **Mean** | **61.19 a** | **58.59 a** | **24.70 b** | **11.56 c** | **3.20 d** |
| **L.S.D. at 0.05 = 39.52** | | | | | |

| **Table (9). Reduction percentage of *T. urticae* on baladi mint *Mentha spicata* leaves .** | | | | | |
| --- | --- | --- | --- | --- | --- |
| Sampling date | ***P. persimilis*** | ***A. swirskii*** | ***C. negevi*** | **Bio-Magic** | **Egyxide** |
| **09 Apr, 2017** | 27.17 | 13.03 | 18.40 | 57.96 | 77.93 |
| **16 Apr, 2017** | 93.72 | 24.40 | 27.64 | 57.27 | 79.16 |
| **23 Apr, 2017** | 95.19 | 40.72 | 44.42 | 54.03 | 71.45 |
| **30 Apr, 2017** | 99.42 | 58.15 | 58.22 | 58.09 | 67.05 |
| **14 May, 2017** | 100.00 | 45.59 | 15.85 | 61.16 | 82.09 |
| **21 May, 2017** | - | 94.70 | 54.18 | 67.93 | 86.64 |
| **28 May, 2017** | - | 100.00 | 76.50 | 71.28 | 73.73 |
| **04 Jun, 2017** | - | - | 93.81 | 59.18 | 36.95 |
| **11 Jun, 2017** | - | - | 100.00 | 46.41 | 15.03 |
| **18 Jun, 2017** | - | - | 100.00 | 32.50 | -34.59 |
| **25 Jun, 2017** | - | - | - | 14.70 | -40.30 |
| **09 Jul, 2017** | - | - | - | 10.27 | 85.07 |
| **16 Jul, 2017** | - | - | - | 19.25 | 58.63 |
| **23 Jul, 2017** | - | - | - | -9.34 | 44.31 |
| **30 Jul, 2017** | - | - | - | -17.55 | 19.51 |
| **06 Aug, 2017** | - | - | - | -22.00 | 13.71 |
| **13 Aug, 2017** | - | - | - | -36.55 | 9.15 |
| **Mean** | **83.10 a** | **53.80 b** | **54.34 b** | **26.90 d** | **41.38 c** |
| **L.S.D. at 0.05 = 43.10** | | | | | |

| **Table (10). Reduction percentage of *T. urticae* on Saudi mint *Mentha longifolia* leaves.** | | | | | |
| --- | --- | --- | --- | --- | --- |
| **Sampling date** | ***P. persimilis*** | ***A. swirskii*** | ***C. negevi*** | **Bio-Magic** | **Egyxide** |
| **09 Apr, 2017** | 36.18 | 10.70 | 12.51 | 48.32 | 65.51 |
| **16 Apr, 2017** | 48.03 | 25.76 | 23.84 | 53.66 | 68.24 |
| **23 Apr, 2017** | 74.33 | 41.64 | 31.04 | 23.41 | 62.90 |
| **30 Apr, 2017** | 89.31 | 55.80 | 36.00 | 1.90 | 30.24 |
| **14 May, 2017** | 100.00 | 27.08 | 24.98 | 72.00 | 84.98 |
| **21 May, 2017** | - | 53.13 | 39.72 | 77.32 | 87.29 |
| **28 May, 2017** | - | 75.00 | 47.62 | 76.30 | 80.86 |
| **04 Jun, 2017** | - | 91.80 | 53.98 | 73.55 | 64.40 |
| **11 Jun, 2017** | - | 96.25 | 66.44 | 51.01 | 50.91 |
| **18 Jun, 2017** | - | 100.00 | 75.00 | 23.38 | 33.74 |
| **25 Jun, 2017** | - | - | 82.66 | 22.82 | 29.81 |
| **09 Jul, 2017** | - | - | 56.61 | 46.16 | 60.40 |
| **16 Jul, 2017** | - | - | 100.00 | 47.60 | 56.29 |
| **23 Jul, 2017** | - | - | - | 39.18 | 32.46 |
| **30 Jul, 2017** | - | - | - | 28.81 | 19.99 |
| **06 Aug, 2017** | - | - | - | 7.91 | -3.69 |
| **13 Aug, 2017** | - | - | - | -18.94 | -37.25 |
| **Mean** | **69.57 a** | **57.72 b** | **50.03 b** | **35.09 c** | **40.30 c** |
| **L.S.D at 0.05 = 56.73** | | | | | |

| **Table (11).   Mean number of *T. tabaci* and *B. tabaci* after releasing *A. swirskii.*** | | | | | | | | |
| --- | --- | --- | --- | --- | --- | --- | --- | --- |
| ***Thrips tabaci*** | | | | | | | | |
|  | ***B. napus*** | **Check** | ***B. oleracea*** | **Check** | ***M. spicata*** | **Check** | ***M. longifolia*** | **Check** |
| **Precount** | **3.84** | **3.25** | **3.75** | **4.68** | **0.18** | **0.17** | **0.06** | **0.08** |
| **1 week** | 2.60 | 5.35 | 2.60 | 6.94 | 0.10 | 0.61 | 0.05 | 0.50 |
| **2 weeks** | 2.09 | 6.30 | 2.29 | 9.00 | 0.10 | 0.70 | 0.02 | 0.75 |
| **3 weeks** | 2.05 | 7.45 | 1.85 | 10.60 | 0.07 | 0.91 | 0.01 | 0.82 |
| **4 weeks** | 1.90 | 8.56 | 1.69 | 13.44 | 0.05 | 0.98 | - | 1.11 |
| **5 weeks** | 1.85 | 12.41 | 1.65 | 15.50 | 0.01 | 1.01 | - | 1.35 |
| **6 weeks** | 1.12 | 15.33 | 1.22 | 19.61 | - | 1.09 | - | 1.68 |
| **7 weeks** | 1.31 | 23.26 | 1.10 | 25.86 | - | 1.11 | - | 2.00 |
| **8 weeks** | 0.90 | 26.25 | 0.63 | 32.73 | - | 1.17 | - | 2.13 |
| **9 weeks** | 0.65 | 31.16 | 0.20 | 36.32 | - | 1.37 | - | 2.12 |
| **10 weeks** | 0.25 | 36.14 | - | 40.22 | - | 1.48 | - | 2.25 |
| **Mean over all** | **1.55** | **18.14** | **1.42** | **21.66** | **0.04** | **1.01** | **0.01** | **1.42** |
| **L.S.D at 0.05 = 14.30** | | | | | | | | |
| ***Bemisia tabaci*** | | | | | | | | |
|  | ***B. napus*** | **Check** | ***B. oleracea*** | **Check** | ***M. spicata*** | **Check** | ***M. longifolia*** | **Check** |
| **Precount** | **8.12** | **8.19** | **3.50** | **5.38** | **3.00** | **3.21** | **2.15** | **2.13** |
| **1 week** | 6.45 | 23.11 | 2.60 | 18.15 | 2.45 | 3.90 | 1.82 | 2.68 |
| **2 weeks** | 5.30 | 44.15 | 2.29 | 28.55 | 2.30 | 4.09 | 1.66 | 2.94 |
| **3 weeks** | 5.00 | 66.18 | 3.05 | 35.19 | 2.17 | 5.51 | 1.21 | 3.06 |
| **4 weeks** | 7.05 | 87.96 | 6.90 | 51.44 | 2.06 | 5.10 | 1.05 | 3.60 |
| **5 weeks** | 9.24 | 100.57 | 10.85 | 62.13 | 1.00 | 5.33 | 0.56 | 3.71 |
| **6 weeks** | 4.68 | 120.18 | 7.12 | 86.15 | 0.19 | 5.64 | 0.20 | 3.90 |
| **7 weeks** | 3.19 | 131.20 | 5.31 | 97.11 | 0.06 | 5.93 | 0.09 | 4.00 |
| **8 weeks** | 2.00 | 156.13 | 3.29 | 113.15 | - | 6.11 | - | 4.15 |
| **9 weeks** | 1.66 | 161.90 | 1.65 | 123.90 | - | 6.22 | - | 4.44 |
| **10 weeks** | 0.33 | 165.21 | 1.25 | 131.70 | - | 6.34 | - | 4.60 |
| **11 weeks** | 0.05 | 172.73 | 1.00 | 145.91 | - | 6.51 | - | 4.80 |
| **12 weeks** | - | 181.76 | 0.50 | 154.20 | - | 6.71 | - | 5.01 |
| **13 weeks** | - | 193.12 | 0.05 | 164.13 | - | 7.00 | - | 5.23 |
| **Mean over all** | **3.79^a^** | **115.17** | **3.53^a^** | **86.94** | **0.95^b^** | **5.54** | **0.62^b^** | **3.88** |
| **L.S.D at 0.05 = 10.32** | | | | | | | | |

| **Table (12). Mean number of *T. tabaci* and *B. tabaci* after releasing *C. negevi.*** | | | | | | | | |
| --- | --- | --- | --- | --- | --- | --- | --- | --- |
| ***Thrips tabaci*** | | | | | | | | |
|  | ***B. napus*** | **Check** | ***B. oleracea*** | **Check** | ***M. spicata*** | **Check** | ***M. longifolia*** | **Check** |
| **Precount** | **6.20** | **6.30** | **10.37** | **9.00** | **0.17** | **0.17** | **0.40** | **0.50** |
| **1 week** | 7.15 | 7.45 | 12.11 | 10.60 | 0.58 | 0.61 | 0.73 | 0.75 |
| **2 weeks** | 8.41 | 8.56 | 14.72 | 13.44 | 0.68 | 0.70 | 0.81 | 0.82 |
| **3 weeks** | 12.30 | 12.41 | 16.13 | 15.50 | 0.90 | 0.91 | 1.10 | 1.11 |
| **4 weeks** | 15.30 | 15.33 | 21.11 | 19.61 | 0.97 | 0.98 | 1.32 | 1.35 |
| **5 weeks** | 23.20 | 23.26 | 22.00 | 25.86 | 0.98 | 1.01 | 1.65 | 1.68 |
| **6 weeks** | 27.25 | 26.25 | 28.76 | 32.73 | 1.00 | 1.09 | 1.99 | 2.00 |
| **7 weeks** | 30.90 | 31.16 | 33.60 | 36.32 | 1.09 | 1.11 | 2.10 | 2.13 |
| **8 weeks** | 35.14 | 36.14 | 35.25 | 40.22 | 1.15 | 1.17 | 2.09 | 2.12 |
| **9 weeks** | 42.10 | 42.24 | 39.98 | 45.07 | 1.36 | 1.37 | 2.23 | 2.25 |
| **10 weeks** | 44.00 | 43.13 | 42.90 | 49.10 | 1.49 | 1.48 | 2.25 | 2.28 |
| **11 weeks** | 38.60 | 39.15 | 45.12 | 48.14 | 1.50 | 1.53 | 2.12 | 2.15 |
| **12 weeks** | 36.15 | 37.10 | 47.09 | 46.19 | 1.49 | 1.60 | 2.00 | 2.00 |
| **13 weeks** | 33.60 | 34.11 | 45.00 | 40.13 | 1.38 | 1.43 | 1.90 | 1.95 |
| **Mean over all** | **25.74^b^** | **25.90** | **29.58^a^** | **30.85** | **1.05^c^** | **1.08** | **1.62^c^** | **1.65** |
| **L.S.D at 0.05 =** | **15.20** | | | | | | | |
| ***Bemisia tabaci*** | | | | | | | | |
|  | ***B. napus*** | **Check** | ***B. oleracea*** | **Check** | ***M. spicata*** | **Check** | ***M. longifolia*** | **Check** |
| **Precount** | **5.00** | **8.19** | **6.12** | **5.38** | **2.37** | **3.21** | **2.18** | **2.13** |
| **1 week** | 4.60 | 23.11 | 6.13 | 18.15 | 1.80 | 3.90 | 2.10 | 2.68 |
| **2 weeks** | 3.47 | 44.15 | 8.11 | 28.55 | 1.68 | 4.09 | 2.00 | 2.94 |
| **3 weeks** | 3.88 | 66.18 | 9.00 | 35.19 | 1.50 | 5.51 | 1.84 | 3.06 |
| **4 weeks** | 3.99 | 87.96 | 10.06 | 51.44 | 1.49 | 5.10 | 1.66 | 3.60 |
| **5 weeks** | 4.84 | 100.57 | 11.10 | 62.13 | 1.12 | 5.33 | 1.42 | 3.71 |
| **6 weeks** | 2.50 | 120.18 | 7.12 | 86.15 | 1.00 | 5.64 | 1.33 | 3.90 |
| **7 weeks** | 1.59 | 131.20 | 6.61 | 97.11 | 0.90 | 5.93 | 1.12 | 4.00 |
| **8 weeks** | 1.03 | 156.13 | 4.90 | 113.15 | 0.24 | 6.11 | 1.02 | 4.15 |
| **9 weeks** | 0.75 | 161.90 | 3.22 | 123.90 | 0.10 | 6.22 | 0.70 | 4.44 |
| **10 weeks** | 0.55 | 165.21 | 2.02 | 131.70 | 0.00 | 6.34 | 0.43 | 4.60 |
| **11 weeks** | 0.10 | 172.73 | 0.50 | 145.91 | 0.00 | 6.51 | 0.13 | 4.80 |
| **12 weeks** | 0.00 | 181.76 | 0.01 | 154.20 | 0.00 | 6.71 | 0.04 | 5.01 |
| **13 weeks** | 0.00 | 193.12 | 0.00 | 164.13 | 0.00 | 7.00 | 0.00 | 5.23 |
| **Mean over all** | **2.31^b^** | **115.17** | **5.35^a^** | **86.94** | **0.87** | **5.54^a^** | **1.14^c^** | **3.88** |
| **L.S.D at 0.05 =** | **9.50** | | | | | | | |

| **Table (13).   Mean number of *T. tabaci* and *B. tabaci* after releasing Bio-Magic.** | | | | | | | | |
| --- | --- | --- | --- | --- | --- | --- | --- | --- |
| ***Thrips tabaci*** | | | | | | | | |
|  | ***B. napus*** | **Check** | ***B. oleracea*** | **Check** | ***M. spicata*** | **Check** | ***M. longifolia*** | **Check** |
| **Precount** | **1.08** | **6.30** | **1.51** | **9.00** | **0.62** | **0.17** | **0.80** | **0.50** |
| **1 week** | 1.90 | 7.45 | 1.80 | 10.60 | 0.39 | 0.61 | 0.49 | 0.75 |
| **2 weeks** | 1.00 | 8.56 | 1.16 | 13.44 | 0.35 | 0.70 | 0.47 | 0.82 |
| **3 weeks** | 0.95 | 12.41 | 1.10 | 15.50 | 0.42 | 0.91 | 0.50 | 1.11 |
| **4 weeks** | 3.23 | 15.33 | 2.95 | 19.61 | 0.84 | 0.98 | 1.00 | 1.35 |
| **5 weeks** | 6.40 | 23.26 | 7.15 | 25.86 | 1.10 | 1.01 | 1.19 | 1.68 |
| **6 weeks** | 3.00 | 26.25 | 4.66 | 32.73 | 0.50 | 1.09 | 0.76 | 2.00 |
| **7 weeks** | 2.64 | 31.16 | 4.50 | 36.32 | 0.48 | 1.11 | 0.73 | 2.13 |
| **8 weeks** | 4.15 | 36.14 | 7.90 | 40.22 | 0.65 | 1.17 | 0.89 | 2.12 |
| **9 weeks** | 9.72 | 42.24 | 13.81 | 45.07 | 0.84 | 1.37 | 1.00 | 2.25 |
| **10 weeks** | 5.86 | 43.13 | 9.21 | 49.10 | 1.10 | 1.48 | 0.85 | 2.28 |
| **11 weeks** | 6.00 | 39.15 | 10.75 | 48.14 | 1.25 | 1.53 | 0.92 | 2.15 |
| **12 weeks** | 6.10 | 37.10 | 11.00 | 46.19 | 1.32 | 1.60 | 1.10 | 2.00 |
| **Mean over all** | **4.00^b^** | **25.27** | **5.96^a^** | **30.14** | **0.76^c^** | **1.06** | **0.82^c^** | **1.63** |
| **L.S.D at 0.05 =** | **21.20** | | | | | | | |
| ***Bemisia tabaci*** | | | | | | | | |
|  | ***B. napus*** | **Check** | ***B. oleracea*** | **Check** | ***M. spicata*** | **Check** | ***M. longifolia*** | **Check** |
| **Precount** | **52.38** | **66.18** | **26.80** | **28.55** | **1.89** | **3.21** | **0.24** | **2.13** |
| **1 week** | 45.12 | 87.96 | 16.20 | 35.19 | 1.56 | 3.90 | 0.16 | 2.68 |
| **2 weeks** | 40.00 | 100.57 | 15.85 | 51.44 | 1.23 | 4.09 | 0.13 | 2.94 |
| **3 weeks** | 38.61 | 120.18 | 20.52 | 62.13 | 1.13 | 5.51 | 0.11 | 3.06 |
| **4 weeks** | 39.70 | 131.20 | 50.46 | 86.15 | 1.49 | 5.10 | 0.15 | 3.60 |
| **5 weeks** | 23.50 | 156.13 | 31.67 | 97.11 | 1.80 | 5.33 | 0.09 | 3.71 |
| **6 weeks** | 16.00 | 161.90 | 30.14 | 113.15 | 1.90 | 5.64 | 0.07 | 3.90 |
| **7 weeks** | 20.01 | 165.21 | 32.21 | 123.90 | 2.15 | 5.93 | 0.10 | 4.00 |
| **8 weeks** | 34.25 | 172.73 | 45.16 | 131.70 | 1.96 | 6.11 | 0.39 | 4.15 |
| **9 weeks** | 41.50 | 181.76 | 60.00 | 145.91 | 1.95 | 6.22 | 0.75 | 4.44 |
| **10 weeks** | 38.00 | 193.12 | 57.55 | 154.20 | 1.67 | 6.34 | 0.70 | 4.60 |
| **11 weeks** | 38.20 | 200.13 | 58.25 | 164.13 | 1.89 | 6.51 | 0.95 | 4.80 |
| **12 weeks** | 44.00 | 225.11 | 63.10 | 173.10 | 2.00 | 6.71 | 1.15 | 5.01 |
| **Mean over all** | **36.25^a^** | **150.94** | **39.07^a^** | **105.13** | **1.74^b^** | **5.43** | **0.38^c^** | **3.77** |
| **L.S.D at 0.05 =** | **30.25** | | | | | | | |

| **Table (14). Mean number of *T. tabaci* and *B. tabaci* after Egyxide application.** | | | | | | | | |
| --- | --- | --- | --- | --- | --- | --- | --- | --- |
| ***Thrips tabaci*** | | | | | | | | |
|  | ***B. napus*** | **Check** | ***B. oleracea*** | **Check** | ***M. spicata*** | **Check** | ***M. longifolia*** | **Check** |
| **Precount** | **2.05** | **6.30** | **1.79** | **9.00** |  |  |  |  |
| **1 week** | 0.90 | 7.45 | 0.99 | 10.60 |  |  |  |  |
| **2 weeks** | 0.93 | 8.56 | 1.05 | 13.44 |  |  |  |  |
| **3 weeks** | 2.66 | 12.41 | 2.25 | 15.50 | **0.72** | **0.91** | **0.81** | **1.11** |
| **4 weeks** | 6.00 | 15.33 | 6.18 | 19.61 | 0.60 | 0.98 | 0.78 | 1.35 |
| **5 weeks** | 3.15 | 23.26 | 3.54 | 25.86 | 0.44 | 1.01 | 0.52 | 1.68 |
| **6 weeks** | 3.00 | 26.25 | 3.29 | 32.73 | 0.42 | 1.09 | 0.50 | 2.00 |
| **7 weeks** | 5.90 | 31.16 | 8.44 | 36.32 | 0.49 | 1.11 | 0.83 | 2.13 |
| **8 weeks** | 10.18 | 36.14 | 12.60 | 40.22 | 0.79 | 1.17 | 1.10 | 2.12 |
| **9 weeks** | 6.12 | 42.24 | 8.99 | 45.07 | 1.00 | 1.37 | 1.05 | 2.25 |
| **10 weeks** | 6.10 | 43.13 | 8.80 | 49.10 | 0.68 | 1.48 | 0.85 | 2.28 |
| **11 weeks** | 7.00 | 39.15 | 13.00 | 48.14 | 0.65 | 1.53 | 0.81 | 2.15 |
| **12 weeks** | 7.00 | 37.10 | 15.25 | 46.19 | 0.72 | 1.60 | 0.89 | 2.00 |
| **Mean over all** | **4.69^b^** | **25.27** | **6.63^a^** | **30.14** | **0.65^c^** | **1.23** | **0.81^c^** | **1.91** |
| **L.S.D at 0.05 =** | **46.07** | | | | | | | |
| ***Bemisia tabaci*** | | | | | | | | |
|  | ***B. napus*** | **Check** | ***B. oleracea*** | **Check** | ***M. spicata*** | **Check** | ***M. longifolia*** | **Check** |
| **Precount** | **35.71** | **66.18** | **13.99** | **28.55** | **2.15** | **3.21** | **0.22** | **2.13** |
| **1 week** | 29.52 | 87.96 | 13.25 | 35.19 | 1.66 | 3.90 | 0.10 | 2.68 |
| **2 weeks** | 31.00 | 100.57 | 16.81 | 51.44 | 1.58 | 4.09 | 0.09 | 2.94 |
| **3 weeks** | 48.16 | 120.18 | 43.12 | 62.13 | 1.60 | 5.51 | 0.12 | 3.06 |
| **4 weeks** | 29.11 | 131.20 | 29.64 | 86.15 | 1.79 | 5.10 | 0.23 | 3.60 |
| **5 weeks** | 27.60 | 156.13 | 27.54 | 97.11 | 2.00 | 5.33 | 0.11 | 3.71 |
| **6 weeks** | 35.99 | 161.90 | 34.30 | 113.15 | 1.85 | 5.64 | 0.10 | 3.90 |
| **7 weeks** | 45.00 | 165.21 | 51.74 | 123.90 | 1.89 | 5.93 | 0.19 | 4.00 |
| **8 weeks** | 51.23 | 172.73 | 72.31 | 131.70 | 1.32 | 6.11 | 0.31 | 4.15 |
| **9 weeks** | 69.40 | 181.76 | 83.65 | 145.91 | 1.76 | 6.22 | 0.69 | 4.44 |
| **10 weeks** | 75.13 | 193.12 | 90.15 | 154.20 | 1.88 | 6.34 | 0.72 | 4.60 |
| **11 weeks** | 65.00 | 200.13 | 78.50 | 164.13 | 2.10 | 6.51 | 0.80 | 4.80 |
| **12 weeks** | 82.25 | 225.11 | 89.35 | 173.10 | 2.00 | 6.71 | 1.15 | 5.01 |
| **Mean over all** | **48.08^a^** | **150.94** | **49.57^a^** | **105.13** | **1.81^b^** | **5.43** | **0.37^c^** | **3.77** |
| **L.S.D at 0.05 =** | **30.73** | | | | | | | |

| **Table (15). Reduction percentage of *T. tabaci* and *B. tabaci* on Russian kale *B. napus var. pabularia.*** | | | | | | | | | | | |
| --- | --- | --- | --- | --- | --- | --- | --- | --- | --- | --- | --- |
| ***Thrips tabaci*** | | | | | | | | | | | |
| **Sampling date** | | ***A. swirskii*** | | | ***C. negevi*** | | | **Bio-Magic** | | | **Egyxide** |
| **23 Apr, 2017** | -66.51 | | | -371.36 | | -5.89 | | | | 29.34 | |
| **30 Apr, 2017** | 20.43 | | | -428.01 | | 26.96 | | | | 49.94 | |
| **07 May, 2017** | 68.39 | | | -164.14 | | 48.95 | | | | 75.27 | |
| **14 May, 2017** | 83.54 | | | -85.05 | | 59.84 | | | | 71.45 | |
| **28 May, 2017** | 88.06 | | | -23.68 | | 72.72 | | | | 85.37 | |
| **04 Jun, 2017** | 100.00 | | | -33.89 | | 78.63 | | | | 88.03 | |
| **11 Jun, 2017** | - | | | -40.04 | | 84.12 | | | | 87.18 | |
| **18 Jun, 2017** | - | | | -33.16 | | 85.42 | | | | 80.38 | |
| **25 Jun, 2017** | - | | | 0.48 | | 78.77 | | | | 78.73 | |
| **09 Jul, 2017** | - | | | 39.39 | | 70.04 | | | | 74.09 | |
| **16 July, 2017** | - | | | 44.83 | | 51.87 | | | | 64.59 | |
| **23 Jul, 2017** | - | | | 48.35 | | 61.87 | | | | 68.19 | |
| **30 July, 2017** | - | | | 53.10 | | 59.26 | | | | 54.75 | |
| **06 Aug, 2017** | - | | | 50.00 | | 53.47 | | | | 47.70 | |
| **13 Aug, 2017** | - | | | 47.66 | | 48.69 | | | | 42.23 | |
| **20 Aug, 2017** | - | | | 40.35 | | 39.04 | | | | 29.66 | |
| **Mean** | **48.99 c** | | | **53.45 b** | | **57.11 b** | | | | **64.18 a** | |
| **L.S.D at 0.05 = 11.56** | | | | | | | | | | | |
| ***Bemisia tabaci*** | | | | | | | | | | | |
|  | ***A. swirskii*** | | ***C. negevi*** | | | | **Bio-Magic** | | **Egyxide** | | |
| **14 May, 2017** | 59.03 | | 49.31 | | | | 55.57 | | 62.42 | | |
| **21 May, 2017** | 72.52 | | 63.42 | | | | 67.85 | | 74.64 | | |
| **28 May, 2017** | 79.36 | | 77.80 | | | | 75.29 | | 78.80 | | |
| **04 Jun, 2017** | 76.19 | | 83.29 | | | | 79.21 | | 73.05 | | |
| **18 Jun, 2017** | 55.70 | | 2.20 | | | | 40.45 | | 17.08 | | |
| **25 Jun, 2017** | 74.73 | | 15.84 | | | | 37.68 | | 9.51 | | |
| **02 Jul, 2017** | 85.49 | | 6.49 | | | | 2.29 | | -3.64 | | |
| **09 Jul, 2017** | 89.88 | | 59.41 | | | | 0.51 | | 0.85 | | |
| **16 July, 2017** | 98.06 | | 75.11 | | | | 12.15 | | -29.53 | | |
| **23 Jul, 2017** | 99.71 | | 84.20 | | | | 13.45 | | -37.41 | | |
| **06 Aug, 2017** | 100.00 | | 30.31 | | | | 1.73 | | -20.25 | | |
| **13 Aug, 2017** | - | | 88.07 | | | | -0.87 | | -24.05 | | |
| **20 Aug, 2017** | - | | 100.00 | | | | -2.66 | | -26.71 | | |
| **Mean** | **80.97 a** | | **56.57 b** | | | | **29.45 c** | | **13.45 d** | | |
| **L.S.D at 0.05 = 45.91** | | | | | | | | | | | |

**Fig. (1 a &b). Releasing results of of *P. persmilis* against *T. urticae* associated with Brassicaceae and Lamicaceae medicinal plants in Om Saber.**

a) T. urticae immatures after releasing P. persimilis

|  | Brassica napus var. pabularia | Brassica oleracea var palmifolia | Mentha spicata | Mentha longifolia |
| --- | --- | --- | --- | --- |
| 02 Apr, 2017 | 20.31 | 32.98 | 2 | 2.46 |
| 09 Apr, 2017 | 18.86 | 26.1 | 1.65 | 1.68 |
| 16 Apr, 2017 | 15.05 | 23.3 | 1.22 | 1.5 |
| 23 Apr, 2017 | 12.7 | 20.71 | 0.98 | 0.76 |
| 30 Apr, 2017 | 11.12 | 18.34 | 0.14 | 0.32 |
| 07 May, 2017 | 10.5 | 15.2 | 0.09 | 0.09 |
| 14 May, 2017 | 6.25 | 10.34 | 0 | 0 |
| 21 May, 2017 | 4.4 | 6.12 | 0 | 0 |
| 28 May, 2017 | 2.64 | 2.51 | 0 | 0 |
| 04 Jun, 2017 | 1 | 1 | 0 | 0 |
| 11 Jun, 2017 | 0.01 | 0 | 0 | 0 |
| 18 Jun, 2017 | 0 | 0 | 0 | 0 |
| 25 Jun, 2017 | 0 | 0 | 0 | 0 |
| 02 Jul, 2017 | 0 | 0 | 0 | 0 |
| 09 Jul, 2017 | 0 | 0 | 0 | 0 |
| 16 July, 2017 | 0 | 0 | 0 | 0 |
| 23 Jul, 2017 | 0 | 0.5 | 0 | 0 |
| 30 July, 2017 | 0 | 0 | 0 | 0 |
| 06 Aug, 2017 | 0 | 0 | 0 | 0 |
| 13 Aug, 2017 | 0 | 0 | 0 | 0 |
| 20 Aug, 2017 | 0 | 0 | 0 | 0 |

b) T. urtica adults after releasing P. persimilis

|  | Brassica napus var. pabularia | Brassica oleracea var palmifolia | Mentha spicata | Mentha longifolia |
| --- | --- | --- | --- | --- |
| 02 Apr, 2017 | 27.1 | 26.5 | 1.93 | 2.34 |
| 09 Apr, 2017 | 24.42 | 24.18 | 1.5 | 2 |
| 16 Apr, 2017 | 21.61 | 21.75 | 1.32 | 1.55 |
| 23 Apr, 2017 | 19.05 | 19.12 | 1.15 | 1.1 |
| 30 Apr, 2017 | 16.44 | 18.55 | 1 | 0.89 |
| 07 May, 2017 | 14.1 | 17.9 | 0.84 | 0.44 |
| 14 May, 2017 | 11.22 | 13.6 | 0.44 | 0.21 |
| 21 May, 2017 | 8.37 | 10.53 | 0.08 | 0.08 |
| 28 May, 2017 | 4.34 | 7.46 | 0 | 0 |
| 04 Jun, 2017 | 3.01 | 4.25 | 0 | 0 |
| 11 Jun, 2017 | 1 | 2 | 0 | 0 |
| 18 Jun, 2017 | 0.1 | 0.87 | 0 | 0 |
| 25 Jun, 2017 | 0 | 0.05 | 0 | 0 |
| 02 Jul, 2017 | 0 | 0 | 0 | 0 |
| 09 Jul, 2017 | 0 | 0 | 0 | 0 |
| 16 July, 2017 | 0 | 0 | 0 | 0 |
| 23 Jul, 2017 | 0 | 0 | 0 | 0 |
| 30 July, 2017 | 0 | 0 | 0 | 0 |
| 06 Aug, 2017 | 0 | 0 | 0 | 0 |
| 13 Aug, 2017 | 0 | 0 | 0 | 0 |
| 20 Aug, 2017 | 0 | 0 | 0 | 0 |

a) A. swirskii results on immatures of the TSSM

|  | Brassica napus var. pabularia | Brassica oleracea var palmifolia | Mentha spicata | Mentha longifolia |
| --- | --- | --- | --- | --- |
| 02 Apr, 2017 | 26.48 | 30.11 | 2.01 | 2.25 |
| 09 Apr, 2017 | 24.2 | 28.17 | 1.98 | 2.15 |
| 16 Apr, 2017 | 22.1 | 25.39 | 1.75 | 1.96 |
| 23 Apr, 2017 | 21.54 | 21.98 | 1.44 | 1.58 |
| 30 Apr, 2017 | 20.64 | 19.3 | 1.2 | 1.21 |
| 07 May, 2017 | 20.5 | 17.26 | 0.8 | 1 |
| 14 May, 2017 | 18.21 | 11.05 | 0.45 | 0.84 |
| 21 May, 2017 | 16.1 | 8.2 | 0.05 | 0.6 |
| 28 May, 2017 | 14.34 | 5.18 | 0 | 0.33 |
| 04 Jun, 2017 | 12.26 | 2.41 | 0 | 0.11 |
| 11 Jun, 2017 | 10.14 | 0.92 | 0 | 0.06 |
| 18 Jun, 2017 | 9.44 | 0 | 0 | 0 |
| 25 Jun, 2017 | 9 | 0 | 0 | 0 |
| 02 Jul, 2017 | 8.91 | 0 | 0 | 0 |
| 09 Jul, 2017 | 5.2 | 0 | 0 | 0 |
| 16 July, 2017 | 3.75 | 0 | 0 | 0 |
| 23 Jul, 2017 | 1.1 | 0 | 0 | 0 |
| 30 July, 2017 | 0.72 | 0 | 0 | 0 |
| 06 Aug, 2017 | 0 | 0 | 0 | 0 |
| 13 Aug, 2017 | 0 | 0 | 0 | 0 |
| 20 Aug, 2017 | 0 | 0 | 0 | 0 |

b A. swirskii results on adults of the TSSM

|  | Brassica napus var. pabularia | Brassica oleracea var palmifolia | Mentha spicata | Mentha longifolia |
| --- | --- | --- | --- | --- |
| 02 Apr, 2017 | 26.48 | 30.11 | 2.01 | 2.25 |
| 09 Apr, 2017 | 24.2 | 28.17 | 1.98 | 2.15 |
| 16 Apr, 2017 | 22.1 | 25.39 | 1.75 | 1.96 |
| 23 Apr, 2017 | 21.54 | 21.98 | 1.44 | 1.58 |
| 30 Apr, 2017 | 20.64 | 19.3 | 1.2 | 1.21 |
| 07 May, 2017 | 20.5 | 17.26 | 0.8 | 1 |
| 14 May, 2017 | 18.21 | 11.05 | 0.45 | 0.84 |
| 21 May, 2017 | 16.1 | 8.2 | 0.05 | 0.6 |
| 28 May, 2017 | 14.34 | 5.18 | 0 | 0.33 |
| 04 Jun, 2017 | 12.26 | 2.41 | 0 | 0.11 |
| 11 Jun, 2017 | 10.14 | 0.92 | 0 | 0.06 |
| 18 Jun, 2017 | 9.44 | 0 | 0 | 0 |
| 25 Jun, 2017 | 9 | 0 | 0 | 0 |
| 02 Jul, 2017 | 8.91 | 0 | 0 | 0 |
| 09 Jul, 2017 | 5.2 | 0 | 0 | 0 |
| 16 July, 2017 | 3.75 | 0 | 0 | 0 |
| 23 Jul, 2017 | 1.1 | 0 | 0 | 0 |
| 30 July, 2017 | 0.72 | 0 | 0 | 0 |
| 06 Aug, 2017 | 0 | 0 | 0 | 0 |
| 13 Aug, 2017 | 0 | 0 | 0 | 0 |
| 20 Aug, 2017 | 0 | 0 | 0 | 0 |

**Fig. (2 a &b). Releasing results of of *A. swirskii* against the TSSM associated with Brassicaceae and Lamicaceae medicinal plants in Om Saber.**

a) A. swirskii releasing results on T. tabaci

|  | Brassica napus var. pabularia | Brassica oleracea var palmifolia | Mentha spicata | Mentha longifolia |
| --- | --- | --- | --- | --- |
| 02 Apr, 2017 | 3.84 | 3.75 | 0.18 | 0.06 |
| 09 Apr, 2017 | 2.6 | 2.6 | 0.1 | 0.05 |
| 16 Apr, 2017 | 2.09 | 2.29 | 0.1 | 0.02 |
| 23 Apr, 2017 | 2.05 | 1.85 | 0.07 | 0.01 |
| 30 Apr, 2017 | 1.9 | 1.69 | 0.05 | 0 |
| 07 May, 2017 | 1.85 | 1.65 | 0.01 | 0 |
| 14 May, 2017 | 1.12 | 1.22 | 0 | 0 |
| 21 May, 2017 | 1.31 | 1.1 | 0 | 0 |
| 28 May, 2017 | 0.9 | 0.63 | 0 | 0 |
| 04 Jun, 2017 | 0.65 | 0.2 | 0 | 0 |
| 11 Jun, 2017 | 0.25 | 0 | 0 | 0 |
| 18 Jun, 2017 | 0 | 0 | 0 | 0 |
| 25 Jun, 2017 | 0 | 0 | 0 | 0 |
| 02 Jul, 2017 | 0 | 0 | 0 | 0 |

b) A. swirskii releasing results on B. tabaci

|  | Brassica napus var. pabularia | Brassica oleracea var palmifolia | Mentha spicata | Mentha longifolia |
| --- | --- | --- | --- | --- |
| 02 Apr, 2017 | 8.12 | 3.5 | 3 | 2.15 |
| 09 Apr, 2017 | 6.45 | 2.6 | 2.45 | 1.82 |
| 16 Apr, 2017 | 5.3 | 2.29 | 2.3 | 1.66 |
| 23 Apr, 2017 | 5 | 3.05 | 2.17 | 1.21 |
| 30 Apr, 2017 | 7.05 | 6.9 | 2.06 | 1.05 |
| 07 May, 2017 | 9.24 | 10.85 | 1 | 0.56 |
| 14 May, 2017 | 4.68 | 7.12 | 0.19 | 0.2 |
| 21 May, 2017 | 3.19 | 5.31 | 0.06 | 0.09 |
| 28 May, 2017 | 2 | 3.29 | 0 | 0 |
| 04 Jun, 2017 | 1.66 | 1.65 | 0 | 0 |
| 11 Jun, 2017 | 0.33 | 1.25 | 0 | 0 |
| 18 Jun, 2017 | 0.05 | 1 | 0 | 0 |
| 25 Jun, 2017 | 0 | 1.05 | 0 | 0 |
| 02 Jul, 2017 | 0 | 0.05 | 0 | 0 |
| 09 Jul, 2017 | 0 | 0 | 0 | 0 |

**Fig. (3 a &b). Releasing results of of *A. swirskii* against *T. tabaci* and *B. tabaci* associated with Brassicaceae and Lamicaceae medicinal plants in Om Saber, season 2017.**

a) C. negevi releasing results against T. urticae immatures

|  | Brassica napus var. pabularia | Brassica oleracea var palmifolia | Mentha spicata | Mentha longifolia |
| --- | --- | --- | --- | --- |
| 02 Apr, 2017 | 26.48 | 30.15 | 1.98 | 2.35 |
| 09 Apr, 2017 | 25.13 | 25.3 | 1.83 | 2.2 |
| 16 Apr, 2017 | 25 | 21.15 | 1.65 | 2.1 |
| 23 Apr, 2017 | 26.4 | 20.48 | 1.33 | 1.95 |
| 30 Apr, 2017 | 27.33 | 19.23 | 1.18 | 1.83 |
| 07 May, 2017 | 27.9 | 19 | 1 | 1.62 |
| 14 May, 2017 | 25.21 | 17.59 | 0.87 | 1.4 |
| 21 May, 2017 | 23.12 | 16.35 | 0.54 | 1.25 |
| 28 May, 2017 | 21.45 | 14.3 | 0.29 | 1.12 |
| 04 Jun, 2017 | 20.6 | 14.84 | 0.08 | 1 |
| 11 Jun, 2017 | 20.34 | 13.7 | 0 | 0.87 |
| 18 Jun, 2017 | 19 | 13 | 0 | 0.66 |
| 25 Jun, 2017 | 18.97 | 12.79 | 0 | 0.54 |
| 02 Jul, 2017 | 19.01 | 12.15 | 0 | 0.2 |
| 09 Jul, 2017 | 16.49 | 11.75 | 0 | 0.09 |
| 16 July, 2017 | 14.24 | 11 | 0 | 0 |
| 23 Jul, 2017 | 11.3 | 10.5 | 0 | 0 |
| 30 July, 2017 | 10.67 | 8.66 | 0 | 0 |
| 06 Aug, 2017 | 9.8 | 7.75 | 0 | 0 |
| 13 Aug, 2017 | 9.49 | 7 | 0 | 0 |
| 20 Aug, 2017 | 8.9 | 7 | 0 | 0 |

b) C. negevi releasing results against T. urticae adults

|  | Brassica napus var. pabularia | Brassica oleracea var palmifolia | Mentha spicata | Mentha longifolia |
| --- | --- | --- | --- | --- |
| 02 Apr, 2017 | 26.22 | 26.75 | 1.87 | 2.63 |
| 09 Apr, 2017 | 25.3 | 24.3 | 1.85 | 2.41 |
| 16 Apr, 2017 | 23.84 | 25 | 1.81 | 2.38 |
| 23 Apr, 2017 | 22.86 | 27.1 | 1.79 | 2.35 |
| 30 Apr, 2017 | 21.71 | 28.37 | 1.75 | 2.33 |
| 07 May, 2017 | 20.17 | 29.5 | 1.7 | 2.3 |
| 14 May, 2017 | 18.35 | 23.1 | 1.66 | 2.24 |
| 21 May, 2017 | 17.57 | 21.65 | 1.64 | 2.21 |
| 28 May, 2017 | 16.25 | 20.58 | 1.61 | 2.18 |
| 04 Jun, 2017 | 16 | 19.79 | 1.58 | 2.15 |
| 11 Jun, 2017 | 17.64 | 18 | 1.55 | 2.12 |
| 18 Jun, 2017 | 19.11 | 17.67 | 1.49 | 2.09 |
| 25 Jun, 2017 | 21.8 | 18.1 | 1.47 | 2.05 |
| 02 Jul, 2017 | 23.7 | 14.37 | 1.46 | 2 |
| 09 Jul, 2017 | 21.34 | 13.5 | 1.31 | 1.9 |
| 16 July, 2017 | 20.5 | 12.1 | 1.29 | 1.85 |
| 23 Jul, 2017 | 19.15 | 11.7 | 1.27 | 1.81 |
| 30 July, 2017 | 18.7 | 13.01 | 1.25 | 1.74 |
| 06 Aug, 2017 | 21.43 | 14.82 | 1.2 | 1.65 |
| 13 Aug, 2017 | 26.5 | 16 | 1.18 | 1.49 |
| 20 Aug, 2017 | 30.25 | 18.33 | 1.11 | 1.38 |

**Fig. (4 a & b). Releasing results of of *C. negevi* against *T. urticae* associated with Brassicaceae and Lamicaceae medicinal plants in Om Saber.**

a) C. negevi releasing results on T. tabaci

|  | Brassica napus var. pabularia | Brassica oleracea var palmifolia | Mentha spicata | Mentha longifolia |
| --- | --- | --- | --- | --- |
| 16 Apr, 2017 | 0.81 | 2.4 |  |  |
| 23 Apr, 2017 | 0.92 | 3.8 |  |  |
| 30 Apr, 2017 | 1.3 | 5 |  |  |
| 07 May, 2017 | 3.2 | 4.96 |  |  |
| 14 May, 2017 | 5.11 | 5 |  |  |
| 21 May, 2017 | 6.2 | 10.37 | 0.17 | 0.04 |
| 28 May, 2017 | 7.15 | 12.11 | 0.58 | 0.73 |
| 04 Jun, 2017 | 8.41 | 14.72 | 0.68 | 0.81 |
| 11 Jun, 2017 | 12.3 | 16.13 | 0.9 | 1.1 |
| 18 Jun, 2017 | 15.3 | 21.11 | 0.97 | 1.32 |
| 25 Jun, 2017 | 23.2 | 22 | 0.98 | 1.65 |
| 02 Jul, 2017 | 27.25 | 28.76 | 1 | 1.99 |
| 09 Jul, 2017 | 30.9 | 33.6 | 1.09 | 2.1 |
| 16 July, 2017 | 35.14 | 35.25 | 1.15 | 2.09 |
| 23 Jul, 2017 | 42.1 | 39.98 | 1.36 | 2.23 |
| 30 July, 2017 | 44 | 42.9 | 1.49 | 2.25 |
| 06 Aug, 2017 | 38.6 | 45.12 | 1.5 | 2.12 |
| 13 Aug, 2017 | 36.15 | 47.09 | 1.49 | 2 |
| 20 Aug, 2017 | 33.6 | 45 | 1.38 | 1.9 |

b) C. negevi releasing results on B. tabaci

|  | Brassica napus var. pabularia | Brassica oleracea var palmifolia | Mentha spicata | Mentha longifolia |
| --- | --- | --- | --- | --- |
| 02 Apr, 2017 | 8.17 | 2.4 |  |  |
| 09 Apr, 2017 | 7 | 2 |  |  |
| 16 Apr, 2017 | 6.87 | 1.79 |  |  |
| 23 Apr, 2017 | 6.98 | 1.96 |  |  |
| 30 Apr, 2017 | 7.35 | 3 |  |  |
| 07 May, 2017 | 9.42 | 8.45 |  |  |
| 14 May, 2017 | 6.56 | 6.11 |  |  |
| 21 May, 2017 | 5 | 6.12 | 2.37 | 2.18 |
| 28 May, 2017 | 4.6 | 6.13 | 1.8 | 2.1 |
| 04 Jun, 2017 | 3.47 | 8.11 | 1.68 | 2 |
| 11 Jun, 2017 | 3.88 | 9 | 1.5 | 1.84 |
| 18 Jun, 2017 | 3.99 | 10.06 | 1.49 | 1.66 |
| 25 Jun, 2017 | 4.84 | 11.1 | 1.12 | 1.42 |
| 02 Jul, 2017 | 2.5 | 7.12 | 1 | 1.33 |
| 09 Jul, 2017 | 1.59 | 6.61 | 0.9 | 1.12 |
| 16 July, 2017 | 1.03 | 4.9 | 0.24 | 1.02 |
| 23 Jul, 2017 | 0.75 | 3.22 | 0.1 | 0.7 |
| 30 July, 2017 | 0.55 | 2.02 | 0 | 0.43 |
| 06 Aug, 2017 | 0.1 | 0.5 | 0 | 0.13 |
| 13 Aug, 2017 | 0 | 0.01 | 0 | 0.04 |
| 20 Aug, 2017 | 0 | 0 | 0 | 0 |

**Fig. (5). Releasing results of of *C. negevi* against *T. tabaci* and *B. tabaci* associated with Brassicaceae and Lamicaceae medicinal plants in Om Saber**

a) Bio-Magic application results on the TSSM immatures

|  | Brassica napus var. pabularia | Brassica oleracea var palmifolia | Mentha spicata | Mentha longifolia |
| --- | --- | --- | --- | --- |
| 02 Apr, 2017 | 19.68 | 17 | 1.89 | 2.17 |
| 09 Apr, 2017 | 14.25 | 11.52 | 0.9 | 1.2 |
| 16 Apr, 2017 | 12.5 | 11.29 | 0.93 | 1.18 |
| 23 Apr, 2017 | 13 | 12.64 | 1.05 | 2 |
| 30 Apr, 2017 | 18.85 | 13.72 | 1.13 | 2.59 |
| 07 May, 2017 | 25.6 | 16.84 | 1.27 | 3.1 |
| 14 May, 2017 | 13.5 | 10.42 | 0.51 | 1 |
| 21 May, 2017 | 21.3 | 10.38 | 0.48 | 0.9 |
| 28 May, 2017 | 32.9 | 11.48 | 0.45 | 0.97 |
| 04 Jun, 2017 | 38.75 | 24.5 | 0.67 | 1.1 |
| 11 Jun, 2017 | 40 | 29.66 | 0.89 | 2.43 |
| 18 Jun, 2017 | 47.25 | 33.84 | 1.15 | 3.87 |
| 25 Jun, 2017 | 17.9 | 13.41 | 1.48 | 4.6 |
| 02 Jul, 2017 | 22.45 | 13.36 | 1.77 | 5.32 |
| 09 Jul, 2017 | 30.25 | 16.67 | 1.74 | 2.97 |
| 16 July, 2017 | 34.15 | 19.51 | 1.55 | 2.75 |
| 23 Jul, 2017 | 36.75 | 25.1 | 2.03 | 3 |
| 30 July, 2017 | 39.8 | 33.47 | 2.15 | 3.11 |

b) Bio-Magic application results on the TSSM adults

|  | Brassica napus var. pabularia | Brassica oleracea var palmifolia | Mentha spicata | Mentha longifolia |
| --- | --- | --- | --- | --- |
| 02 Apr, 2017 | 17 | 11.42 | 3.45 | 2.17 |
| 09 Apr, 2017 | 12.2 | 7 | 3.51 | 1.2 |
| 16 Apr, 2017 | 11 | 8.81 | 3.42 | 1.18 |
| 23 Apr, 2017 | 13.5 | 13.34 | 3.34 | 2 |
| 30 Apr, 2017 | 16.35 | 20.5 | 3.1 | 2.59 |
| 07 May, 2017 | 19.45 | 28 | 3.25 | 3.1 |
| 14 May, 2017 | 10.5 | 26.7 | 3.39 | 1 |
| 21 May, 2017 | 12 | 26 | 3.32 | 0.9 |
| 28 May, 2017 | 17.65 | 28.88 | 3.15 | 0.97 |
| 04 Jun, 2017 | 25.8 | 29.91 | 3 | 1.1 |
| 11 Jun, 2017 | 29.25 | 31.62 | 3.26 | 2.43 |
| 18 Jun, 2017 | 37.4 | 33.88 | 3.59 | 3.87 |
| 25 Jun, 2017 | 18.29 | 14.5 | 3.82 | 4.6 |
| 02 Jul, 2017 | 17.55 | 14.48 | 4 | 5.32 |
| 09 Jul, 2017 | 20.43 | 21.9 | 4.35 | 2.97 |
| 16 July, 2017 | 22.18 | 27.1 | 4.21 | 2.75 |
| 23 Jul, 2017 | 25.6 | 28 | 3.89 | 3 |
| 30 July, 2017 | 29.97 | 28.9 | 3.76 | 3.11 |

**Fig. (6 a & b). Application results of of Bio-Magic against the *T. urticae* associated with Brassicaceae and Lamicaceae medicinal plants in Om Saber.**

a) Bio-Magic application results on T. tabaci

|  | Brassica napus var. pabularia | Brassica oleracea var palmifolia | Mentha spicata | Mentha longifolia |
| --- | --- | --- | --- | --- |
| 21 May, 2017 | 1.08 | 1.51 | 0.62 | 0.8 |
| 28 May, 2017 | 1.9 | 1.8 | 0.39 | 0.49 |
| 04 Jun, 2017 | 1 | 1.16 | 0.35 | 0.47 |
| 11 Jun, 2017 | 0.95 | 1.1 | 0.42 | 0.5 |
| 18 Jun, 2017 | 3.23 | 2.95 | 0.84 | 1 |
| 25 Jun, 2017 | 6.4 | 7.15 | 1.1 | 1.19 |
| 02 Jul, 2017 | 3 | 4.66 | 0.5 | 0.76 |
| 09 Jul, 2017 | 2.64 | 4.5 | 0.48 | 0.73 |
| 16 July, 2017 | 4.15 | 7.9 | 0.65 | 0.89 |
| 23 Jul, 2017 | 9.72 | 13.81 | 0.84 | 1 |
| 30 July, 2017 | 5.86 | 9.21 | 1.1 | 0.85 |
| 06 Aug, 2017 | 6 | 10.75 | 1.25 | 0.92 |
| 13 Aug, 2017 | 6.1 | 11 | 1.32 | 1.1 |

b) Bio-Magic application results on B. tabaci

|  | Brassica napus var. pabularia | Brassica oleracea var palmifolia | Mentha spicata | Mentha longifolia |
| --- | --- | --- | --- | --- |
| 21 May, 2017 | 52.38 | 26.8 | 1.89 | 0.24 |
| 28 May, 2017 | 45.12 | 16.2 | 1.56 | 0.16 |
| 04 Jun, 2017 | 40 | 15.85 | 1.23 | 0.13 |
| 11 Jun, 2017 | 38.61 | 20.52 | 1.13 | 0.11 |
| 18 Jun, 2017 | 39.7 | 50.46 | 1.49 | 0.15 |
| 25 Jun, 2017 | 23.5 | 31.67 | 1.8 | 0.09 |
| 02 Jul, 2017 | 16 | 30.14 | 1.9 | 0.07 |
| 09 Jul, 2017 | 20.01 | 32.21 | 2.15 | 0.1 |
| 16 July, 2017 | 34.25 | 45.16 | 1.96 | 0.39 |
| 23 Jul, 2017 | 41.5 | 60 | 1.95 | 0.75 |
| 30 July, 2017 | 38 | 57.55 | 1.67 | 0.7 |
| 06 Aug, 2017 | 38.2 | 58.25 | 1.89 | 0.95 |
| 13 Aug, 2017 | 44 | 63.1 | 2 | 1.15 |

**Fig. (7 a & b). Application results of Bio-Magic against *T. tabaci* and *B. tabaci* associated with Brassicaceae and Lamicaceae medicinal plants in Om Saber.**

a) Egyxide application results on the TSSM immatures

|  | Brassica napus var. pabularia | Brassica oleracea var palmifolia | Mentha spicata | Mentha longifolia |
| --- | --- | --- | --- | --- |
| 02 Apr, 2017 | 19.5 | 16.86 | 2 | 2.71 |
| 09 Apr, 2017 | 9.36 | 10.34 | 0.5 | 1 |
| 16 Apr, 2017 | 10.45 | 10.2 | 0.48 | 1.01 |
| 23 Apr, 2017 | 16.75 | 11 | 0.69 | 1.21 |
| 30 Apr, 2017 | 23.6 | 12.36 | 0.94 | 2.3 |
| 07 May, 2017 | 27.22 | 14.94 | 1.08 | 2.89 |
| 14 May, 2017 | 10.45 | 9 | 0.2 | 0.5 |
| 21 May, 2017 | 17.08 | 9.35 | 0.17 | 0.47 |
| 28 May, 2017 | 22.9 | 10.88 | 0.35 | 0.73 |
| 04 Jun, 2017 | 26.55 | 32.49 | 0.88 | 1.38 |
| 11 Jun, 2017 | 32.45 | 34.95 | 1.2 | 2.27 |
| 18 Jun, 2017 | 40.15 | 40 | 1.95 | 3.12 |
| 25 Jun, 2017 | 15 | 19.05 | 2.07 | 3.9 |
| 02 Jul, 2017 | 20.15 | 18.22 | 2.14 | 4.87 |
| 09 Jul, 2017 | 33.75 | 22.5 | 0.35 | 2 |
| 16 July, 2017 | 36.55 | 26.1 | 0.96 | 2.1 |
| 23 Jul, 2017 | 38.04 | 33.78 | 1.25 | 3.05 |
| 30 July, 2017 | 40.8 | 43.96 | 1.78 | 3.2 |

b) Egyxide application results on the TSSM adults

|  | Brassica napus var. pabularia | Brassica oleracea var palmifolia | Mentha spicata | Mentha longifolia |
| --- | --- | --- | --- | --- |
| 02 Apr, 2017 | 16.8 | 11.35 | 3.57 | 2.71 |
| 09 Apr, 2017 | 16.8 | 10.12 | 2.45 | 1 |
| 16 Apr, 2017 | 9.64 | 12 | 2.4 | 1.01 |
| 23 Apr, 2017 | 17.5 | 16.49 | 2.64 | 1.21 |
| 30 Apr, 2017 | 20.7 | 26.72 | 2.89 | 2.3 |
| 07 May, 2017 | 23.25 | 32.64 | 3 | 2.89 |
| 14 May, 2017 | 15 | 29.51 | 2.69 | 0.5 |
| 21 May, 2017 | 14.62 | 28.75 | 2.58 | 0.47 |
| 28 May, 2017 | 25.12 | 30.18 | 2.79 | 0.73 |
| 04 Jun, 2017 | 30.46 | 32.97 | 2.98 | 1.38 |
| 11 Jun, 2017 | 38.1 | 36 | 3.12 | 2.27 |
| 18 Jun, 2017 | 45.12 | 40.44 | 3.33 | 3.12 |
| 25 Jun, 2017 | 19.02 | 25.1 | 3.86 | 3.9 |
| 02 Jul, 2017 | 20.56 | 26.6 | 4.15 | 4.87 |
| 09 Jul, 2017 | 27.12 | 31.56 | 3.4 | 2 |
| 16 July, 2017 | 30 | 35.15 | 3.29 | 2.1 |
| 23 Jul, 2017 | 34.85 | 38 | 3.52 | 3.05 |
| 30 July, 2017 | 39.5 | 39.09 | 3.7 | 3.2 |

**Fig. (8 a & b). Application results of Egyxide against *T. urticae* associated with Brassicaceae and Lamicaceae medicinal plants in Om Saber.**

a) Egyxide application results on T. tabaci

|  | Brassica napus var. pabularia | Brassica oleracea var palmifolia | Mentha spicata | Mentha longifolia |
| --- | --- | --- | --- | --- |
| 30 Apr, 2017 | 1 | 1.49 |  |  |
| 07 May, 2017 | 2.05 | 1.79 |  |  |
| 14 May, 2017 | 0.9 | 0.99 |  |  |
| 21 May, 2017 | 0.93 | 1.05 |  |  |
| 28 May, 2017 | 2.66 | 2.25 | 0.72 | 0.81 |
| 04 Jun, 2017 | 6 | 6.18 | 0.6 | 0.78 |
| 11 Jun, 2017 | 3.15 | 3.54 | 0.44 | 0.52 |
| 18 Jun, 2017 | 3 | 3.29 | 0.42 | 0.5 |
| 25 Jun, 2017 | 5.9 | 8.44 | 0.49 | 0.83 |
| 02 Jul, 2017 | 10.18 | 12.6 | 0.79 | 1.1 |
| 09 Jul, 2017 | 6.12 | 8.99 | 1 | 1.05 |
| 16 July, 2017 | 6.1 | 8.8 | 0.68 | 0.85 |
| 23 Jul, 2017 | 7 | 13 | 0.65 | 0.81 |
| 30 July, 2017 | 7 | 15.25 | 0.72 | 0.89 |

b) Egyxide application results on B. tabaci

|  | Brassica napus var. pabularia | Brassica oleracea var palmifolia | Mentha spicata | Mentha longifolia |
| --- | --- | --- | --- | --- |
| 30 Apr, 2017 | 49.01 | 27.44 |  |  |
| 07 May, 2017 | 35.71 | 13.99 |  |  |
| 14 May, 2017 | 29.52 | 13.25 | 2.15 | 0.22 |
| 21 May, 2017 | 31 | 16.81 | 1.66 | 0.1 |
| 28 May, 2017 | 48.16 | 43.12 | 1.58 | 0.09 |
| 04 Jun, 2017 | 29.11 | 29.64 | 1.6 | 0.12 |
| 11 Jun, 2017 | 27.6 | 27.54 | 1.79 | 0.23 |
| 18 Jun, 2017 | 35.99 | 34.3 | 2 | 0.11 |
| 25 Jun, 2017 | 45 | 51.74 | 1.85 | 0.1 |
| 02 Jul, 2017 | 51.23 | 72.31 | 1.89 | 0.19 |
| 09 Jul, 2017 | 69.4 | 83.65 | 1.32 | 0.31 |
| 16 July, 2017 | 75.13 | 90.15 | 1.76 | 0.69 |
| 23 Jul, 2017 | 65 | 78.5 | 1.88 | 0.72 |
| 30 July, 2017 | 82.25 | 89.35 | 2.1 | 0.8 |

**Fig. (9 a & b). Application results of Egyxide against *T. tabaci* and *B. tabaci* associated with Brassicaceae and Lamicaceae medicinal plants in Om Saber.**

| **Table (16). Recorded weather data in Kom Oshim, Fayoum Governorate during experimental season 2017.** | | | | |
| --- | --- | --- | --- | --- |
| **Sample date** | **Optimum Temperature**  **°C (Max. – Min.)** | | **Relative Humidity**  **R.H.% (Max. – Min.)** | |
| **04 Apr, 2017** | 20.50 | (24-17) | 55.50 | (77-34) |
| **11 Apr, 2017** | 20.50 | (24-17) | 55.50 | (77-34) |
| **18 Apr, 2017** | 26.50 | (32-21) | 35.50 | (57-14) |
| **25 Apr, 2017** | 21.50 | (26-17) | 48.00 | (68-28) |
| **02 May, 2017** | 24.50 | (29-20) | 57.00 | (83-31) |
| **09 May, 2017** | 30.50 | (39-22) | 22.00 | (34-10) |
| **16 May, 2017** | 28.00 | (34-22) | 50.50 | (78-23) |
| **23 May, 2017** | 24.50 | (29-20) | 52.00 | (73-31) |
| **30 May, 2017** | 28.50 | (35-22) | 48.00 | (78-18) |
| **06 Jun, 2017** | 26.00 | (31-21) | 57.50 | (88-27) |
| **13 Jun, 2017** | 30.00 | (37-23) | 49.00 | (78-20) |
| **20 Jun, 2017** | 29.00 | (33-25) | 61.50 | (89-34) |
| **27 Jun, 2017** | 30.50 | (36-25) | 55.50 | (83-28) |
| **04 Jul, 2017** | 31.50 | (37-26) | 62.00 | (89-35) |
| **11 Jul, 2017** | 32.00 | (37-27) | 45.00 | (74-16) |
| **18 Jul, 2017** | 33.00 | (39-27) | 47.50 | (79-16) |
| **25 Jul, 2017** | 31.50 | (37-26) | 60.00 | (89-31) |
| **01 Aug, 2017** | 31.50 | (37-26) | 59.00 | (94-24) |
| **08 Aug, 2017** | 31.00 | (35-27) | 62.50 | (84-41) |
| **15 Aug, 2017** | 30.00 | (34-26) | 60.00 | (79-41) |
| **22 Aug, 2017** | 30.50 | (35-26) | 57.00 | (84-30) |
| **29 Aug, 2017** | 29.50 | (34-25) | 55.00 | (78-32) |

tested organic medicinal plant species

| **Table (17). Mean number *T. urticae* after releasing *P. persimilis* in Kom Oshim.** | | | | | | | | |
| --- | --- | --- | --- | --- | --- | --- | --- | --- |
|  | ***B. napus*** | **check** | ***B. oleracea*** | **check** | ***M. spicata*** | **check** | ***M. longifolia*** | **check** |
| **Precount** | **38.83** | **36.45** | **27.63** | **29.25** | **5.40** | **4.26** | **6.70** | **5.80** |
| **04 April, 2017** | 23.22 | 41.30 | 20.46 | 32.98 | 4.85 | 4.46 | 5.04 | 6.29 |
| **11 April, 2017** | 18.20 | 44.67 | 16.48 | 36.40 | 3.21 | 5.09 | 3.80 | 7.36 |
| **18 April, 2017** | 16.53 | 49.23 | 14.89 | 42.00 | 2.58 | 5.82 | 2.80 | 8.66 |
| **25 April, 2017** | 14.11 | 53.03 | 10.67 | 46.26 | 1.55 | 5.94 | 1.90 | 9.65 |
| **02 May, 2017** | 10.34 | 56.37 | 6.56 | 51.40 | 1.48 | 6.15 | 1.22 | 10.61 |
| **09 May, 2017** | 6.65 | 61.28 | 3.36 | 58.95 | 1.19 | 6.75 | 0.31 | 12.15 |
| **16 May, 2017** | 2.90 | 65.39 | 2.30 | 65.32 | 1.00 | 7.17 | 0.20 | 13.05 |
| **23 May, 2017** | 1.00 | 71.21 | 0.73 | 70.66 | 0.94 | 9.10 | 0.12 | 14.59 |
| **30 May, 2017** | 0.29 | 79.60 | 0.00 | 78.89 | 0.69 | 9.59 | 0.06 | 15.37 |
| **06 June, 2017** | 0.00 | 86.75 | 0.00 | 84.65 | 0.26 | 10.50 | 0.00 | 16.49 |
| **13 June, 2017** | 0.00 | 104.60 | 0.00 | 87.95 | 0.08 | 10.98 | 0.00 | 17.60 |
| **20 June, 2017** | 0.87 | 102.38 | 0.00 | 87.37 | 1.16 | 11.53 | 0.00 | 18.72 |
| **27 June, 2017** | 0.00 | 98.60 | 0.00 | 94.26 | 0.00 | 10.73 | 0.00 | 18.25 |
| **Mean over all** | **9.50** | **67.92** | **7.36** | **61.88** | **1.74** | **7.72** | **1.58** | **12.47** |
| ***P*_0.05_** | **0.000** |  | **0.000** |  | **0.000** |  | **0.000** |  |

| **Table (18). Mean number *T. urticae* after releasing *A. swirskii*, Kom Oshim.** | | | | | | | | |
| --- | --- | --- | --- | --- | --- | --- | --- | --- |
|  | ***B. napus*** | **Check** | ***B. oleracea*** | **Check** | ***M. spicata*** | **Check** | **M. longifolia** | **Check** |
| **Precount** | **36.61** | **36.45** | **26.73** | **29.25** | **5.51** | **4.26** | **4.64** | **5.80** |
| **04 April, 2017** | 30.70 | 41.30 | 26.00 | 32.98 | 5.42 | 4.46 | 4.40 | 6.29 |
| **11 April, 2017** | 26.61 | 44.67 | 25.22 | 36.40 | 5.14 | 5.09 | 4.22 | 7.36 |
| **18 April, 2017** | 25.90 | 49.23 | 26.15 | 42.00 | 4.64 | 5.82 | 4.14 | 8.66 |
| **25 April, 2017** | 24.35 | 53.03 | 33.19 | 46.26 | 4.32 | 5.94 | 4.00 | 9.65 |
| **02 May, 2017** | 20.90 | 56.37 | 42.02 | 51.40 | 3.90 | 6.15 | 3.90 | 10.61 |
| **09 May, 2017** | 18.45 | 61.28 | 45.54 | 58.95 | 3.45 | 6.75 | 3.08 | 12.15 |
| **16 May, 2017** | 15.35 | 65.39 | 41.73 | 65.32 | 3.10 | 7.17 | 2.20 | 13.05 |
| **23 May, 2017** | 13.30 | 71.21 | 40.01 | 70.66 | 2.25 | 9.10 | 1.80 | 14.59 |
| **30 May, 2017** | 13.05 | 79.60 | 36.93 | 78.89 | 1.65 | 9.59 | 1.32 | 15.37 |
| **13 June, 2017** | 12.40 | 86.75 | 36.72 | 84.65 | 1.22 | 10.50 | 0.24 | 16.49 |
| **20 June, 2017** | 17.70 | 104.60 | 39.40 | 87.95 | 1.00 | 10.98 | 0.06 | 17.60 |
| **27 June, 2017** | 12.75 | 102.38 | 26.22 | 87.37 | 0.86 | 11.53 | 0.00 | 18.72 |
| **04 July, 2017** | 10.53 | 98.60 | 23.46 | 94.26 | 0.20 | 10.73 | 0.00 | 18.25 |
| **11 July, 2017** | 9.81 | 94.50 | 19.50 | 90.65 | 0.00 | 10.05 | 0.00 | 17.00 |
| **18 July, 2017** | 10.22 | 89.98 | 14.36 | 82.53 | 0.00 | 9.65 | 0.00 | 16.49 |
| **25 July, 2017** | 11.05 | 87.60 | 12.19 | 76.30 | 0.00 | 8.95 | 0.00 | 14.82 |
| **Mean over all** | **12.11** | **92.22** | **9.23** | **71.21** | **0.00** | **8.55** | **0.00** | **13.91** |
| ***P*_0.05_** | **0.000** |  | **0.000** |  | **0.002** |  | **0.000** |  |

| **Table (19). Mean number of *T. urticae* after releasing *C. negevi*, Kom Oshim.** | | | | | | | | |
| --- | --- | --- | --- | --- | --- | --- | --- | --- |
|  | ***B. napus*** | **Check** | ***B. oleracea*** | **Check** | ***M. spicata*** | **Check** | ***M. longifolia*** | **Check** |
| **Precount** | **36.34** | **36.45** | **26.69** | **29.25** | **5.41** | **4.26** | **4.70** | **5.80** |
| **04 Apr, 2017** | 32.48 | 41.30 | 28.81 | 32.98 | 5.22 | 4.46 | 4.52 | 6.29 |
| **11 Apr, 2017** | 29.60 | 44.67 | 28.71 | 36.40 | 4.91 | 5.09 | 4.36 | 7.36 |
| **18 Apr, 2017** | 27.80 | 49.23 | 32.34 | 42.00 | 4.50 | 5.82 | 4.00 | 8.66 |
| **25 Apr, 2017** | 24.30 | 53.03 | 35.89 | 46.26 | 4.28 | 5.94 | 3.78 | 9.65 |
| **02 May, 2017** | 27.00 | 56.37 | 44.17 | 51.40 | 3.98 | 6.15 | 3.80 | 10.61 |
| **09 May, 2017** | 21.37 | 61.28 | 45.20 | 58.95 | 3.54 | 6.75 | 3.70 | 12.15 |
| **16 May, 2017** | 23.84 | 65.39 | 45.21 | 65.32 | 3.09 | 7.17 | 3.54 | 13.05 |
| **23 May, 2017** | 23.20 | 71.21 | 47.40 | 70.66 | 2.94 | 9.10 | 3.20 | 14.59 |
| **30 May, 2017** | 23.00 | 79.60 | 48.13 | 78.89 | 2.76 | 9.59 | 3.02 | 15.37 |
| **06 Jun, 2017** | 25.03 | 86.75 | 48.95 | 84.65 | 2.29 | 10.50 | 2.91 | 16.49 |
| **13 Jun, 2017** | 28.39 | 104.60 | 47.59 | 87.95 | 2.14 | 10.98 | 2.77 | 17.60 |
| **20 Jun, 2017** | 20.24 | 102.38 | 46.32 | 87.37 | 2.00 | 11.53 | 2.73 | 18.72 |
| **27 Jun, 2017** | 17.28 | 98.60 | 41.43 | 94.26 | 1.50 | 10.73 | 2.71 | 18.25 |
| **04 Jul, 2017** | 17.15 | 94.50 | 34.82 | 90.65 | 1.22 | 10.05 | 2.64 | 17.00 |
| **11 Jul, 2017** | 19.73 | 89.98 | 33.11 | 82.53 | 1.08 | 9.65 | 2.58 | 16.49 |
| **18 Jul, 2017** | 18.64 | 87.60 | 28.79 | 76.30 | 0.60 | 8.95 | 2.45 | 14.82 |
| **25 Jul, 2017** | 20.88 | 92.22 | 24.63 | 71.21 | 0.22 | 8.55 | 2.24 | 13.91 |
| **Mean over all** | **24.24** | **73.06** | **38.23** | **65.95** | **2.87** | **8.07** | **3.31** | **13.16** |
| ***P*_0.05_** | **0.000** |  | **0.000** |  | **0.002** |  | **0.000** |  |

| **Table (20). Mean number *T. urticae* after releasing Bio-Magic, Kom Oshim.** | | | | | | | | |
| --- | --- | --- | --- | --- | --- | --- | --- | --- |
|  | ***B. napus*** | **Check** | ***B. oleracea*** | **Check** | ***M. spicata*** | **Check** | ***M. longifolia*** | **Check** |
| **Precount** | **36.68** | **36.45** | **28.42** | **29.25** | **5.34** | **4.26** | **5.17** | **5.80** |
| **04 Apr, 2017** | 26.45 | 41.30 | 18.52 | 32.98 | 4.41 | 4.46 | 4.04 | 6.29 |
| **11 Apr, 2017** | 23.50 | 44.67 | 20.10 | 36.40 | 4.35 | 5.09 | 3.97 | 7.36 |
| **18 Apr, 2017** | 26.50 | 49.23 | 25.98 | 42.00 | 4.39 | 5.82 | 4.83 | 8.66 |
| **25 Apr, 2017** | 35.20 | 53.03 | 34.22 | 46.26 | 4.88 | 5.94 | 5.58 | 9.65 |
| **02 May, 2017** | 45.05 | 56.37 | 44.84 | 51.40 | 5.22 | 6.15 | 6.20 | 10.61 |
| **09 May, 2017** | 24.00 | 61.28 | 37.12 | 58.95 | 3.90 | 6.75 | 3.50 | 12.15 |
| **16 May, 2017** | 33.30 | 65.39 | 36.38 | 65.32 | 3.80 | 7.17 | 3.35 | 13.05 |
| **23 May, 2017** | 50.55 | 71.21 | 40.36 | 70.66 | 3.60 | 9.10 | 3.66 | 14.59 |
| **30 May, 2017** | 64.55 | 79.60 | 54.41 | 78.89 | 3.67 | 9.59 | 4.05 | 15.37 |
| **06 Jun, 2017** | 69.25 | 86.75 | 61.28 | 84.65 | 4.15 | 10.50 | 5.63 | 16.49 |
| **13 Jun, 2017** | 84.65 | 104.60 | 67.72 | 87.95 | 4.74 | 10.98 | 7.75 | 17.60 |
| **20 Jun, 2017** | 36.19 | 102.38 | 27.91 | 87.37 | 5.30 | 11.53 | 8.92 | 18.72 |
| **27 Jun, 2017** | 40.00 | 98.60 | 27.84 | 94.26 | 5.77 | 10.73 | 10.57 | 18.25 |
| **04 Jul, 2017** | 50.68 | 94.50 | 38.57 | 90.65 | 6.09 | 10.05 | 6.97 | 17.00 |
| **11 Jul, 2017** | 56.33 | 89.98 | 46.61 | 82.53 | 5.76 | 9.65 | 7.50 | 16.49 |
| **18 Jul, 2017** | 62.35 | 87.60 | 53.10 | 76.30 | 5.92 | 8.95 | 8.30 | 14.82 |
| **25 Jul, 2017** | 69.77 | 92.22 | 62.37 | 71.21 | 5.91 | 8.55 | 9.03 | 13.91 |
| **Mean over all** | **46.39** | **73.06** | **40.32** | **65.95** | **4.84** | **8.07** | **6.06** | **13.16** |
| ***P*_0.05_** | **0.000** |  | **0.000** |  | **0.002** |  | **0.000** |  |

| **Table (21). Mean number of *T. urticae* after Egyxide application, Kom Oshim.** | | | | | | | | |
| --- | --- | --- | --- | --- | --- | --- | --- | --- |
|  | ***B. napus*** | **Check** | ***B. oleracea*** | **Check** | ***M. spicata*** | **Check** | ***M. longifolia*** | **Check** |
| **Precount** | **36.30** | **36.45** | **28.21** | **29.25** | **5.57** | **4.26** | **6.42** | **5.80** |
| **04 Apr, 2017** | 26.16 | 41.30 | 20.46 | 32.98 | 2.95 | 4.46 | 4.50 | 6.29 |
| **11 Apr, 2017** | 20.09 | 44.67 | 22.20 | 36.40 | 2.88 | 5.09 | 4.47 | 7.36 |
| **18 Apr, 2017** | 34.25 | 49.23 | 27.49 | 42.00 | 3.33 | 5.82 | 4.75 | 8.66 |
| **25 Apr, 2017** | 44.30 | 53.03 | 39.08 | 46.26 | 3.83 | 5.94 | 6.19 | 9.65 |
| **02 May, 2017** | 50.47 | 56.37 | 47.58 | 51.40 | 4.08 | 6.15 | 6.89 | 10.61 |
| **09 May, 2017** | 25.45 | 61.28 | 38.51 | 58.95 | 2.89 | 6.75 | 4.25 | 12.15 |
| **16 May, 2017** | 31.70 | 65.39 | 38.10 | 65.32 | 2.75 | 7.17 | 4.24 | 13.05 |
| **23 May, 2017** | 48.02 | 71.21 | 41.06 | 70.66 | 3.14 | 9.10 | 4.73 | 14.59 |
| **30 May, 2017** | 57.01 | 79.60 | 65.46 | 78.89 | 3.86 | 9.59 | 5.73 | 15.37 |
| **06 Jun, 2017** | 70.55 | 86.75 | 70.95 | 84.65 | 4.32 | 10.50 | 7.11 | 16.49 |
| **13 Jun, 2017** | 85.27 | 104.60 | 80.44 | 87.95 | 5.28 | 10.98 | 8.12 | 17.60 |
| **20 Jun, 2017** | 34.02 | 102.38 | 44.15 | 87.37 | 5.93 | 11.53 | 9.13 | 18.72 |
| **27 Jun, 2017** | 40.71 | 98.60 | 44.82 | 94.26 | 6.29 | 10.73 | 10.52 | 18.25 |
| **04 Jul, 2017** | 60.87 | 94.50 | 54.06 | 90.65 | 3.75 | 10.05 | 6.90 | 17.00 |
| **11 Jul, 2017** | 66.55 | 89.98 | 61.25 | 82.53 | 4.25 | 9.65 | 7.06 | 16.49 |
| **18 Jul, 2017** | 72.89 | 87.60 | 71.78 | 76.30 | 4.77 | 8.95 | 8.17 | 14.82 |
| **25 Jul, 2017** | 80.30 | 92.22 | 83.05 | 71.21 | 5.48 | 8.55 | 8.40 | 13.91 |
| **Mean over all** | **49.16** | **73.06** | **48.81** | **65.95** | **4.19** | **8.07** | **6.53** | **13.16** |
| ***P*_0.05_** | **0.000** |  | **0.000** |  | **0.002** |  | **0.000** |  |

| **Table (22). Reduction percentage of *P. persimilis* against *T. urticae.*** | | | | |
| --- | --- | --- | --- | --- |
| **Sampling date** | ***B. napus*** | ***B. oleracea*** | ***M. spicata*** | ***M. longifolia*** |
| **04 Apr, 2017** | 47.22 | 34.65 | 14.21 | 30.64 |
| **11 Apr, 2017** | 61.75 | 51.33 | 50.25 | 55.30 |
| **18 Apr, 2017** | 68.48 | 60.10 | 65.03 | 72.01 |
| **25 Apr, 2017** | 75.02 | 73.46 | 79.41 | 82.96 |
| **02 May, 2017** | 82.78 | 84.65 | 81.02 | 90.05 |
| **09 May, 2017** | 89.81 | 92.77 | 86.09 | 97.79 |
| **16 May, 2017** | 95.84 | 95.36 | 89.00 | 98.67 |
| **23 May, 2017** | 98.68 | 98.65 | 91.85 | 99.29 |
| **30 May, 2017** | 99.66 | 100.00 | 94.32 | 99.66 |
| **06 Jun, 2017** | 100.00 | - | 98.05 | 100.00 |
| **13 Jun, 2017** | - | - | 99.43 | - |
| **Mean over all** | **81.93** | **76.77** | **74.92** | **82.64** |
| **L.S.D. at 0.05 = 43.10** | | | | |

| **Table (23). Reduction percentage of *A. swirskii* against *T. urticae.*** | | | | |
| --- | --- | --- | --- | --- |
| **Sampling date** | ***B. napus*** | ***B. oleracea*** | ***M. spicata*** | ***M. longifolia*** |
| **04 Apr, 2017** | 25.99 | 13.73 | 6.04 | 12.56 |
| **11 Apr, 2017** | 40.69 | 24.18 | 21.93 | 28.33 |
| **18 Apr, 2017** | 47.62 | 31.87 | 38.36 | 40.24 |
| **25 Apr, 2017** | 54.28 | 21.49 | 43.77 | 48.19 |
| **02 May, 2017** | 63.09 | 10.54 | 50.97 | 54.05 |
| **09 May, 2017** | 70.02 | 15.47 | 60.48 | 68.31 |
| **16 May, 2017** | 76.63 | 30.09 | 66.57 | 78.93 |
| **23 May, 2017** | 81.40 | 38.04 | 80.88 | 84.58 |
| **30 May, 2017** | 83.68 | 48.77 | 86.70 | 89.26 |
| **13 Jun, 2017** | 85.77 | 52.53 | 91.02 | 98.18 |
| **20 Jun, 2017** | 83.15 | 50.98 | 92.96 | 99.57 |
| **27 Jun, 2017** | 87.60 | 67.16 | 94.23 | 100.00 |
| **04 Jul, 2017** | 89.37 | 72.76 | 98.56 | - |
| **11 Jul, 2017** | 89.66 | 76.46 | 100.00 | - |
| **18 Jul, 2017** | 88.69 | 80.96 | - | - |
| **25 Jul, 2017** | 87.44 | 82.52 | - | - |
| **Mean** | **62.92** | **28.67** | **54.67** | **60.26** |
| **L.S.D. at 0.05 = 24.10** | | | | |

| **Table (24). Reduction percentage of *C. negevi* against *T. urticae.*** | | | | |
| --- | --- | --- | --- | --- |
| **Sampling date** | ***B. napus*** | ***B. oleracea*** | ***M. spicata*** | ***M. longifolia*** |
| **04 Apr, 2017** | 21.12 | 4.27 | 7.84 | 11.32 |
| **11 Apr, 2017** | 33.54 | 13.56 | 24.04 | 26.90 |
| **18 Apr, 2017** | 43.36 | 15.61 | 39.12 | 43.00 |
| **25 Apr, 2017** | 54.04 | 14.98 | 43.26 | 51.66 |
| **02 May, 2017** | 51.96 | 5.82 | 49.04 | 55.80 |
| **09 May, 2017** | 65.02 | 15.97 | 58.70 | 62.42 |
| **16 May, 2017** | 63.43 | 24.15 | 66.06 | 66.52 |
| **23 May, 2017** | 67.32 | 26.48 | 74.56 | 72.93 |
| **30 May, 2017** | 71.02 | 33.14 | 77.34 | 75.75 |
| **13 Jun, 2017** | 71.06 | 36.63 | 82.83 | 78.22 |
| **20 Jun, 2017** | 72.78 | 40.70 | 84.65 | 80.58 |
| **27 Jun, 2017** | 80.17 | 41.90 | 86.34 | 82.00 |
| **04 Jul, 2017** | 82.42 | 51.83 | 88.99 | 81.68 |
| **11 Jul, 2017** | 81.80 | 57.90 | 90.44 | 80.84 |
| **18 Jul, 2017** | 78.01 | 56.03 | 91.19 | 80.69 |
| **25 Jul, 2017** | 78.66 | 58.65 | 94.72 | 79.60 |
| **01 Aug, 2017** | 77.29 | 62.09 | 97.97 | 80.13 |
| **Mean** | **64.29** | **32.92** | **68.06** | **65.30** |
| **L.S.D. at 0.05 = 13.25** | | | | |

| **Table (25). Reduction percentage of Bio-Magic applications against *T. urticae.*** | | | | |
| --- | --- | --- | --- | --- |
| **Sampling date** | ***B. napus*** | ***B. oleracea*** | ***M. spicata*** | ***M. longifolia*** |
| **04 Apr, 2017** | 36.36 | 42.20 | 21.12 | 27.94 |
| **11 Apr, 2017** | 47.72 | 43.17 | 31.82 | 39.49 |
| **18 Apr, 2017** | 46.51 | 36.34 | 39.83 | 37.43 |
| **25 Apr, 2017** | 34.04 | 23.87 | 34.46 | 35.13 |
| **02 May, 2017** | 20.58 | 10.21 | 32.29 | 34.44 |
| **09 May, 2017** | 61.08 | 35.19 | 53.91 | 67.68 |
| **16 May, 2017** | 49.39 | 42.68 | 57.72 | 71.20 |
| **23 May, 2017** | 29.46 | 41.21 | 68.44 | 71.86 |
| **30 May, 2017** | 19.42 | 29.02 | 69.47 | 70.44 |
| **13 Jun, 2017** | 20.67 | 25.49 | 68.47 | 61.70 |
| **20 Jun, 2017** | 19.58 | 20.75 | 65.56 | 50.60 |
| **27 Jun, 2017** | 64.87 | 67.12 | 63.33 | 46.54 |
| **04 Jul, 2017** | 59.69 | 69.60 | 57.10 | 35.02 |
| **11 Jul, 2017** | 46.71 | 56.21 | 51.66 | 54.00 |
| **18 Jul, 2017** | 37.79 | 41.87 | 52.38 | 48.98 |
| **25 Jul, 2017** | 29.27 | 28.37 | 47.23 | 37.17 |
| **01 Aug, 2017** | 24.82 | 9.86 | 44.86 | 27.17 |
| **Mean** | **38.12^c^** | **36.66^c^** | **50.57^a^** | **48.05^b^** |
| **L.S.D. at 0.05 = 13.25** | | | | |

| **Table (26). Reduction percentage of Egyxide applications against *T. urticae.*** | | | | |
| --- | --- | --- | --- | --- |
| **Sampling date** | ***B. napus*** | ***B. oleracea*** | ***M. spicata*** | ***M. longifolia*** |
| **04 Apr, 2017** | 36.40 | 35.68 | 49.41 | 35.37 |
| **11 Apr, 2017** | 54.84 | 36.76 | 56.73 | 45.13 |
| **18 Apr, 2017** | 30.14 | 32.13 | 56.24 | 50.45 |
| **25 Apr, 2017** | 16.12 | 12.41 | 50.69 | 42.05 |
| **02 May, 2017** | 10.10 | 4.02 | 49.26 | 41.33 |
| **09 May, 2017** | 58.30 | 32.27 | 67.25 | 68.40 |
| **16 May, 2017** | 51.32 | 39.52 | 70.67 | 70.65 |
| **23 May, 2017** | 32.29 | 39.75 | 73.61 | 70.71 |
| **30 May, 2017** | 28.08 | 13.96 | 69.22 | 66.32 |
| **13 Jun, 2017** | 18.34 | 13.09 | 68.53 | 61.05 |
| **20 Jun, 2017** | 18.14 | 5.17 | 63.22 | 58.32 |
| **27 Jun, 2017** | 66.63 | 47.60 | 60.66 | 55.94 |
| **04 Jul, 2017** | 58.54 | 50.70 | 55.17 | 47.92 |
| **11 Jul, 2017** | 35.32 | 38.17 | 71.46 | 63.33 |
| **18 Jul, 2017** | 25.73 | 23.05 | 66.32 | 61.32 |
| **25 Jul, 2017** | 16.45 | 2.46 | 59.24 | 50.20 |
| **01 Aug, 2017** | 12.57 | -20.93 | 50.98 | 45.44 |
| **Mean** | **33.49^c^** | **23.87^d^** | **61.10^a^** | **54.94^b^** |
| **L.S.D. at 0.05 = 40.15** | | | | |

| **Table (27). Mean number of *T. tabaci* and *B. tabaci* after Bio-Magic application.** | | | | | | | | |
| --- | --- | --- | --- | --- | --- | --- | --- | --- |
| ***Thrips tabaci*** | | | | | | | | |
|  | ***B. napus*** | **Check** | ***B. oleracea*** | **Check** | ***M. spicata*** | **Check** | ***M. longifolia*** | **Check** |
| **Precount** | 3.23 | 3.30 | **2.95** | 3.70 | 0.62 | 0.15 | 0.80 | 0.42 |
| **1 week** | 6.40 | 5.24 | 7.15 | 6.89 | 0.39 | 0.38 | 0.49 | 0.50 |
| **2 weeks** | 3.00 | 6.60 | 4.66 | 9.40 | 0.35 | 0.60 | 0.47 | 0.79 |
| **3 weeks** | 2.64 | 7.50 | 4.50 | 12.44 | 0.42 | 0.74 | 0.50 | 1.14 |
| **4 weeks** | 4.15 | 8.62 | 7.90 | 14.38 | 0.84 | 1.55 | 1.00 | 1.36 |
| **5 weeks** | 9.72 | 13.20 | 13.81 | 17.26 | 1.10 | 1.63 | 1.19 | 1.68 |
| **6 weeks** | 5.86 | 17.25 | 9.21 | 22.14 | 0.50 | 1.44 | 0.76 | 1.70 |
| **7 weeks** | 5.00 | 20.11 | 8.75 | 25.13 | 0.48 | 1.35 | 0.73 | 1.65 |
| **8 weeks** | 5.10 | 20.27 | 11.00 | 28.19 | 0.57 | 1.33 | 0.93 | 1.56 |
| **9 weeks** | 7.75 | 20.33 | 12.00 | 29.34 | 1.10 | 1.28 | 0.97 | 1.53 |
| **Mean over all** | **5.29** | **12.24** | **8.19** | **16.89** | **0.64** | **1.05** | **0.78** | **1.23** |
| **L.S.D at 0.05 =** | | | | | | | | |
| ***Bemisia tabaci*** | | | | | | | | |
|  | ***B. napus*** | **Check** | ***B. oleracea*** | **Check** | ***M. spicata*** | **Check** | ***M. longifolia*** | **Check** |
| **Precount** | 52.38 | 50.33 | 26.80 | 26.33 | 1.89 | 2.00 | 0.24 | 0.23 |
| **1 week** | 45.12 | 64.82 | 16.20 | 37.10 | 1.80 | 2.13 | 0.16 | 0.98 |
| **2 weeks** | 40.00 | 90.34 | 15.85 | 52.11 | 1.90 | 2.19 | 0.13 | 1.50 |
| **3 weeks** | 38.61 | 116.34 | 20.52 | 70.40 | 2.15 | 2.23 | 0.11 | 1.78 |
| **4 weeks** | 39.70 | 123.20 | 50.46 | 93.11 | 1.96 | 2.28 | 0.15 | 1.94 |
| **5 weeks** | 23.50 | 140.61 | 31.67 | 100.25 | 1.95 | 2.30 | 0.09 | 2.00 |
| **6 weeks** | 16.00 | 156.53 | 30.14 | 111.56 | 2.10 | 2.48 | 0.07 | 2.04 |
| **7 weeks** | 20.01 | 160.24 | 32.21 | 114.20 | 2.23 | 2.64 | 0.10 | 2.03 |
| **8 weeks** | 34.25 | 168.32 | 45.16 | 120.13 | 2.34 | 2.92 | 0.39 | 2.40 |
| **9 weeks** | 41.50 | 172.74 | 60.00 | 119.24 | 2.59 | 3.00 | 0.75 | 2.30 |
| **Mean over all** | **35.11** | **124.35** | **32.90** | **84.44** | **2.09** | **2.42** | **0.22** | **1.72** |
| **L.S.D at 0.05 =** | | | | | | | | |

| **Table (28). Mean number of *T. tabaci* and *B. tabaci* after *A. swirskii* releasing.** | | | | | | | | |
| --- | --- | --- | --- | --- | --- | --- | --- | --- |
| ***Thrips tabaci*** | | | | | | | | |
|  | ***B. napus*** | **Check** | ***B. oleracea*** | **Check** | ***M. spicata*** | **Check** | ***M. longifolia*** | **Check** |
| **Precount** | **1.05** | 3.30 | **1.46** | 3.70 | 0.58 | 0.15 | 0.79 | 0.42 |
| **1 week** | 0.70 | 5.24 | 0.98 | 6.89 | 0.25 | 0.38 | 0.52 | 0.50 |
| **2 weeks** | 0.55 | 6.60 | 0.70 | 9.40 | 0.11 | 0.60 | 0.31 | 0.79 |
| **3 weeks** | 0.50 | 7.50 | 0.51 | 12.44 | 0.05 | 0.74 | 0.19 | 1.14 |
| **4 weeks** | 0.48 | 8.62 | 0.32 | 14.38 | 0.00 | 1.55 | 0.09 | 1.36 |
| **5 weeks** | 0.21 | 13.20 | 0.17 | 17.26 |  | 1.63 | 0.01 | 1.68 |
| **6 weeks** | 0.10 | 17.25 | 0.08 | 22.14 |  | 1.44 |  | 1.70 |
| **7 weeks** | 0.00 | 20.11 | 0.00 | 25.13 |  | 1.35 |  | 1.65 |
| **Mean over all** | **0.45** | **10.23** | **0.53** | **13.92** | **0.20** | **0.98** | **0.32** | **1.16** |
| **L.S.D at 0.05 = 28.35** | | | | | | | | |
| ***Bemisia tabaci*** | | | | | | | | |
|  | ***B. napus*** | **Check** | ***B. oleracea*** | **Check** | ***M. spicata*** | **Check** | ***M. longifolia*** | **Check** |
| **Precount** | 49.01 | 50.33 | 27.00 | 26.33 | 1.90 | 2.00 | 0.25 | 0.23 |
| **1 week** | 40.07 | 64.82 | 23.11 | 37.10 | 1.10 | 2.13 | 0.13 | 0.98 |
| **2 weeks** | 31.85 | 90.34 | 18.20 | 52.11 | 0.90 | 2.19 | 0.10 | 1.50 |
| **3 weeks** | 25.4 | 116.34 | 9.50 | 70.40 | 0.63 | 2.23 | 0.09 | 1.78 |
| **4 weeks** | 19.31 | 123.20 | 8.90 | 93.11 | 0.29 | 2.28 | 0.05 | 1.94 |
| **5 weeks** | 12.05 | 140.61 | 5.31 | 100.25 | 0.21 | 2.30 | 0.01 | 2.00 |
| **6 weeks** | 6.41 | 156.53 | 2.00 | 111.56 | 0.17 | 2.48 | 0.00 | 2.04 |
| **7 weeks** | 2.01 | 160.24 | 0.05 | 114.20 | 0.12 | 2.64 | 0.00 | 2.03 |
| **8 weeks** | 0.20 | 168.32 | 0.00 | 120.13 | 0.08 | 2.92 | 0.00 | 2.40 |
| **Mean over all** | **20.70** | **118.97** | **10.45** | **80.58** | **0.60** | **2.35** | **0.11** | **1.66** |
| **L.S.D at 0.05 = 12.08** | | | | | | | | |

| **Table (29). Reduction percentage of *A. swirskii* against *T. tabaci.*** | | | | |
| --- | --- | --- | --- | --- |
| **Sampling date** | ***B. napus*** | ***B. oleracea*** | ***M. spicata*** | ***M. longifolia*** |
| **04 April, 2017** | 58.02 | 63.95 | 82.99 | 44.71 |
| **11 April, 2017** | 73.81 | 81.13 | 53.54 | 79.14 |
| **18 April, 2017** | 79.05 | 89.61 | 82.88 | 91.14 |
| **25 April, 2017** | 82.50 | 94.36 | 100.00 | 96.48 |
| **02 May, 2017** | 95.00 | 97.50 |  | 99.68 |
| **09 May, 2017** | 98.18 | 99.08 |  |  |
| **16 May, 2017** | 100.00 | 100.00 |  |  |
| **Mean over all** | **83.79** | **89.38** | **79.85** | **82.23** |
| **L.S.D. at 0.05 = 6.05** | | | | |

| **Table (30). Mean number of *T. tabaci* and *B. tabaci* after Egyxide application.** | | | | | | | | |
| --- | --- | --- | --- | --- | --- | --- | --- | --- |
| ***Thrips tabaci*** | | | | | | | | |
|  | ***B. napus*** | **Check** | ***B. oleracea*** | **Check** | ***M. spicata*** | **Check** | ***M. longifolia*** | **Check** |
| **Precount** | 3.23 | 3.30 | **2.95** | 3.70 | 0.60 | 0.15 | 0.78 | 0.42 |
| **1 week** | 6.40 | 5.24 | 7.15 | 6.89 | 0.44 | 0.38 | 0.52 | 0.50 |
| **2 weeks** | 3.00 | 6.60 | 4.66 | 9.40 | 0.42 | 0.60 | 0.50 | 0.79 |
| **3 weeks** | 2.64 | 7.50 | 4.50 | 12.44 | 0.49 | 0.74 | 0.83 | 1.14 |
| **4 weeks** | 4.15 | 8.62 | 7.90 | 14.38 | 0.79 | 1.55 | 1.10 | 1.36 |
| **5 weeks** | 9.72 | 13.20 | 13.81 | 17.26 | 1.00 | 1.63 | 1.05 | 1.68 |
| **6 weeks** | 5.86 | 17.25 | 9.21 | 22.14 | 0.68 | 1.44 | 0.85 | 1.70 |
| **7 weeks** | 5.00 | 20.11 | 8.75 | 25.13 | 0.65 | 1.35 | 0.81 | 1.65 |
| **8 weeks** | 5.10 | 20.27 | 11.00 | 28.19 | 0.67 | 1.33 | 0.93 | 1.56 |
| **9 weeks** | 0.65 | 20.33 | 0.81 | 29.34 | 0.75 | 1.28 | 0.97 | 1.53 |
| **Mean over all** | **4.58** | **12.24** | **7.07** | **16.89** | **0.65** | **1.05** | **0.83** | **1.23** |
| **L.S.D at 0.05 = 19.45** | | | | | | | | |
| ***Bemisia tabaci*** | | | | | | | | |
|  | ***B. napus*** | **Check** | ***B. oleracea*** | **Check** | ***M. spicata*** | **Check** | ***M. longifolia*** | **Check** |
| **Precount** | 49.01 | 50.33 | 27.44 | 26.33 | 2.15 | 2.00 | 0.22 | 0.23 |
| **1 week** | 35.71 | 64.82 | 13.99 | 37.10 | 1.66 | 2.13 | 0.10 | 0.98 |
| **2 weeks** | 29.52 | 90.34 | 13.25 | 52.11 | 1.58 | 2.19 | 0.09 | 1.50 |
| **3 weeks** | 31.00 | 116.34 | 16.81 | 70.40 | 1.60 | 2.23 | 0.12 | 1.78 |
| **4 weeks** | 48.16 | 123.20 | 43.12 | 93.11 | 1.79 | 2.28 | 0.23 | 1.94 |
| **5 weeks** | 29.11 | 140.61 | 29.64 | 100.25 | 2.00 | 2.30 | 0.11 | 2.00 |
| **6 weeks** | 27.6 | 156.53 | 27.54 | 111.56 | 1.85 | 2.48 | 0.10 | 2.04 |
| **7 weeks** | 33.98 | 160.24 | 34.30 | 114.20 | 1.89 | 2.64 | 0.19 | 2.03 |
| **8 weeks** | 45.00 | 168.32 | 51.74 | 120.13 | 1.32 | 2.92 | 0.31 | 2.40 |
| **9 weeks** | 65.81 | 172.74 | 72.31 | 119.24 | 1.76 | 3.00 | 0.69 | 2.30 |
| **Mean over all** | **39.49** | **124.35** | **33.01** | **84.44** | **1.76** | **2.42** | **0.22** | **1.72** |
| **L.S.D at 0.05 = 24.19** | | | | | | | | |

| **Table (31). Reduction percentage of Bio-Magic against *T. tabaci.*** | | | | |
| --- | --- | --- | --- | --- |
| **Sampling date** | ***B. napus*** | ***B. oleracea*** | ***M. spicata*** | ***M. longifolia*** |
| **04 April, 2017** | -22.14 | -30.16 | 75.17 | 48.55 |
| **11 April, 2017** | 54.55 | 37.82 | 85.89 | 68.77 |
| **18 April, 2017** | 64.80 | 54.63 | 86.27 | 76.97 |
| **25 April, 2017** | 51.86 | 31.10 | 86.89 | 61.40 |
| **02 May, 2017** | 26.36 | -0.35 | 83.67 | 62.81 |
| **09 May, 2017** | 66.03 | 47.83 | 91.60 | 76.53 |
| **16 May, 2017** | 75.14 | 56.33 | 91.40 | 76.77 |
| **23 May, 2017** | 74.84 | 51.06 | 89.63 | 68.70 |
| **30 May, 2017** | 61.88 | 48.70 | 79.21 | 66.72 |
| **Mean over all** | **50.37^c^** | **32.99^d^** | **85.52^a^** | **67.47^b^** |
| **L.S.D. at 0.05 = 45.35** | | | | |

| **Table (32). Reduction percentage of *A. swirskii* applications against *B. tabaci.*** | | | | |
| --- | --- | --- | --- | --- |
| **Sampling date** | ***B. napus*** | ***B. oleracea*** | ***M. spicata*** | ***M. longifolia*** |
| **04 April, 2017** | 36.52 | 39.25 | 45.64 | 87.80 |
| **11 April, 2017** | 63.79 | 65.94 | 56.74 | 93.87 |
| **18 April, 2017** | 77.58 | 86.84 | 70.26 | 95.35 |
| **25 April, 2017** | 83.90 | 90.68 | 86.61 | 97.63 |
| **02 May, 2017** | 91.20 | 94.83 | 90.39 | 99.54 |
| **09 May, 2017** | 95.79 | 98.25 | 92.78 | 100.00 |
| **16 May, 2017** | 98.71 | 99.96 | 95.22 |  |
| **23 May, 2017** | 99.88 | 100.00 | 97.12 |  |
| **Mean over all** | **80.92^b^** | **84.47^b^** | **79.34^c^** | **95.70^a^** |
| **L.S.D. at 0.05 = 15.30** | | | | |

| **Table (33). Mean number of *T. tabaci* and *B. tabaci* after *C. negevi* releasing.** | | | | | | | | |
| --- | --- | --- | --- | --- | --- | --- | --- | --- |
| ***Thrips tabaci*** | | | | | | | | |
|  | ***B. napus*** | **Check** | ***B. oleracea*** | **Check** | ***M. spicata*** | **Check** | ***M. longifolia*** | **Check** |
| **Precount** | **2.15** | **2.11** | **1.46** | **1.78** | **0.61** | **0.15** | **0.81** | **0.42** |
| **1 week** | 1.97 | 2.95 | 0.98 | 2.00 | 0.77 | 0.38 | 1.10 | 0.5 |
| **2 weeks** | 3.00 | 3.30 | 0.70 | 3.75 | 1.63 | 0.6 | 1.33 | 0.79 |
| **3 weeks** | 4.21 | 5.24 | 0.51 | 6.89 | 1.70 | 0.74 | 1.60 | 1.14 |
| **4 weeks** | 6.13 | 6.61 | 0.32 | 9.47 | 1.88 | 1.55 | 1.71 | 1.36 |
| **5 weeks** | 7.00 | 7.50 | 0.17 | 12.44 | 2.03 | 1.63 | 1.67 | 1.68 |
| **6 weeks** | 7.88 | 8.62 | 0.08 | 14.38 | 2.14 | 1.44 | 1.54 | 1.7 |
| **7 weeks** | 9.90 | 13.20 | 0.00 | 17.26 | 2.30 | 1.35 | 1.49 | 1.65 |
| **8 weeks** | 13.40 | 17.25 |  | 22.14 | 2.55 | 1.33 | 1.56 | 1.56 |
| **9 weeks** | 18.64 | 20.11 |  | 25.13 | 2.86 | 1.28 | 1.74 | 1.53 |
| **Mean over all** | **7.33** | **8.69** | **0.53** | **11.51** | **1.85** | **1.05** | **1.46** | **1.23** |
| **L.S.D at 0.05 = 16.99** | | | | | | | | |
| ***Bemisia tabaci*** | | | | | | | | |
|  | ***B. napus*** | **Check** | ***B. oleracea*** | **Check** | ***M. spicata*** | **Check** | ***M. longifolia*** | **Check** |
| **Precount** | 51.64 | 50.33 | 25.30 | 26.33 | 2.10 | 2.00 | 0.24 | 0.23 |
| **1 week** | 44.32 | 64.82 | 20.05 | 37.10 | 1.75 | 2.13 | 0.20 | 0.98 |
| **2 weeks** | 35.40 | 90.34 | 17.34 | 52.11 | 1.59 | 2.19 | 0.16 | 1.50 |
| **3 weeks** | 29.33 | 116.34 | 16.00 | 70.40 | 1.00 | 2.23 | 0.14 | 1.78 |
| **4 weeks** | 26.20 | 123.20 | 15.59 | 93.11 | 0.80 | 2.28 | 0.12 | 1.94 |
| **5 weeks** | 13.10 | 140.61 | 11.40 | 100.25 | 0.64 | 2.30 | 0.09 | 2.00 |
| **6 weeks** | 8.05 | 156.53 | 9.00 | 111.56 | 0.44 | 2.48 | 0.06 | 2.04 |
| **7 weeks** | 4.66 | 160.24 | 7.05 | 114.20 | 0.37 | 2.64 | 0.06 | 2.03 |
| **8 weeks** | 1.00 | 168.32 | 3.12 | 120.13 | 0.25 | 2.92 | 0.03 | 2.40 |
| **9 weeks** | 0.05 | 175.14 | 1.00 | 128.25 | 0.19 | 3.09 | 0.00 | 2.37 |
| **10 weeks** | 0.00 | 180.82 | 0.22 | 135.05 | 0.00 | 3.29 | 0.00 | 2.47 |
| **Mean over all** | **19.43** | **129.70** | **11.46** | **89.86** | **0.83** | **2.50** | **0.10** | **1.79** |
| **L.S.D at 0.05 = 19.54** | | | | | | | | |

| **Table (34). Reduction percentage of *C. negevi* applications against *T. tabaci and B. tabaci.*** | | | | | | | | | | | |
| --- | --- | --- | --- | --- | --- | --- | --- | --- | --- | --- | --- |
|  | ***Thrips tabaci*** | | | | ***Bemisia tabaci*** | | | | | | |
| **Sampling date** | ***B. napus*** | ***B. oleracea*** | ***M. spicata*** | ***M. longifolia*** | ***B. napus*** | | ***B. oleracea*** | | ***M. spicata*** | | ***M. longifolia*** |
| **02 May, 2017** | 34.46 | 40.26 | 50.17 | -14.07 | 33.36 | | 43.76 | | 91.06 | | 80.44 |
| **09 May, 2017** | 10.78 | 76.93 | 33.20 | 12.71 | 61.81 | | 65.37 | | 93.04 | | 89.84 |
| **16 May, 2017** | 21.15 | 90.98 | 43.51 | 27.23 | 75.43 | | 76.35 | | 94.02 | | 92.51 |
| **23 May, 2017** | 8.85 | 95.85 | 70.17 | 34.80 | 79.27 | | 82.57 | | 94.99 | | 94.11 |
| **30 May, 2017** | 8.40 | 98.33 | 69.38 | 48.46 | 90.92 | | 88.17 | | 96.27 | | 95.71 |
| **13 June, 2017** | 10.29 | 99.32 | 63.46 | 53.03 | 94.99 | | 91.60 | | 97.70 | | 97.20 |
| **20 June, 2017** | 26.40 | 100.00 | 58.11 | 53.18 | 97.17 | | 93.58 | | 97.84 | | 97.19 |
| **27 June, 2017** | 23.76 |  | 52.85 | 48.15 | 99.42 | | 97.30 | | 99.02 | | 98.81 |
| **04 July, 2017** | 9.03 |  | 45.06 | 41.03 | 100.00 | | 99.19 | | 100.00 | | 100.00 |
| **11 July, 2017** |  |  |  |  |  | | 99.83 | |  | |  |
| **Mean** | **17.01** | **85.95** | **53.99** | **33.83** | **81.37** | | **81.99** | | **95.99** | | **93.98** |
| **L.S.D. at 0.05 = 33.71** | | | | | |  | |  | |  | |

a) P. persimilis releasing results on immatures of the TSSM

|  | Brassica napus var. pabularia | Brassica oleracea var palmifolia | Mentha spicata | Mentha longifolia |
| --- | --- | --- | --- | --- |
| 2017-04-04 | 20.53 | 16.23 | 2 | 2.4 |
| 2017-04-11 | 10 | 14.46 | 1.65 | 2.02 |
| 2017-04-18 | 8 | 11 | 1.22 | 1.3 |
| 2017-04-25 | 7 | 9.77 | 0.98 | 1.15 |
| 2017-05-02 | 5.36 | 6.3 | 0.14 | 1 |
| 2017-05-09 | 2.5 | 3.5 | 0.09 | 0.77 |
| 2017-05-16 | 1 | 1.26 | 0 | 0.31 |
| 2017-05-23 | 0.45 | 0.8 | 0 | 0.2 |
| 2017-05-30 | 0 | 0 | 0 | 0.12 |
| 2017-06-06 | 0 | 0 | 0 | 0.06 |
| 2017-06-13 | 0 | 0 | 0 | 0 |

**Fig. (10 a & b). Releasing results of of *P. persmilis* against the TSSM associated with Brassicaceae and Lamicaceae medicinal plants in Kom Oshim, season 2017.**

b) P. persimilis releasing results on adults of the TSSM

|  | Brassica napus var. pabularia | Brassica oleracea var palmifolia | Mentha spicata | Mentha longifolia |
| --- | --- | --- | --- | --- |
| 2017-04-04 | 18.3 | 11.4 | 3.4 | 4.3 |
| 2017-04-11 | 13.22 | 6 | 3.2 | 3.02 |
| 2017-04-18 | 10.2 | 5.48 | 1.99 | 2.5 |
| 2017-04-25 | 9.53 | 5.12 | 1.6 | 1.65 |
| 2017-05-02 | 8.75 | 4.37 | 1.41 | 0.9 |
| 2017-05-09 | 7.84 | 3.06 | 1.39 | 0.45 |
| 2017-05-16 | 5.65 | 2.1 | 1.19 | 0 |
| 2017-05-23 | 2.45 | 1.5 | 1 |  |
| 2017-05-30 | 1 | 0.73 | 0.94 |  |
| 2017-06-06 | 0.29 |  | 0.69 |  |
| 2017-06-13 |  |  | 0.26 |  |
| 2017-06-20 |  |  | 0.08 |  |

a) A. swirskii released on T. urticae immatures

|  | Brassica napus var. pabularia | Brassica oleracea var palmifolia | Mentha spicata | Mentha longifolia |
| --- | --- | --- | --- | --- |
| 2017-04-04 | 20.11 | 15.2 | 2.01 | 2.32 |
| 2017-04-11 | 15.25 | 14 | 1.98 | 2.2 |
| 2017-04-18 | 13 | 13.1 | 1.75 | 2.11 |
| 2017-04-25 | 13.1 | 11.64 | 1.44 | 2.07 |
| 2017-05-02 | 12.6 | 10.19 | 1.2 | 2 |
| 2017-05-09 | 10.9 | 9.21 | 0.8 | 1.95 |
| 2017-05-16 | 8.75 | 16.54 | 0.45 | 1.54 |
| 2017-05-23 | 7 | 14.6 | 0.32 | 1.1 |
| 2017-05-30 | 6.4 | 12.81 | 0.2 | 0.9 |
| 2017-06-06 | 5.9 | 10.75 |  | 0.66 |
| 2017-06-13 | 5.4 | 10.06 |  | 0.12 |
| 2017-06-20 | 8.9 | 15 |  | 0.03 |
| 2017-06-27 | 6.75 | 10 |  |  |
| 2017-07-04 | 6 | 9 |  |  |
| 2017-07-11 | 5.45 | 9 |  |  |
| 2017-07-18 | 5.24 | 5 |  |  |
| 2017-07-25 | 5 | 5 |  |  |
| 2017-08-01 | 4.9 | 4 |  |  |

**Fig. (11 a & b). Releasing results of of *A. swirskii* against the TSSM associated with Brassicaceae and Lamicaceae medicinal plants in Kom Oshim, season 2017.**

b) A. swirskii released on T. urticae adults

|  | Brassica napus var. pabularia | Brassica oleracea var palmifolia | Mentha spicata | Mentha longifolia |
| --- | --- | --- | --- | --- |
| 2017-04-04 | 16.5 | 11.53 | 3.5 | 2.32 |
| 2017-04-11 | 15.45 | 12 | 3.44 | 2.2 |
| 2017-04-18 | 13.61 | 12.12 | 3.39 | 2.11 |
| 2017-04-25 | 12.8 | 14.51 | 3.2 | 2.07 |
| 2017-05-02 | 11.75 | 23 | 3.12 | 2 |
| 2017-05-09 | 10 | 32.81 | 3.1 | 1.95 |
| 2017-05-16 | 9.7 | 29 | 3 | 1.54 |
| 2017-05-23 | 8.35 | 27.13 | 2.78 | 1.1 |
| 2017-05-30 | 6.9 | 27.2 | 2.05 | 0.9 |
| 2017-06-06 | 7.15 | 26.18 | 1.65 | 0.66 |
| 2017-06-13 | 7 | 26.66 | 1.22 | 0.12 |
| 2017-06-20 | 8.8 | 24.4 | 1 | 0.03 |
| 2017-06-27 | 6 | 16.22 | 0.86 |  |
| 2017-07-04 | 4.53 | 14.46 | 0.2 |  |
| 2017-07-11 | 4.36 | 10.5 |  |  |
| 2017-07-18 | 4.98 | 9.36 |  |  |
| 2017-07-25 | 6.05 | 7.19 |  |  |
| 2017-08-01 | 7.21 | 5.23 |  |  |

a) A. swirskii releasing results on T. tabaci

|  | Brassica napus var. pabularia | Brassica oleracea var palmifolia | Mentha spicata | Mentha longifolia |
| --- | --- | --- | --- | --- |
| 2017-05-09 | 1.05 | 1.46 | 0.58 | 0.79 |
| 2017-05-16 | 0.7 | 0.98 | 0.25 | 0.52 |
| 2017-05-23 | 0.55 | 0.7 | 0.11 | 0.31 |
| 2017-05-30 | 0.5 | 0.51 | 0.05 | 0.19 |
| 2017-06-06 | 0.48 | 0.32 | 0 | 0.09 |
| 2017-06-13 | 0.21 | 0.17 |  | 0.01 |
| 2017-06-20 | 0.1 | 0.08 |  |  |
| 2017-06-27 | 0 | 0 |  |  |

**Fig. (12 a & b). Releasing results of of *A. swirskii* against the active stages of *T. tabaci* and *B. tabaci* associated with Brassicaceae and Lamicaceae medicinal plants in Kom Oshim, 2017**

b) A. swirskii releasing results on B. tabaci

|  | Brassica napus var. pabularia | Brassica oleracea var palmifolia | Mentha spicata | Mentha longifolia |
| --- | --- | --- | --- | --- |
| 2017-05-09 | 49.01 | 27 | 1.9 | 0.25 |
| 2017-05-16 | 40.07 | 23.11 | 1.1 | 0.13 |
| 2017-05-23 | 31.85 | 18.2 | 0.9 | 0.1 |
| 2017-05-30 | 25.4 | 9.5 | 0.63 | 0.09 |
| 2017-06-06 | 19.31 | 8.9 | 0.29 | 0.05 |
| 2017-06-13 | 12.05 | 5.31 | 0.21 | 0.01 |
| 2017-06-20 | 6.41 | 2 | 0.17 | 0 |
| 2017-06-27 | 2.01 | 0.05 | 0.12 | 0 |
| 2017-07-04 | 0.2 | 0 | 0.08 | 0 |
| 2017-07-11 | 0 | 0 | 0.01 | 0 |

a) C. negevi against TSSM immatures

|  | Brassica napus var. pabularia | Brassica oleracea var palmifolia | Mentha spicata | Mentha longifolia |
| --- | --- | --- | --- | --- |
| 2017-04-04 | 19.54 | 14.99 | 1.98 | 2.35 |
| 2017-04-11 | 18.2 | 13.81 | 1.83 | 2.26 |
| 2017-04-18 | 16.15 | 12.55 | 1.65 | 2.18 |
| 2017-04-25 | 14.8 | 11.34 | 1.33 | 2 |
| 2017-05-02 | 10.15 | 10 | 1.18 | 1.89 |
| 2017-05-09 | 11 | 11.45 | 1 | 1.9 |
| 2017-05-16 | 9 | 14 | 0.87 | 1.85 |
| 2017-05-23 | 12.45 | 15.21 | 0.64 | 1.77 |
| 2017-05-30 | 12.8 | 15.3 | 0.54 | 1.6 |
| 2017-06-06 | 11.75 | 14.98 | 0.5 | 1.6 |
| 2017-06-13 | 12 | 14.5 | 0.29 | 1.53 |
| 2017-06-20 | 13.7 | 12.31 | 0.25 | 1.42 |
| 2017-06-27 | 8.35 | 12.2 | 0.25 | 1.38 |
| 2017-07-04 | 6.9 | 12.45 | 0.2 | 1.35 |
| 2017-07-11 | 7.15 | 10.82 | 0.1 | 1.35 |
| 2017-07-18 | 7 | 10.5 | 0.1 | 1.36 |
| 2017-07-25 | 5 | 7.79 | 0.1 | 1.29 |
| 2017-08-01 | 4.9 | 5 | 0.1 | 1.22 |

b) C. negevi against TSSM adults

|  | Brassica napus var. pabularia | Brassica oleracea var palmifolia | Mentha spicata | Mentha longifolia |
| --- | --- | --- | --- | --- |
| 2017-04-04 | 16.8 | 11.7 | 3.43 | 2.35 |
| 2017-04-11 | 14.28 | 15 | 3.39 | 2.26 |
| 2017-04-18 | 13.45 | 16.16 | 3.26 | 2.18 |
| 2017-04-25 | 13 | 21 | 3.17 | 2 |
| 2017-05-02 | 14.15 | 25.89 | 3.1 | 1.89 |
| 2017-05-09 | 16 | 32.72 | 2.98 | 1.9 |
| 2017-05-16 | 12.37 | 31.2 | 2.67 | 1.85 |
| 2017-05-23 | 11.39 | 30 | 2.45 | 1.77 |
| 2017-05-30 | 10.4 | 32.1 | 2.4 | 1.6 |
| 2017-06-06 | 11.25 | 33.15 | 2.26 | 1.42 |
| 2017-06-13 | 13.03 | 34.45 | 2 | 1.38 |
| 2017-06-20 | 14.69 | 35.28 | 1.89 | 1.35 |
| 2017-06-27 | 11.89 | 34.12 | 1.75 | 1.35 |
| 2017-07-04 | 10.38 | 28.98 | 1.3 | 1.36 |
| 2017-07-11 | 10 | 24 | 1.12 | 1.29 |
| 2017-07-18 | 12.73 | 22.61 | 0.98 | 1.22 |
| 2017-07-25 | 13.64 | 21 | 0.5 | 1.16 |
| 2017-08-01 | 15.98 | 19.63 | 0.12 | 1.02 |

**Fig. (13 a & b). Releasing results of of *C. negevi* against the TSSM associated with Brassicaceae and Lamicaceae medicinal plants in Kom Oshim 2017.**

**Fig. (14 a & b). Releasing results of of *C. negevi* against *T. tabaci* and *B. tabaci* associated with Brassicaceae and Lamicaceae medicinal plants in Kom Oshim, 2017.**

a) C. negevi releasing results on T. tabaci

|  | Brassica napus var. pabularia | Brassica oleracea var palmifolia | Mentha spicata | Mentha longifolia |
| --- | --- | --- | --- | --- |
| 2017-05-02 | 1.15 | 1.6 | 0.61 | 0.81 |
| 2017-05-09 | 1.97 | 2 | 0.77 | 1.1 |
| 2017-05-16 | 3 | 2.13 | 1.63 | 1.33 |
| 2017-05-23 | 4.21 | 3.88 | 1.7 | 1.6 |
| 2017-05-30 | 6.13 | 7.2 | 1.88 | 1.71 |
| 2017-06-06 | 7 | 9.94 | 2.03 | 1.67 |
| 2017-06-13 | 7.88 | 12.69 | 2.14 | 1.54 |
| 2017-06-20 | 9.9 | 15 | 2.3 | 1.49 |
| 2017-06-27 | 13.4 | 17.48 | 2.55 | 1.56 |
| 2017-07-04 | 18.64 | 21.65 | 2.86 | 1.74 |

b) C. negevi releasing results on B. tabaci

|  | Brassica napus var. pabularia | Brassica oleracea var palmifolia | Mentha spicata | Mentha longifolia |
| --- | --- | --- | --- | --- |
| 2017-05-02 | 51.64 | 25.3 | 2.1 | 0.24 |
| 2017-05-09 | 44.32 | 20.05 | 1.75 | 0.2 |
| 2017-05-16 | 35.4 | 17.34 | 1.59 | 0.16 |
| 2017-05-23 | 29.33 | 16 | 1 | 0.14 |
| 2017-05-30 | 26.2 | 15.59 | 0.8 | 0.12 |
| 2017-06-06 | 13.1 | 11.4 | 0.64 | 0.09 |
| 2017-06-13 | 8.05 | 9 | 0.44 | 0.06 |
| 2017-06-20 | 4.66 | 7.05 | 0.37 | 0.06 |
| 2017-06-27 | 1 | 3.12 | 0.25 | 0.03 |
| 2017-07-04 | 0.05 | 1 | 0.19 | 0 |
| 2017-07-11 | 0 | 0.22 | 0 | 0 |

a) Bio-Magic application on TSSM immatures

|  | Brassica napus var. pabularia | Brassica oleracea var palmifolia | Mentha spicata | Mentha longifolia |
| --- | --- | --- | --- | --- |
| 2017-04-04 | 19.68 | 17 | 1.89 | 2.17 |
| 2017-04-11 | 14.25 | 11.52 | 0.9 | 1.2 |
| 2017-04-18 | 12.5 | 11.29 | 0.93 | 1.18 |
| 2017-04-25 | 13 | 12.64 | 1.05 | 2 |
| 2017-05-02 | 18.85 | 13.72 | 1.13 | 2.59 |
| 2017-05-09 | 25.6 | 16.84 | 1.27 | 3.1 |
| 2017-05-16 | 13.5 | 10.42 | 0.51 | 1 |
| 2017-05-23 | 21.3 | 10.38 | 0.48 | 0.9 |
| 2017-05-30 | 32.9 | 11.48 | 0.45 | 0.97 |
| 2017-06-06 | 38.75 | 24.5 | 0.67 | 1.1 |
| 2017-06-13 | 40 | 29.66 | 0.89 | 2.43 |
| 2017-06-20 | 47.25 | 33.84 | 1.15 | 3.87 |
| 2017-06-27 | 17.9 | 13.41 | 1.48 | 4.6 |
| 2017-07-04 | 22.45 | 13.36 | 1.77 | 5.32 |
| 2017-07-11 | 30.25 | 16.67 | 1.74 | 2.97 |
| 2017-07-18 | 34.15 | 19.51 | 1.55 | 2.75 |
| 2017-07-25 | 36.75 | 25.1 | 2.03 | 3 |
| 2017-08-01 | 39.8 | 33.47 | 2.15 | 3.11 |

**Fig. (15 a & b). Application results of of Bio-Magic against the TSSM associated with Brassicaceae and Lamicaceae medicinal plants in Kom Oshim, 2017.**

b) Bio-Magic application on TSSM adults

|  | Brassica napus var. pabularia | Brassica oleracea var palmifolia | Mentha spicata | Mentha longifolia |
| --- | --- | --- | --- | --- |
| 2017-04-04 | 17 | 11.42 | 3.45 | 3 |
| 2017-04-11 | 12.2 | 7 | 3.51 | 2.84 |
| 2017-04-18 | 11 | 8.81 | 3.42 | 2.79 |
| 2017-04-25 | 13.5 | 13.34 | 3.34 | 2.83 |
| 2017-05-02 | 16.35 | 20.5 | 3.75 | 2.99 |
| 2017-05-09 | 19.45 | 28 | 3.95 | 3.1 |
| 2017-05-16 | 10.5 | 26.7 | 3.39 | 2.5 |
| 2017-05-23 | 12 | 26 | 3.32 | 2.45 |
| 2017-05-30 | 17.65 | 28.88 | 3.15 | 2.69 |
| 2017-06-06 | 25.8 | 29.91 | 3 | 2.95 |
| 2017-06-13 | 29.25 | 31.62 | 3.26 | 3.2 |
| 2017-06-20 | 37.4 | 33.88 | 3.59 | 3.88 |
| 2017-06-27 | 18.29 | 14.5 | 3.82 | 4.32 |
| 2017-07-04 | 17.55 | 14.48 | 4 | 5.25 |
| 2017-07-11 | 20.43 | 21.9 | 4.35 | 4 |
| 2017-07-18 | 22.18 | 27.1 | 4.21 | 4.75 |
| 2017-07-25 | 25.6 | 28 | 3.89 | 5.3 |
| 2017-08-01 | 29.97 | 28.9 | 3.76 | 5.92 |

a) Bio-Magic on T. tabaci

|  | Brassica napus var. pabularia | Brassica oleracea var palmifolia | Mentha spicata | Mentha longifolia |
| --- | --- | --- | --- | --- |
| 2017-05-02 | 1.08 | 1.51 | 0.62 | 0.8 |
| 2017-05-09 | 1.9 | 1.8 | 0.39 | 0.49 |
| 2017-05-16 | 1 | 1.16 | 0.35 | 0.47 |
| 2017-05-23 | 0.95 | 1.1 | 0.42 | 0.5 |
| 2017-05-30 | 3.23 | 2.95 | 0.84 | 1 |
| 2017-06-06 | 6.4 | 7.15 | 1.1 | 1.19 |
| 2017-06-13 | 3 | 4.66 | 0.5 | 0.76 |
| 2017-06-20 | 2.64 | 4.5 | 0.48 | 0.73 |
| 2017-06-27 | 4.15 | 7.9 |  |  |
| 2017-07-04 | 9.72 | 13.81 |  |  |
| 2017-07-11 | 5.86 | 9.21 |  |  |
| 2017-07-18 | 5 | 8.75 |  |  |
| 2017-07-25 | 5.1 | 11 |  |  |

**Fig. (16 a & b). Application results of Bio-Magic against *T. tabaci* and *B. tabaci* associated with Brassicaceae and Lamicaceae medicinal plants in Kom Oshim, 2017.**

b) Bio-Magic on B. tabaci

|  | Brassica napus var. pabularia | Brassica oleracea var palmifolia | Mentha spicata | Mentha longifolia |
| --- | --- | --- | --- | --- |
| 2017-05-09 | 52.38 | 26.8 | 1.89 | 0.24 |
| 2017-05-16 | 45.12 | 16.2 | 1.8 | 0.16 |
| 2017-05-23 | 40 | 15.85 | 1.9 | 0.13 |
| 2017-05-30 | 38.61 | 20.52 | 2.15 | 0.11 |
| 2017-06-06 | 39.7 | 50.46 | 1.96 | 0.15 |
| 2017-06-13 | 23.5 | 31.67 | 1.95 | 0.09 |
| 2017-06-20 | 16 | 30.14 | 2.1 | 0.07 |
| 2017-06-27 | 20.01 | 32.21 | 2.23 | 0.1 |
| 2017-07-04 | 34.25 | 45.16 | 2.34 | 0.39 |
| 2017-07-11 | 41.5 | 60 | 2.59 | 0.75 |

a) Egyxide application on the TSSM immatures

|  | Brassica napus var. pabularia | Brassica oleracea var palmifolia | Mentha spicata | Mentha longifolia |
| --- | --- | --- | --- | --- |
| 2017-04-04 | 19.5 | 16.86 | 2 | 2.71 |
| 2017-04-11 | 9.36 | 10.34 | 0.5 | 1 |
| 2017-04-18 | 10.45 | 10.2 | 0.48 | 1.01 |
| 2017-04-25 | 16.75 | 11 | 0.69 | 1.21 |
| 2017-05-02 | 23.6 | 12.36 | 0.94 | 2.3 |
| 2017-05-09 | 27.22 | 14.94 | 1.08 | 2.89 |
| 2017-05-16 | 10.45 | 9 | 0.2 | 0.5 |
| 2017-05-23 | 17.08 | 9.35 | 0.17 | 0.47 |
| 2017-05-30 | 22.9 | 10.88 | 0.35 | 0.73 |
| 2017-06-06 | 26.55 | 32.49 | 0.88 | 1.38 |
| 2017-06-13 | 32.45 | 34.95 | 1.2 | 2.27 |
| 2017-06-20 | 40.15 | 40 | 1.95 | 3.12 |
| 2017-06-27 | 15 | 19.05 | 2.07 | 3.9 |
| 2017-07-04 | 20.15 | 18.22 | 2.14 | 4.87 |
| 2017-07-11 | 33.75 | 22.5 | 0.35 | 2 |
| 2017-07-18 | 36.55 | 26.1 | 0.96 | 2.1 |
| 2017-07-25 | 38.04 | 33.78 | 1.25 | 3.05 |
| 2017-08-01 | 40.8 | 43.96 | 1.78 | 3.2 |

**Fig. (17). Application results of Egyxide against the TSSM associated with Brassicaceae and Lamicaceae medicinal plants in Kom Oshim, 2017.**

b) Egyxide application on the TSSM adults

|  | Brassica napus var. pabularia | Brassica oleracea var palmifolia | Mentha spicata | Mentha longifolia |
| --- | --- | --- | --- | --- |
| 2017-04-04 | 16.8 | 11.35 | 3.57 | 3.71 |
| 2017-04-11 | 16.8 | 10.12 | 2.45 | 3.5 |
| 2017-04-18 | 9.64 | 12 | 2.4 | 3.46 |
| 2017-04-25 | 17.5 | 16.49 | 2.64 | 3.54 |
| 2017-05-02 | 20.7 | 26.72 | 2.89 | 3.89 |
| 2017-05-09 | 23.25 | 32.64 | 3 | 4 |
| 2017-05-16 | 15 | 29.51 | 2.69 | 3.75 |
| 2017-05-23 | 14.62 | 28.75 | 2.58 | 3.77 |
| 2017-05-30 | 25.12 | 30.18 | 2.79 | 4 |
| 2017-06-06 | 30.46 | 32.97 | 2.98 | 4.35 |
| 2017-06-13 | 38.1 | 36 | 3.12 | 4.84 |
| 2017-06-20 | 45.12 | 40.44 | 3.33 | 5 |
| 2017-06-27 | 19.02 | 25.1 | 3.86 | 5.23 |
| 2017-07-04 | 20.56 | 26.6 | 4.15 | 5.65 |
| 2017-07-11 | 27.12 | 31.56 | 3.4 | 4.9 |
| 2017-07-18 | 30 | 35.15 | 3.29 | 4.96 |
| 2017-07-25 | 34.85 | 38 | 3.52 | 5.12 |
| 2017-08-01 | 39.5 | 39.09 | 3.7 | 5.2 |

a) Egyxide on T. tabaci

|  | Brassica napus var. pabularia | Brassica oleracea var palmifolia | Mentha spicata | Mentha longifolia |
| --- | --- | --- | --- | --- |
| 2017-04-25 | 1 | 1.49 |  |  |
| 2017-05-02 | 2.05 | 1.79 |  |  |
| 2017-05-09 | 0.9 | 0.99 | 0.6 | 0.78 |
| 2017-05-16 | 0.93 | 1.05 | 0.44 | 0.52 |
| 2017-05-23 | 2.66 | 2.25 | 0.42 | 0.5 |
| 2017-05-30 | 6 | 6.18 | 0.49 | 0.83 |
| 2017-06-06 | 3.15 | 3.54 | 0.79 | 1.1 |
| 2017-06-13 | 3 | 3.29 | 1 | 1.05 |
| 2017-06-20 | 5.9 | 8.44 | 0.68 | 0.85 |
| 2017-06-27 | 10.18 | 12.6 | 0.65 | 0.81 |
| 2017-07-04 | 6.12 | 8.99 |  |  |
| 2017-07-11 | 6.1 | 8.8 |  |  |
| 2017-07-18 | 7 | 13 |  |  |

**Fig. (18 a & b). Application results of Egyxide against *T. tabaci* and *B. tabaci* associated with Brassicaceae and Lamicaceae medicinal plants in Kom Oshim, 2017.**

b) Egyxide on B. tabaci

|  | Brassica napus var. pabularia | Brassica oleracea var palmifolia | Mentha spicata | Mentha longifolia |
| --- | --- | --- | --- | --- |
| 2017-05-02 | 49.01 | 27.44 | 2.15 | 0.22 |
| 2017-05-09 | 35.71 | 13.99 | 1.66 | 0.1 |
| 2017-05-16 | 29.52 | 13.25 | 1.58 | 0.09 |
| 2017-05-23 | 31 | 16.81 | 1.6 | 0.12 |
| 2017-05-30 | 48.16 | 43.12 | 1.79 | 0.23 |
| 2017-06-06 | 29.11 | 29.64 | 2 | 0.11 |
| 2017-06-13 | 27.6 | 27.54 | 1.85 | 0.1 |
| 2017-06-20 | 33.98 | 34.3 | 1.89 | 0.19 |
| 2017-06-27 | 45 | 51.74 | 1.32 | 0.31 |
| 2017-07-04 | 65.81 | 72.31 | 1.76 | 0.69 |
