## Supplementary file 2 for "Predatory mites, a green pesticide, and an Entomopathogenic compound: A proposed IPM tactic based on pest species diversity indices and population dynamics"

**Supplementary 2: figures**

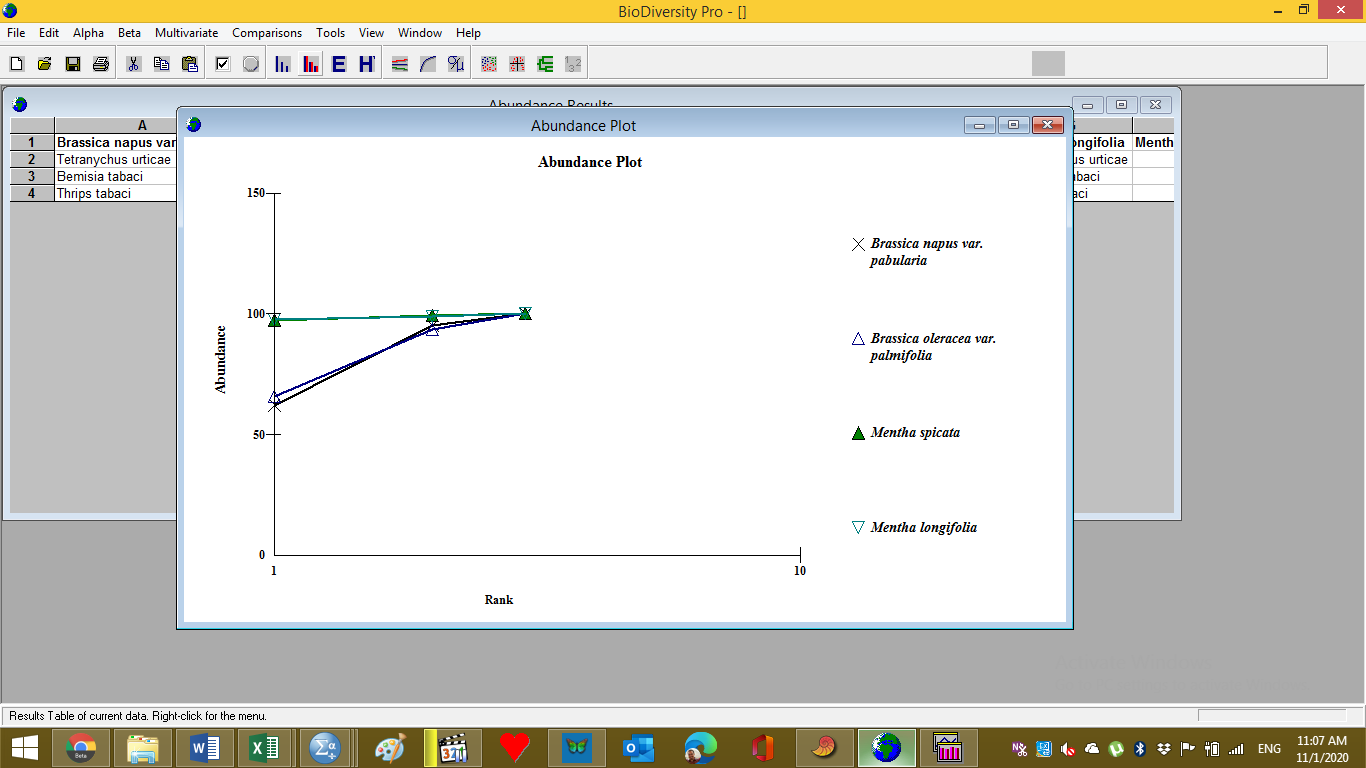

**Suppl. (1). Abundance of pest species populations on *B. napus var. pabularia*, *B. oleracea var. palmifolia* (Brassicaceae) *M. spicata*, and *M. longifolia* (Lamiaceae), in Kom Oshim, 2017.**

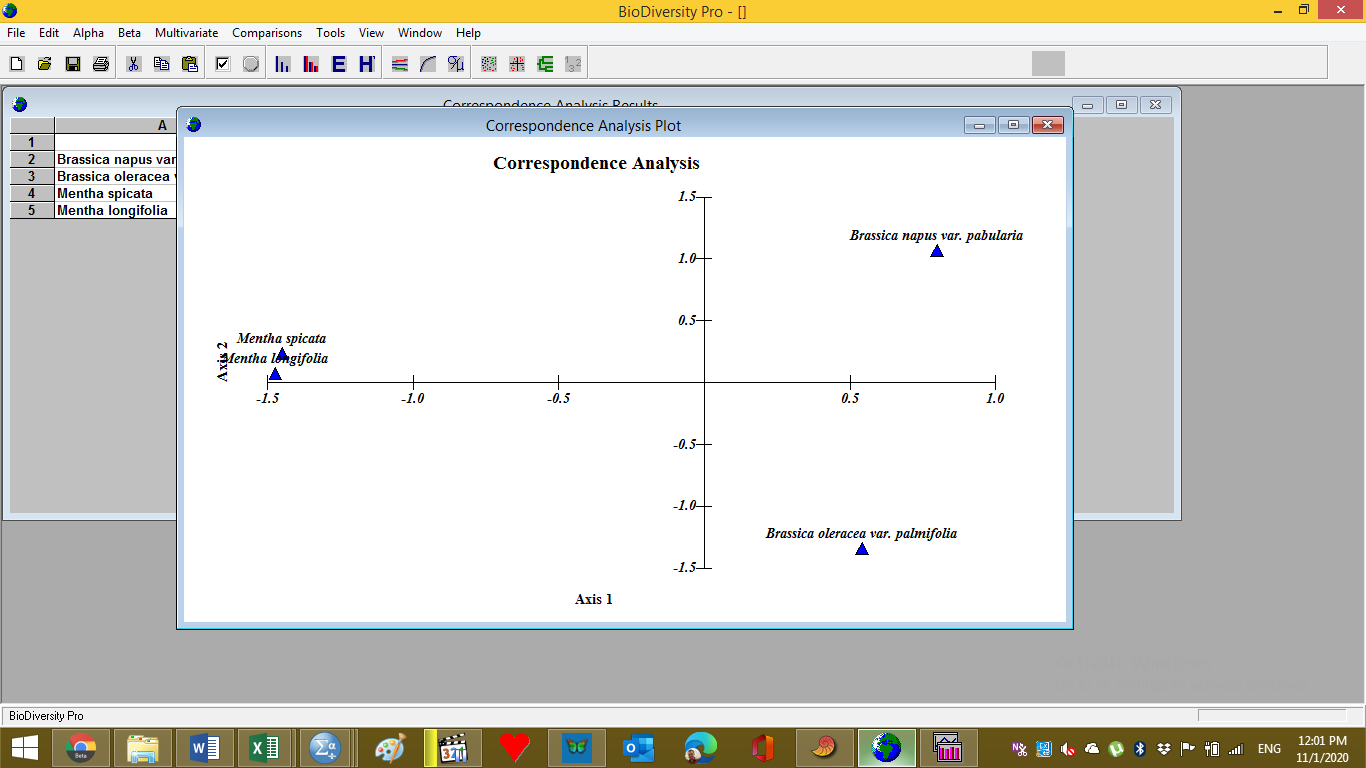

**Suppl. (2). Correspondence analysis plot of pest species populations on *B. napus var. pabularia*, *B. oleracea var. palmifolia* (Brassicaceae) *M. spicata*, and *M. longifolia* (Lamiaceae), in Kom Oshim, 2017.**

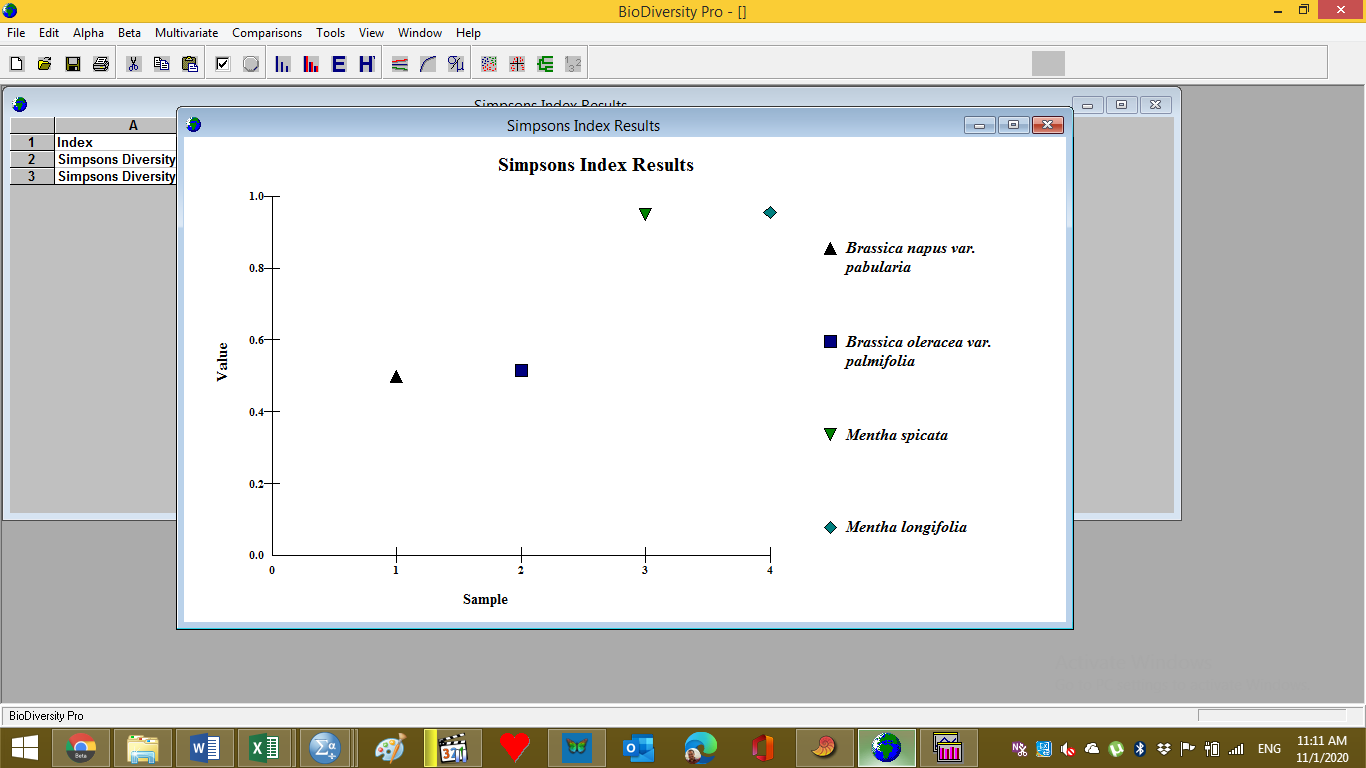

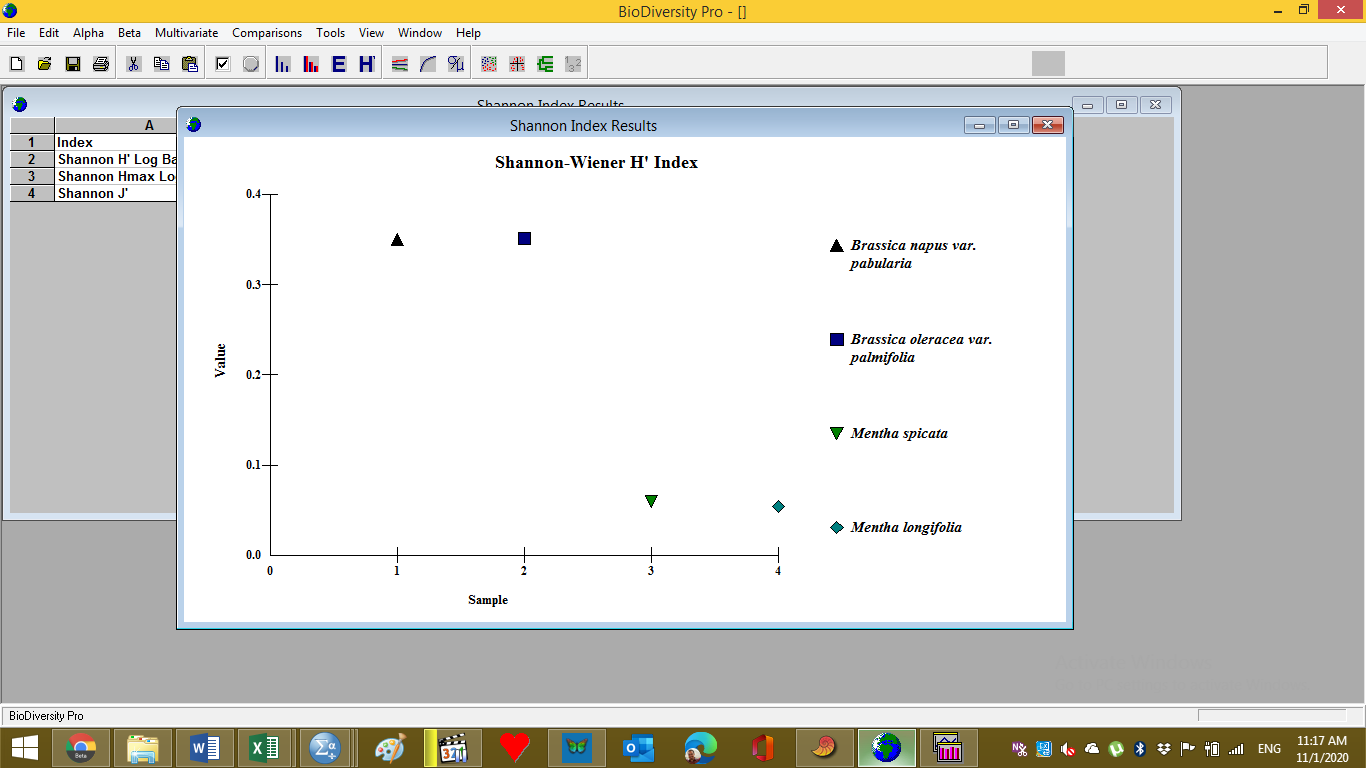

**Suppl. (4). Simpson’s index (D) of pest populations biodiversity on *B. napus var. pabularia*, *B. oleracea var. palmifolia* (Brassicaceae) *M. spicata*, and *M. longifolia* (Lamiaceae), in Kom Oshim, 2016.**

**Suppl. (3). Shanon-Wienner (H') index of populations biodiversity on *B. napus var. pabularia*, *B. oleracea var. palmifolia* (Brassicaceae) *M. spicata*, and *M. longifolia* (Lamiaceae), in Kom Oshim, 2016.**

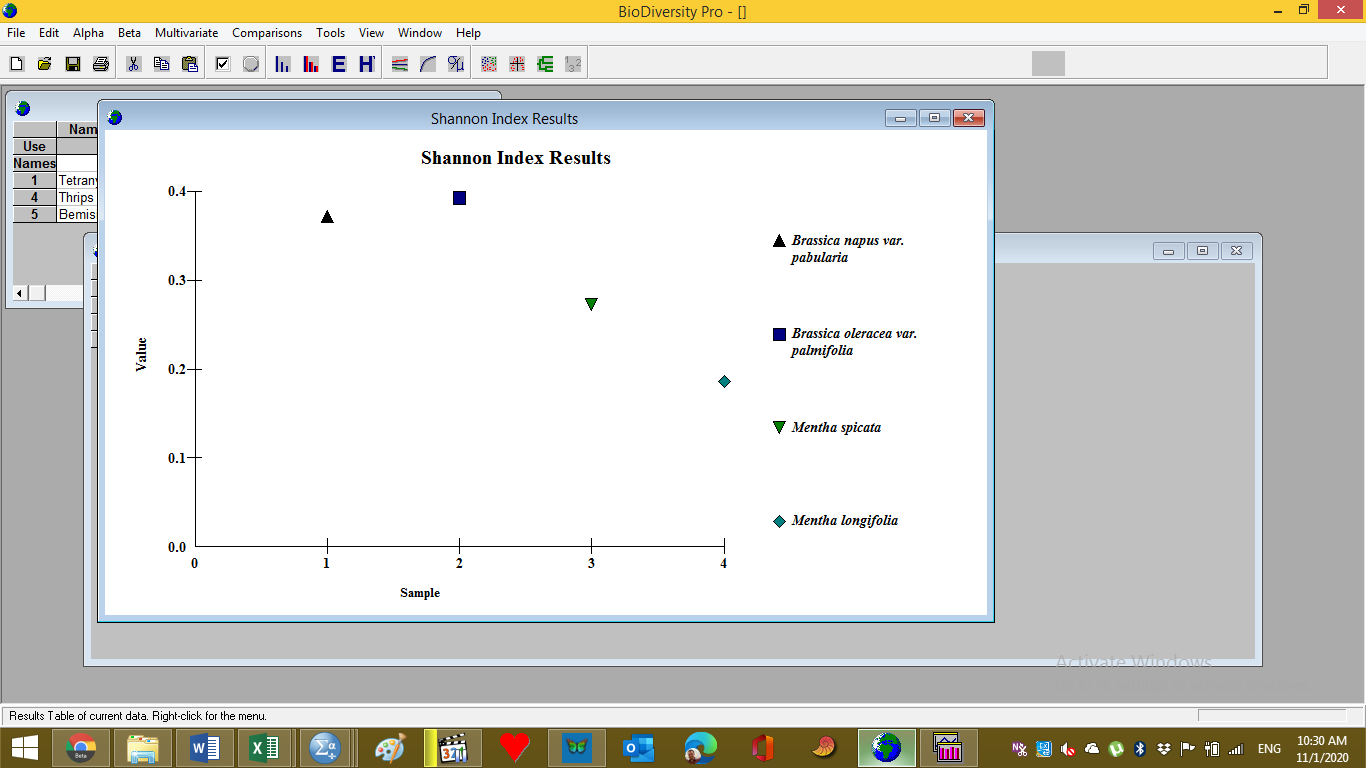

**Suppl. (5). Shanon-Wienner (H') index of populations biodiversity on *B. napus var. pabularia*, *B. oleracea var. palmifolia* (Brassicaceae) *M. spicata*, and *M. longifolia* (Lamiaceae), in Kom Oshim, 2016.**

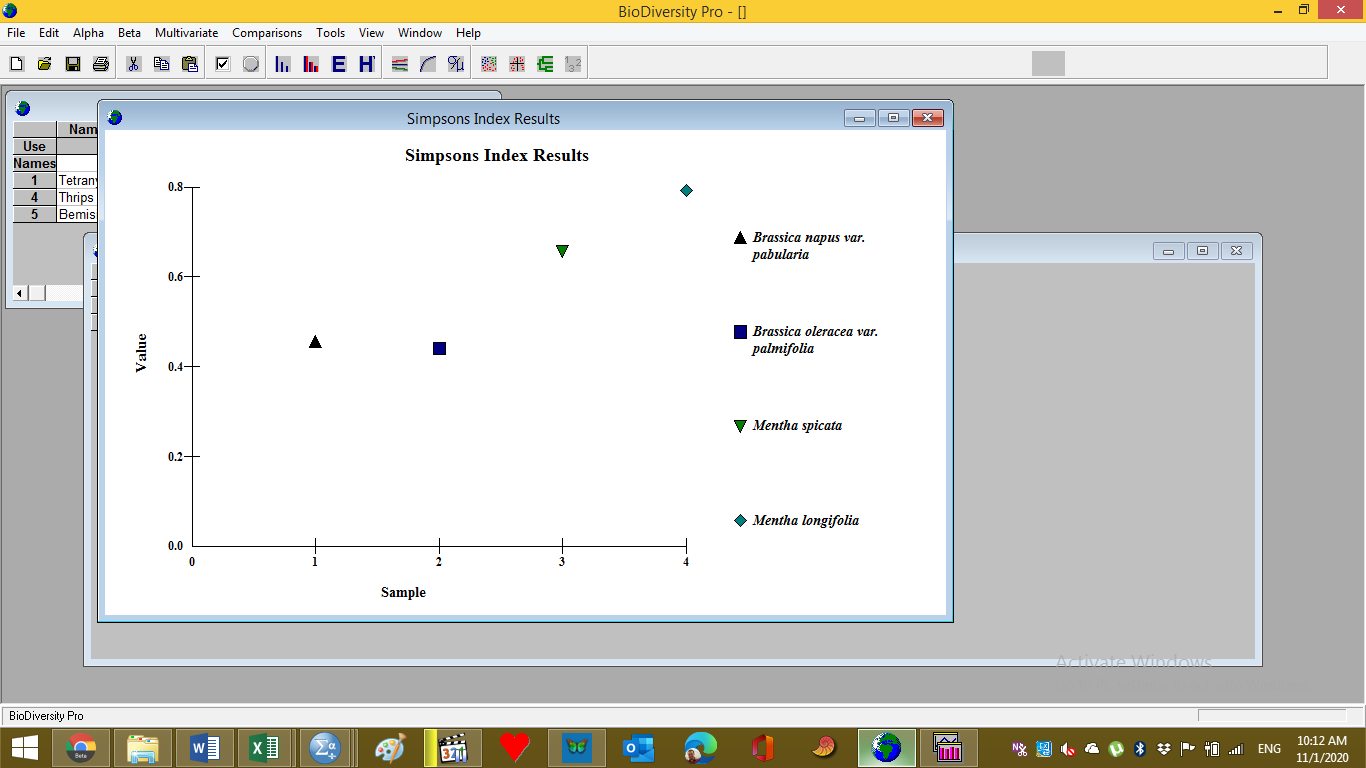

**Suppl. (6). Simpson’s index (D) of pest populations biodiversity on *B. napus var. pabularia*, *B. oleracea var. palmifolia* (Brassicaceae) *M. spicata*, and *M. longifolia* (Lamiaceae), in Kom Oshim, 2016.**

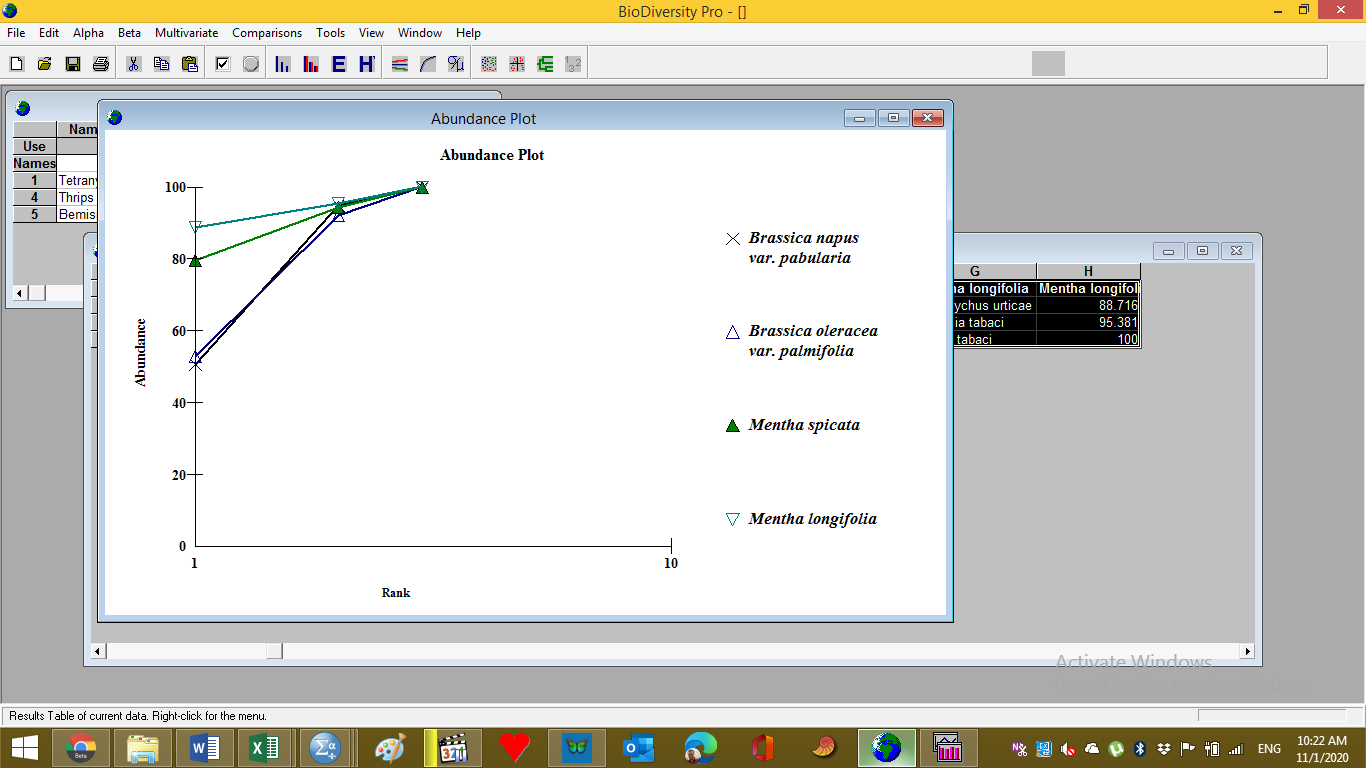

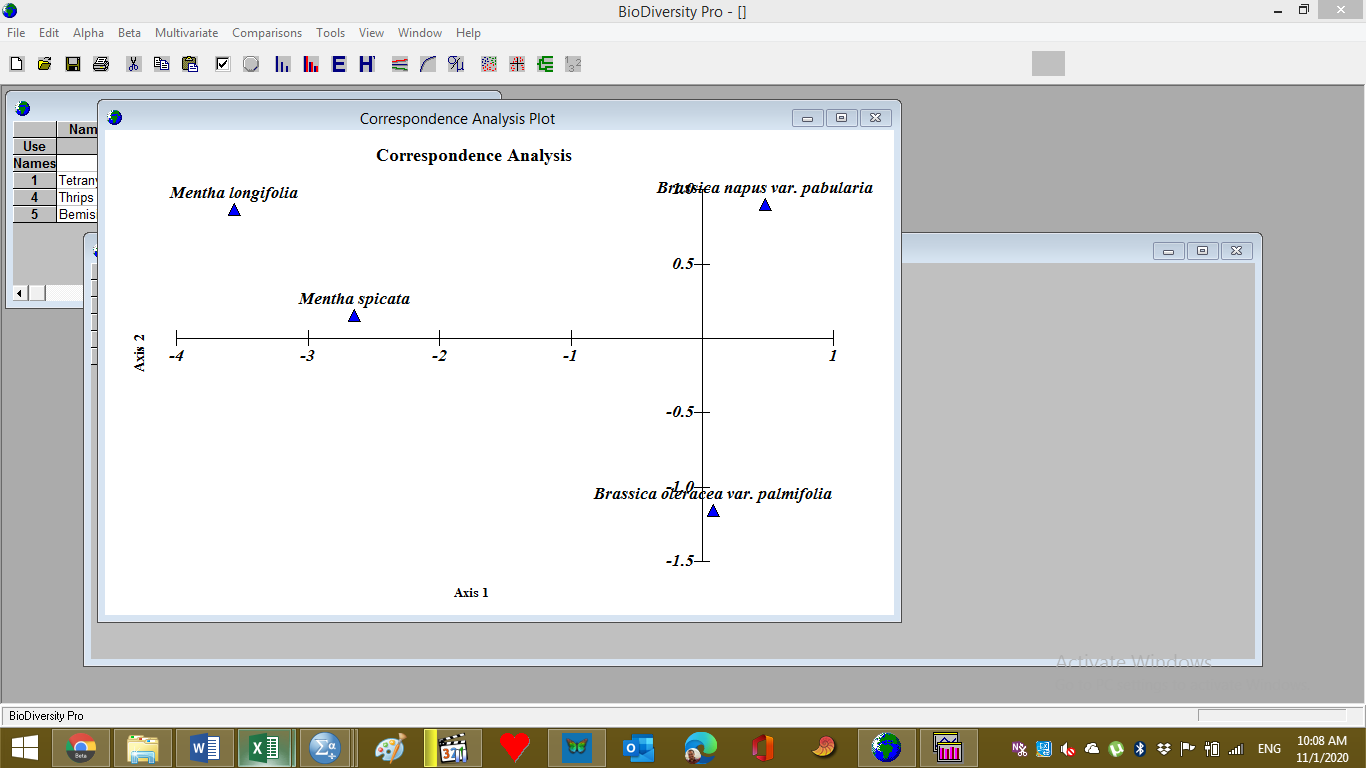

**Suppl. (7). Abundance of pest species populations on *B. napus var. pabularia*, *B. oleracea var. palmifolia* (Brassicaceae) *M. spicata*, and *M. longifolia* (Lamiaceae), in Kom Oshim, 2016.**

**Suppl. (8). Correspondence analysis plot of pest species populations on *B. napus var. pabularia*, *B. oleracea var. palmifolia* (Brassicaceae) *M. spicata*, and *M. longifolia* (Lamiaceae), in Kom Oshim, 2016.**

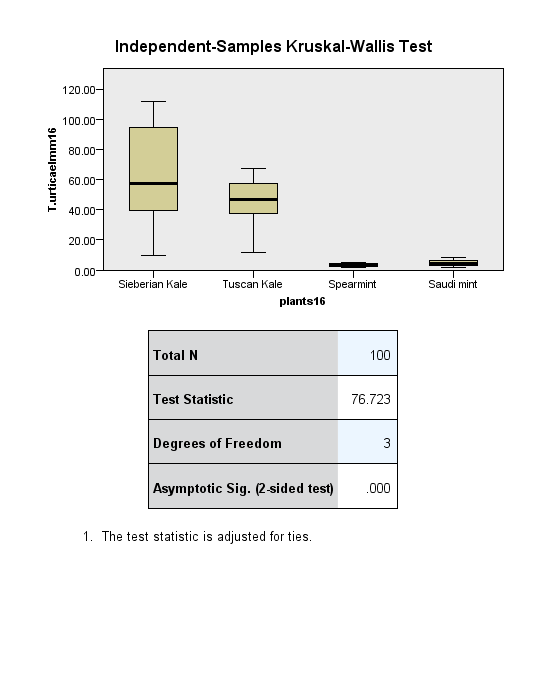

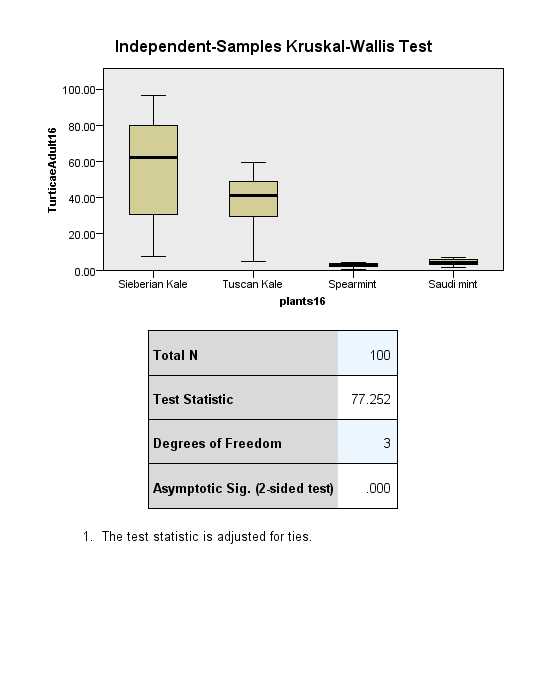

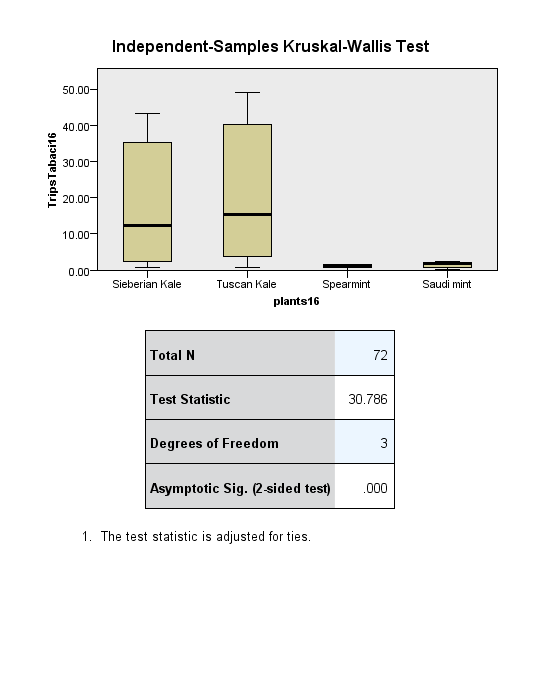

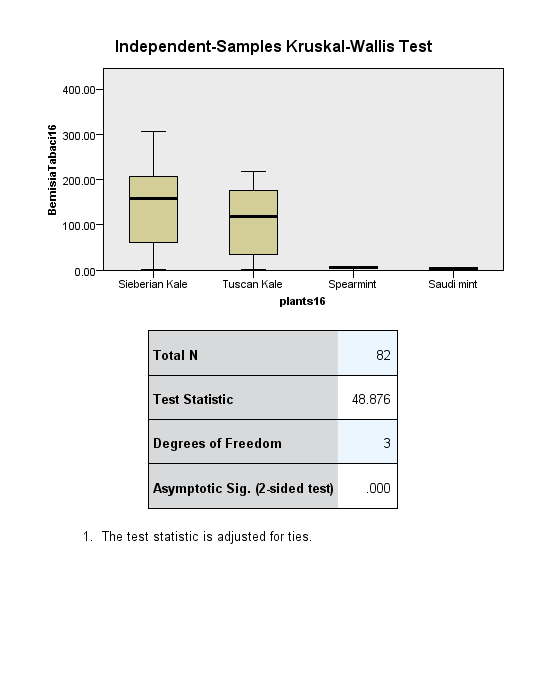

**Suppl. (9).** **Independent samples (Kruskal-Wallis) test, a hypothesis test of *Tetranychus urticae* active stages (immatures and adults), *Thrips tabaci* (active stages), and *Bemisia tabaci* (crawlers and adult female) distribution in Om Saber, El Beheira, 2016 at significant level = 95%.**

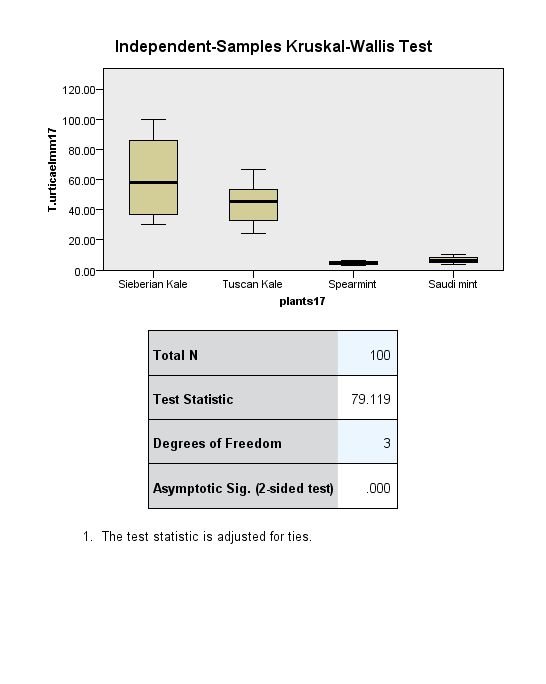

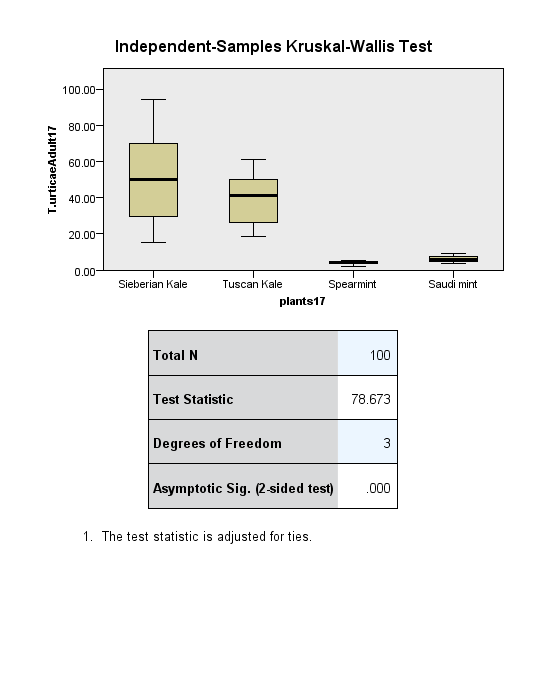

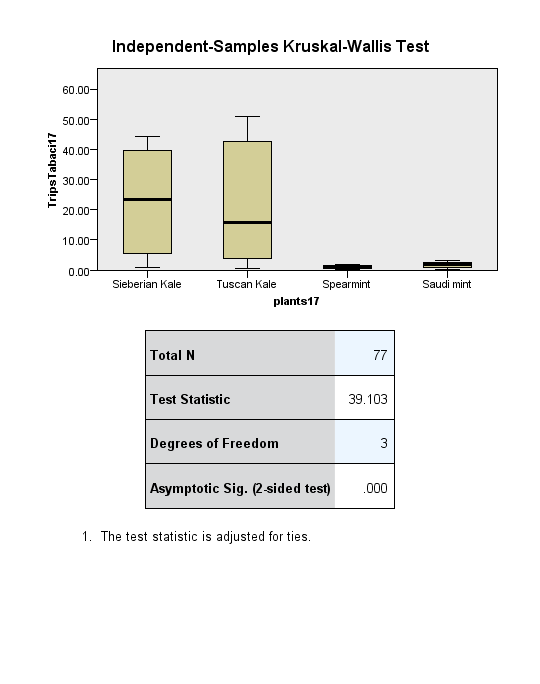

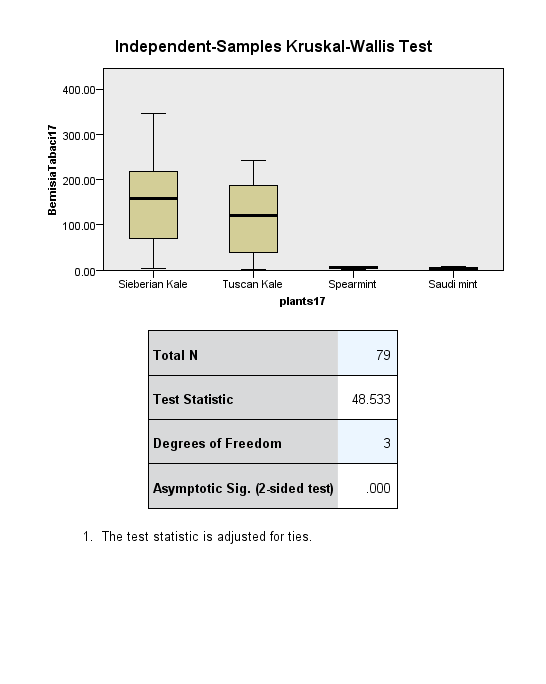

**Suppl. (10).** **Independent samples (Kruskal-Wallis) test, a hypothesis test of *Tetranychus urticae* active stages (immatures and adults), *Thrips tabaci* (active stages), and *Bemisia tabaci* (crawlers and adult female) distribution in Om Saber, 2017 at significant level = 95%.**

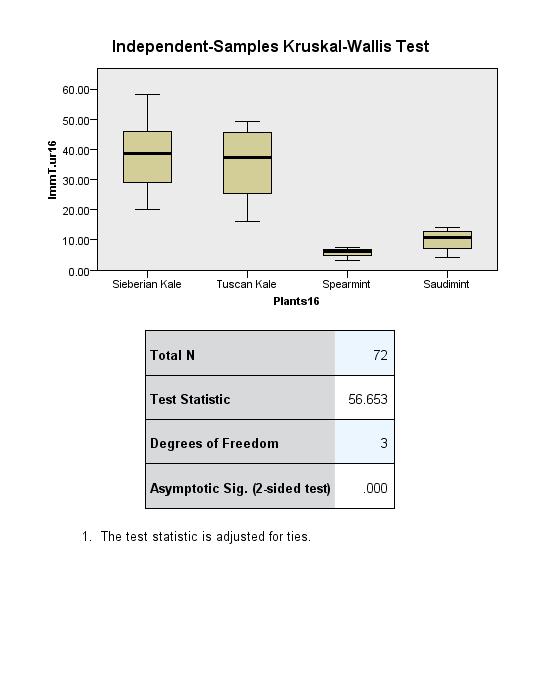

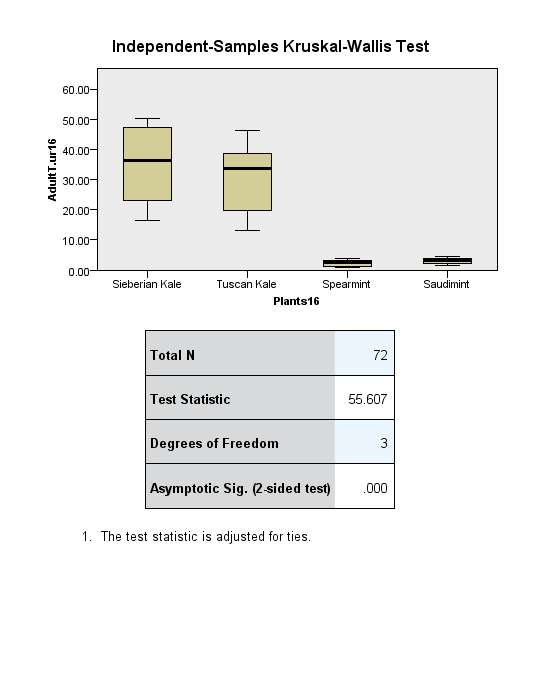

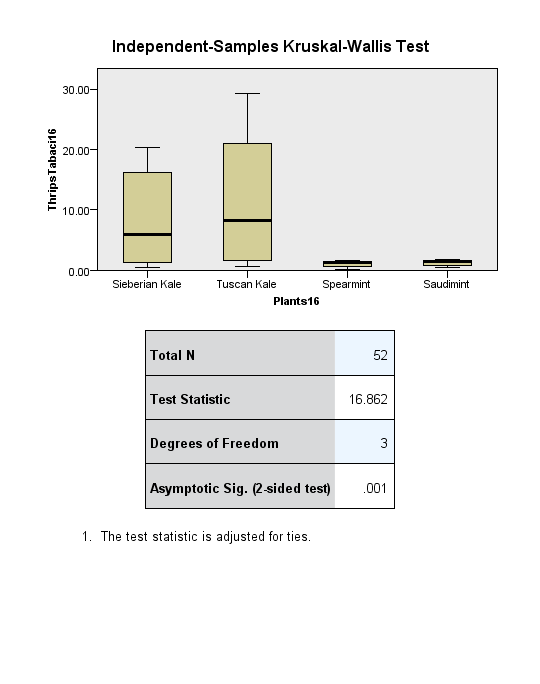

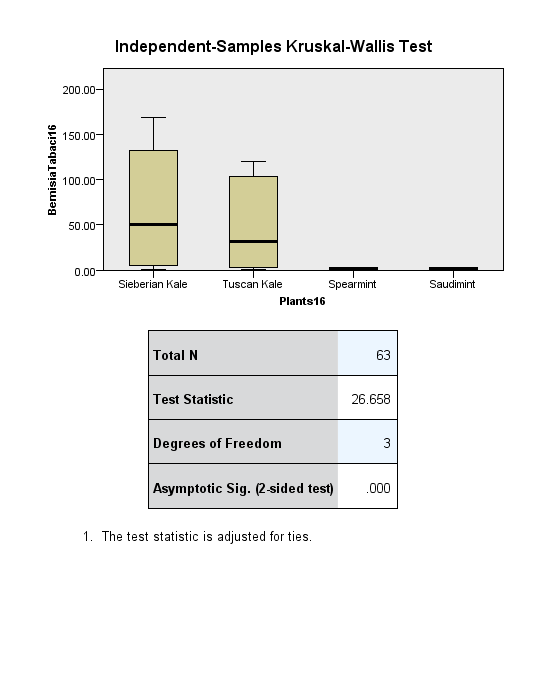

**Suppl. (11).** **Independent samples (Kruskal-Wallis) test, a hypothesis test of *Tetranychus urticae* active stages (immatures and adults), *Thrips tabaci* (active stages), and *Bemisia tabaci* (crawlers and adult female) distribution in Kom Oshim, 2016 at significant level = 95%.**

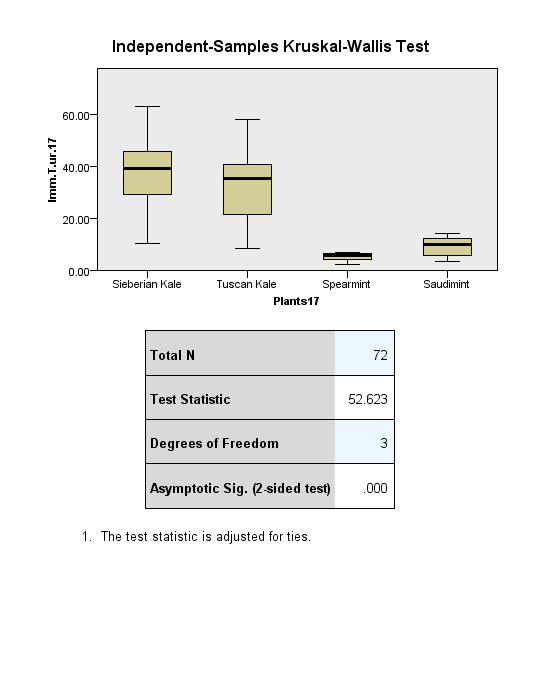

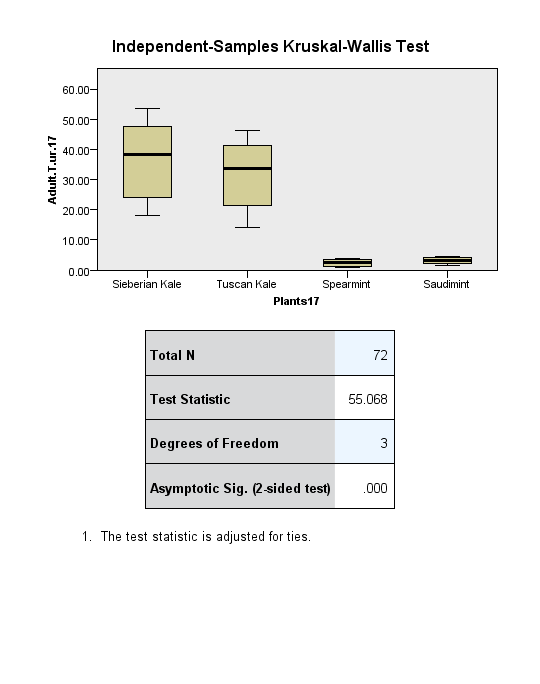

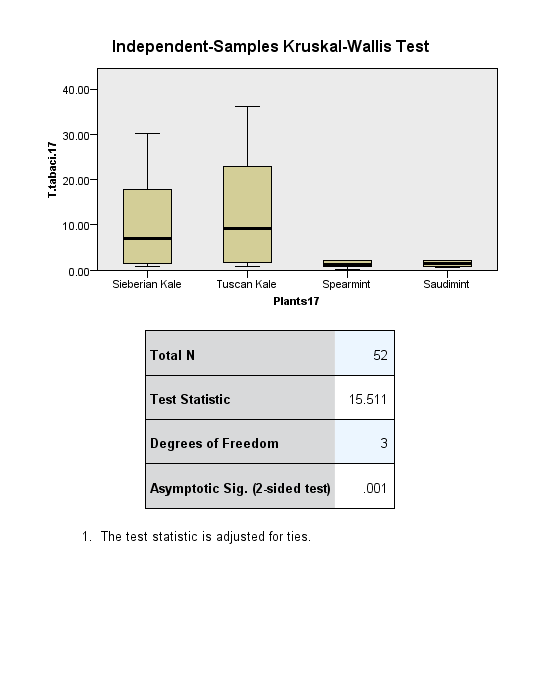

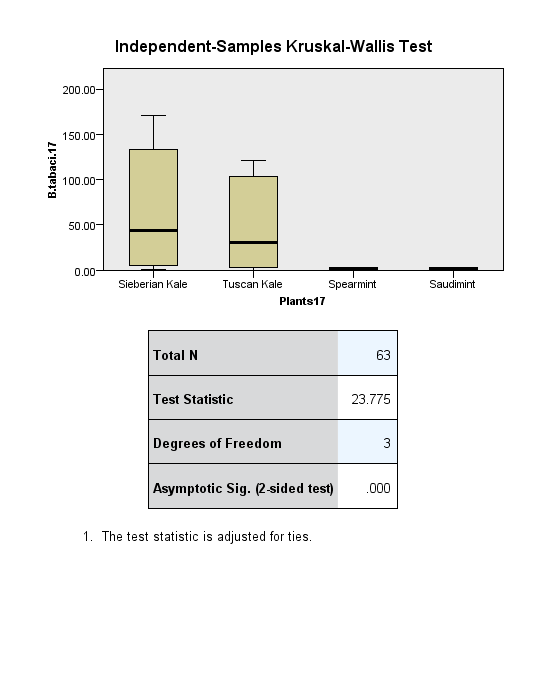

**Suppl. (12).** **Independent samples (Kruskal-Wallis) test, a hypothesis test of *Tetranychus urticae* active stages (immatures and adults), *Thrips tabaci* (active stages), and *Bemisia tabaci* (crawlers and adult female) distribution in Kom Oshim, 2017 at significant level = 95%.**

**Suppl. (13 a). Pest species associated with Brassicaceae plants in Om Saber, 2017, after application of *P. persmilis*, *A. swirskii*, *C. negevi,* Egyxide and Bio-Magic.**

a) Brassica napus var. pabularia

|  | T. urticae nymphs | T. urticae adults | T. tabaci | B. tabaci |
| --- | --- | --- | --- | --- |
| Phytoseiulus persimilis | 4.9 | 7.18 | 19.85 | 112.91 |
| Amblyseius swirskii | 11.65 | 14.34 | 0.82 | 3.32 |
| Cydnoseius negevi | 19.37 | 21.35 | 19.56 | 5.2 |
| Bio-Magic | 20.27 | 23.11 | 5.96 | 19.85 |
| Egyxide | 20.34 | 23.73 | 9.6 | 16.58 |
| Control | 71.31 | 65 | 19.7 | 113.68 |

b) Brassica oleraceae var palmifolia

|  | T. urticae nymphs | T. urticae adults | T. tabaci | B. tabaci |
| --- | --- | --- | --- | --- |
| Phytoseiulus persimilis | 7.48 | 7.94 | 24.08 | 114.64 |
| Amblyseius swirskii | 8.09 | 16.66 | 0.96 | 2.38 |
| Cydnoseius negevi | 8.93 | 19.7 | 22.91 | 4.8 |
| Bio-Magic | 26.28 | 22 | 5.96 | 27.86 |
| Egyxide | 20.42 | 22.92 | 9.6 | 38.27 |
| Control | 50.19 | 43.55 | 23.65 | 114.74 |

**Fig. (13 b). Pest species associated with Lamicaceae plants in Om Saber, 2017**

**Suppl. (14 a). Pest species associated with Brassicaceae plants in Kom Oshim, 2017, after application of *P. persmilis*, *A. swirskii*, *C. negevi,* Egyxide and Bio-Magic**

c) Mentha spicata

|  | T. urticae nymphs | T. urticae adults | T. tabaci | B. tabaci |
| --- | --- | --- | --- | --- |
| Phytoseiulus persimilis | 0.289524 | 0.393333 | 1.09357 | 5.48286 |
| Amblyseius swirskii | 0.460952 | 0.677619 | 0.0364286 | 0.945 |
| Cydnoseius negevi | 0.511905 | 1.51619 | 1.05286 | 0.871429 |
| Bio-Magic | 1.43381 | 0.968095 | 0.524286 | 2.52714 |
| Egyxide | 1.28762 | 0.94619 | 0.685714 | 3.22714 |
| Control | 3.73667 | 3.06286 | 1.08286 | 5.27214 |

d) Mentha longifolia

|  | T. urticae nymphs | T. urticae adults | T. tabaci | B. tabaci |
| --- | --- | --- | --- | --- |
| Phytoseiulus persimilis | 0.324286 | 0.41 | 1.62929 | 4.74571 |
| Amblyseius swirskii | 0.575714 | 1.20048 | 0.01 | 0.624286 |
| Cydnoseius negevi | 0.913333 | 2.06 | 1.595 | 1.14071 |
| Bio-Magic | 2.09857 | 2.44238 | 0.005 | 1.66429 |
| Egyxide | 2.27238 | 1.97095 | 0.604286 | 1.96143 |
| Control | 5.32905 | 4.68857 | 1.61714 | 3.875 |

a) Brassica napus var. pabularia

|  | TSSM nymph | TSSM adult | Thrips tabaci | Bemisia tabaci |
| --- | --- | --- | --- | --- |
| Phytoseiulus persimilis | 6.855 | 7.723 | 10.6669 | 125.127 |
| Amblyseius swirskii | 8.925 | 8.95222 | 0.548333 | 20.7011 |
| Cydnoseius negevi | 11.2633 | 13.0794 | 10.4323 | 21.375 |
| Bio-Magic | 26.6044 | 19.7844 | 3.84846 | 35.107 |
| Egyxide | 24.4861 | 24.6756 | 4.23 | 39.49 |
| Control | 37.8856 | 35.1722 | 9.88308 | 122.435 |

b) Brassica oleracea var palmifolia

|  | TSSM nymph | TSSM adult | Thrips tabaci | Bemisia tabaci |
| --- | --- | --- | --- | --- |
| Phytoseiulus persimilis | 7.915 | 6.824 | 13.7808 | 85.897 |
| Amblyseius swirskii | 10.8389 | 18.3056 | 0.678333 | 11.7588 |
| Cydnoseius negevi | 18.0894 | 26.055 | 13.6177 | 12.585 |
| Bio-Magic | 18.045 | 22.2744 | 5.80769 | 32.901 |
| Egyxide | 20.8878 | 27.9261 | 5.57 | 33.014 |
| Control | 84.955 | 57.6222 | 13.3962 | 84.443 |

**Suppl. (14 b). Pest species associated with Lamiaceae plants in Kom Oshim, 2017**

c) Mentha spicata

|  | T. urticae nymph | T. urticae adult | Thrips tabaci | Bemisia tabaci |
| --- | --- | --- | --- | --- |
| Phytoseiulus persimilis | 1.01333 | 1.42917 | 1.71111 | 4.172 |
| Amblyseius swirskii | 1.12778 | 2.32214 | 0.2475 | 0.541 |
| Cydnoseius negevi | 0.955833 | 2.15389 | 1.30125 | 0.571667 |
| Bio-Magic | 1.22444 | 3.545 | 0.5875 | 1.77944 |
| Egyxide | 1.055 | 3.13111 | 0.63375 | 1.79833 |
| Control | 3.77056 | 5.75444 | 1.26625 | 2.417 |

d) Mentha longifolia

|  | T. urticae nymph | T. urticae adult | Thrips tabaci | Bemisia tabaci |
| --- | --- | --- | --- | --- |
| Phytoseiulus persimilis | 0.64 | 0.983 | 1.75 | 3.69 |
| Amblyseius swirskii | 0.766667 | 1.41667 | 0.318333 | 0.105 |
| Cydnoseius negevi | 1.63056 | 2.30556 | 1.40625 | 0.111 |
| Bio-Magic | 3.45889 | 2.45889 | 0.7425 | 0.219 |
| Egyxide | 3.15056 | 2.15056 | 0.805 | 0.216 |
| Control | 5.03444 | 3.43444 | 1.42625 | 3.92 |
